# Open-State Dynamics and Allosteric Modulation of the α1β3γ2 GABA_A_ Receptor Stabilized by L9′T/S Substitutions

**DOI:** 10.64898/2026.02.02.703268

**Authors:** Ayobami Diyaolu, Cecilia M Borghese, Marcel P Goldschen-Ohm, Senthil Natesan

## Abstract

GABA type A receptors (GABA_A_Rs) mediate inhibitory neurotransmission, and their dysfunction contributes to epilepsy, anxiety, and depression. Although closed and desensitized structures of heteropentameric GABA_A_Rs are known, an open-state conformation has been difficult to capture. Here we use in-silico mutagenesis and Gaussian-accelerated MD to stabilize open-like ensembles of the α1β3γ2 receptor via hydrophilic substitutions at the hydrophobic 9′ gate (L9′T/L9′S). The mutants expand the pore at 9′ and 20′ (extracellular pore entry), increase hydration and water flux, and lower Cl⁻ permeation barriers at 9′ and -2′ (desensitization gate) from ∼19/∼9 kcal mol⁻¹ (WT) to ∼1.3-1.6/∼3.0-3.4 kcal mol^−1^, yielding ohmic conductance ∼10-30 pS in computational electrophysiology. Conformationally, the mutants show reduced twist, outward M2 tilts, and C-loop closure, consistent with activation-like signatures. On this open-like background, PAMs (diazepam, ganaxolone) primarily tune the residual -2′ constriction, whereas bicuculline (orthosteric antagonist) drives a time-ordered, bottom-to-top closing sequence (-2′ first, then 9′/20′, then twist) via an asymmetric collapse of the M2 bundle that approaches closed-state landmarks while sampling desensitized-like ECD-TMD coupling. Two-electrode voltage clamp confirms spontaneous activity in L9′T-containing receptors, supporting the mutant-stabilized open-like ensembles. These results provide a coherent atomistic framework for gating and allosteric modulation in α1β3γ2 GABA_A_Rs and establish a tractable, ligand-responsive platform for state-selective structure-based design.

## INTRODUCTION

Pentameric ligand-gated ion channels (pLGICs), including γ-amino butyric acid type A (GABA_A_) and glycine receptors, mediate fast synaptic neurotransmission^1–3^. Each receptor comprises five subunits with an extracellular domain (ECD) harboring orthosteric agonist sites and a transmembrane domain (TMD) containing four helices (M1-M4), with M2 forming the ion-conducting pore^4,5^. GABA_A_ receptors (GABA_A_Rs) are central to inhibitory signaling in the brain and are implicated in disorders such as anxiety, autism, depression, essential tremor, and epilepsy^6–9^. They are also targets of numerous FDA-approved drugs, including benzodiazepines, z-drugs, neurosteroids, and anesthetics^10,11^. Native GABA_A_Rs are typically heteropentameric, most commonly α1β2γ2 and α1β3γ2 subtypes, in which two α, two β, and one γ subunit assemble into a pseudosymmetric pentamer^12–14^.

Upon agonist binding, GABA_A_Rs transition from a closed state to an open, chloride-conducting state, and with prolonged exposure, to desensitized states^4,15^. A dual-gate framework is often used to describe gating in which the hydrophobic gate at the 9′ position and the desensitization gate near the −2′ position modulate permeation^16–18^. Nonetheless, electrophysiological studies indicate a richer landscape with multiple intermediates and desensitized states^19–21^.

Recent cryo-electron microscopy (cryo-EM) structures have captured closed, open, and desensitized conformations across several homomeric pLGICs, clarifying conserved activation motifs such as ECD untwisting with β-sandwich contraction and outward tilting of the M2 helices in the TMD^18,22–27^. However, an open-state structure for heteropentameric GABA_A_Rs remains elusive, likely due to rapid desensitization and pseudosymmetry^28,29^. This gap limits mechanistic understanding of gating transitions and the structural basis of allosteric modulation, complicating rational design of state-selective ligands.

Hydrophilic substitutions at the 9′ position (e.g., L9′T and L9′S) destabilize the hydrophobic gate and produce spontaneously active channels across multiple subunit combinations^30–32^. These mutations increase GABA sensitivity (Table S1) and spontaneous activity and have been widely used to probe allosteric modulation by agents, such as bicuculline, picrotoxin, diazepam, and neurosteroids^33–35^. Yet, the atomistic mechanisms by which these substitutions reshape receptor conformations and gating transitions remain incompletely defined.

Here, we combine in silico mutagenesis with enhanced molecular dynamics, specifically Gaussian-accelerated MD (GaMD), steered MD (SMD), umbrella sampling (US), and computational electrophysiology (CompEL) to characterize the structural and functional consequences of L9′T and L9′S substitutions in the closed state (*apo*) α1β3γ2 GABA_A_Rs. Using the high-resolution cryo-EM structure of the α1β3γ2 receptor (PDB 7QNE) as the starting model, we substituted the leucine at the 9′ position of each M2 helix with either threonine or serine (L9′T/S), quantified changes in pore geometry and hydration, delineated energetics of chloride permeation, and evaluated voltage-dependent conductance and ion selectivity. We further examined how positive (PAMs; diazepam, ganaxolone) and negative allosteric modulators (NAMs; bicuculline, picrotoxin) influence these open-like ensembles. Two-electrode voltage-clamp recordings in *Xenopus oocytes* expressing wild-type and mutant α1β3γ2 GABA_A_Rs corroborate the gain-of-function phenotype conferred by L9′T substitutions, supporting the computationally derived open-state ensembles. Together, these complementary strategies establish a tractable model for the open-like state in a heteropentameric GABA_A_R and clarify how allosteric ligands bias the gating landscape.

## RESULTS

To elucidate the structural and functional consequences of hydrophilic L9′T and L9′S substitutions in the α1β3γ2 GABA_A_R, we combined electrophysiological and computational approaches. Two-electrode voltage-clamp recordings of wild-type and mutant α1β3γ2 GABA_A_Rs in *Xenopus laevis* oocytes validated the gain-of-function phenotype conferred by the L9′T substitution. We then performed in-silico mutagenesis and enhanced sampling molecular dynamics simulations to characterize mutation-induced changes in pore geometry, hydration, ion permeation energetics, and conformational dynamics in atomistic detail. We further examined how positive and negative allosteric modulators influence these open-state ensembles. Together, these complementary strategies provide a detailed framework for understanding gating transitions and allosteric modulation in the open-state.

### Two-Electrode Voltage Clamp Electrophysiology

To experimentally validate the gain-of-function phenotype, we expressed wild-type and L9′T mutant α1β3γ2 GABA_A_ receptors in *Xenopus laevis* oocytes and carried out two-electrode voltage clamp recordings. Specifically, we assessed channel activity in response to GABA or the pore-blocker picrotoxin (PTX), focusing on the effects of L9′T substitutions in individual or all subunits.

Representative traces from the recordings are shown in **Figure S1**.

Incorporation of the gain-of-function L9′T substitution in either the α1 or γ2 subunit enhanced sensitivity to GABA, as evidenced by left-ward shifts in the GABA concentration-response curve (CRC) (**Figure 1A**). Channels containing the L9′T substitution in the β3 subunit (both the single and triple mutant) exhibited small current amplitudes and pronounced rundown, complicating CRC acquisition.

**Figure 1.**
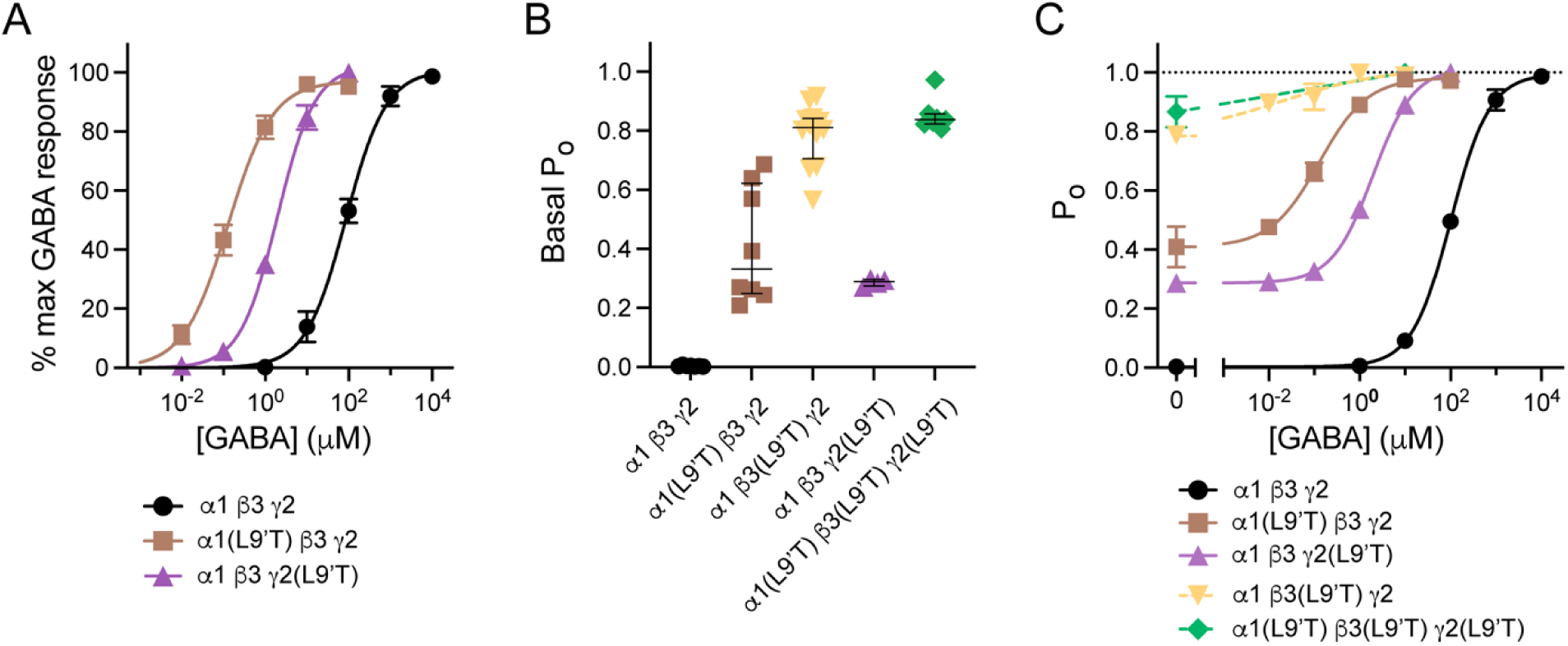
Experimental measures of GABA sensitivity and unliganded open probability for L9′T substitutions in individual or all subunits. **(A)** GABA concentration-response curves (CRCs) normalized to the maximal current amplitude elicited by saturating GABA (*I*_GABA_, as shown in Figure S1). Data points are mean ± SEM across oocytes, and curves are global fits with the Hill equation (Eq. 1; α1β3γ2: *EC*_50_ = 88 mM, *n*_H_ = 0.90, *n* = 7; α1(L9′T) β3γ2: *EC*_50_ = 0.13 mM, *n*_H_ = 0.81, *n* = 8; α1β3γ2(L9′T): *EC*_50_ = 2.0 mM, *n*_H_ = 0.96, *n* = 4; where *n* is the number of oocytes). Low expression and/or rundown prevented us from obtaining reliable CRCs for α1β3(L9′T)γ2 or α1(L9′T)β3(L9′T)γ2(L9′T) mutants, although we were able to observe their response to both PTX and saturating GABA (see panels B-C). **(B)** Basal unliganded open probability (*P*_0_) estimated from the ratio of PTX-sensitive to total current (*I*_PTX_/*I*_total_) as illustrated in Figure S1. Data points are from individual oocytes, and bars are median ± quartiles. Number of oocytes in each group: 4-12. **(C)** GABA CRCs, as described in panel A, expressed as *P*_0_ versus GABA concentration by setting the minimal *P*_0_ to the mean estimate for the basal unliganded *P*_0_, as shown in panel B. For β3(L9′T)-containing receptors (dashed lines), small currents and rundown made the determination of the CRCs difficult. This is exacerbated by their high unliganded *P*_0_, which leaves little room for additional activation by GABA. Number of oocytes per group: α1β3γ2: *n* = 7; α1(L9′T) β3γ2: *n* = 8; α1β3(L9′T)γ2: *n* = 12 (2 for some concentrations); α1β3γ2(L9′T): *n* = 4; α1(L9′T)β3(L9′T)γ2(L9′T): *n* = 3.

Nevertheless, recordings suggest that β3(L9′T)-containing receptors are even more sensitive to GABA (i.e., the GABA CRC is further left-shifted) than those with L9′T substitutions in either α1 or γ2 subunits (**Figure 1C**).

To estimate the basal unliganded open probability (*P*_*o*_) typically conferred by L9′T substitutions in α1β3γ2 receptors, we applied PTX to block any spontaneous unliganded standing current and compared the magnitude of the PTX-sensitive current to the total current (current elicited by saturating GABA plus the current blocked by PTX; Figure S1)^36,37^. Under the assumption that channels are fully activated (*P*_0_∼1) by saturating GABA, the unliganded *P*_0_ is given by the ratio of PTX-sensitive (*I*_*PTX*_) to total (*I*_*Total*_) current amplitudes (**Figures 1B and S1**). Given that the L9′T substitution has previously been shown to stabilize the open conformation in α1β3γ2 receptors, this assumption is likely to be reasonable^38^. Using this approach, we estimate that the L9′T substitution in either the α1 or γ2 subunit confers an unliganded *P*_0_ of ∼0.3-0.4, similar to previous reports in α1β2γ2 receptors^36,37^, whereas the single and triple mutant containing the L9′T substitution in the β3 subunit are highly open at rest with an unliganded *P*_0_ of ∼0.8. This high basal activity is consistent with β3(L9′T)-containing receptors being highly sensitized to activation.

We next turned to atomistic simulations to characterize how L9′T and L9′S substitutions remodel the pore, stabilize hydrated conductive states, reshape the energy landscape for chloride permeation, and bias receptor conformation under positive and negative allosteric modulation.

### In-Silico Studies

To investigate the structural and functional consequences of hydrophilic L9′T and L9′S mutations, we used the high-resolution cryo-EM structure of the α1β3γ2 GABA_A_R in complex with the benzodiazepine antagonist, Ro1504513 (PDB ID: 7QNE, resolution = 2.7Å) as the starting model^13^. Ligands were removed, and site-directed mutagenesis introduced L9′T and L9′S substitutions at the 9′ position of the M2 helix in all five subunits (all-subunit mutants) (**Figure 2**). We then carried out Gaussian-accelerated MD (GaMD) simulations (0.5 µs per system, triplicates) to quantify mutation-induced changes in pore geometry and hydration, followed by steered MD (SMD) and umbrella sampling (US) to compute the potential of mean force (PMF) for chloride permeation. Computational electrophysiology (CompEL) was used to estimate voltage-dependent conductance and ion selectivity. Additional receptor-ligand simulations (diazepam, ganaxolone, bicuculline, and picrotoxin) probed allosteric effects on the mutant-stabilized open-like ensembles.

**Figure 2.**
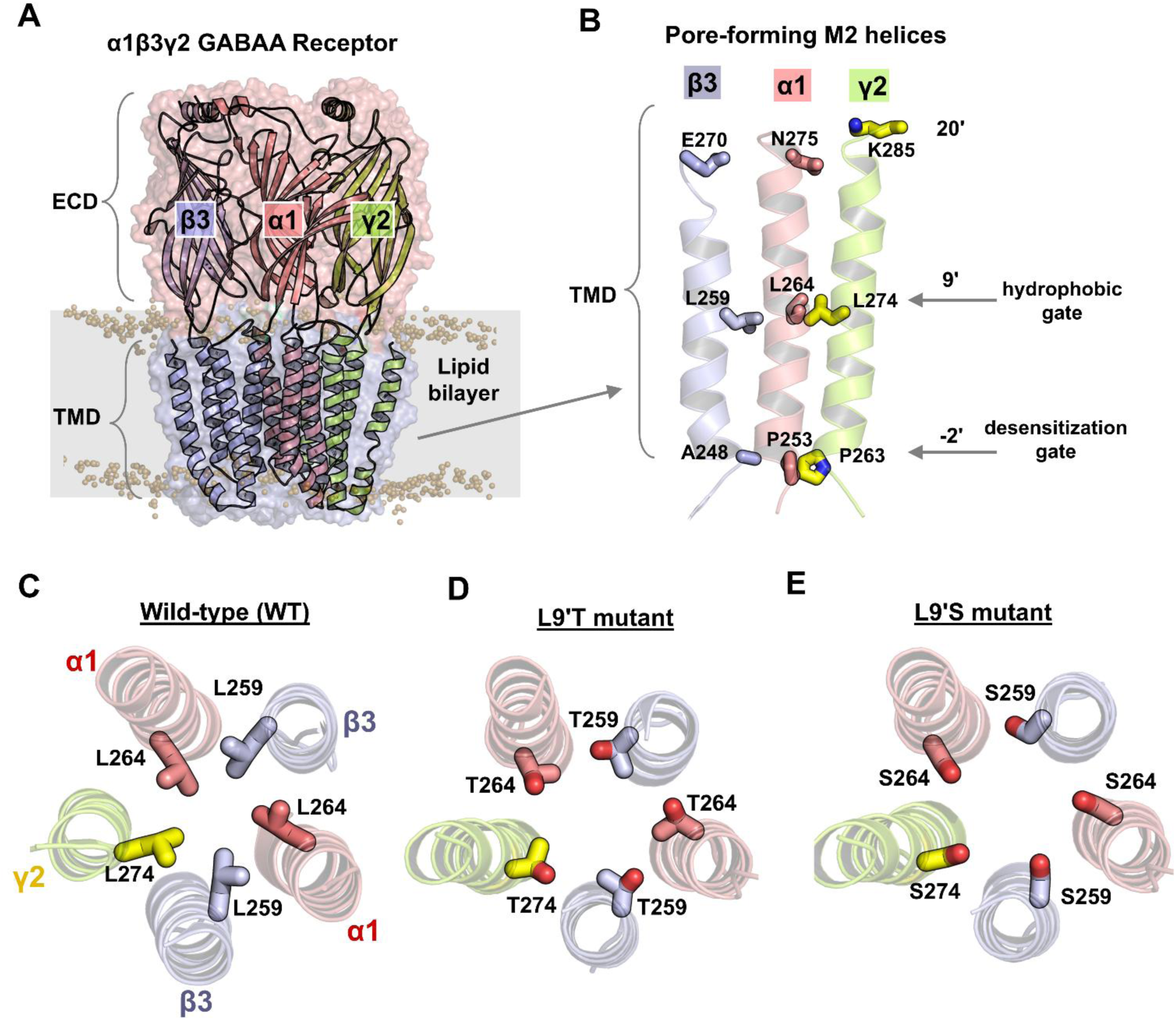
Structural architecture of the α_1_β_3_γ_2_ GABA_A_ receptor and key pore-lining residues. (**A**) Side view of the α1β3γ2 GABA_A_ receptor embedded in a lipid bilayer (gray), showing three of the five subunits. The extracellular (ECD) and transmembrane (TMD) domains are depicted in secondary structure representation. (**B**) Side membrane view of the transmembrane domain, highlighting three of the five pore-forming M2 helices. Key gating residues, 20’ (pore entry), 9’ (hydrophobic gate), and -2’ (desensitization gate) are shown in licorice representation for β3 (light blue), α1 (salmon), and γ2 (lemon) subunits. (**C-E**) Top-down views of the M2 helices forming the channel pore in wild-type (C), L9′T mutant (D), and L9′S mutant systems. Mutations were introduced only at the 9’ position in all five subunits and are shown in licorice representation.

This multi-pronged, comprehensive approach provided high-resolution atomistic insights into gating transitions and the structural basis of open-state stabilization (Box 1).

##### Box 1. Quantitative structural and functional observables compared between WT and mutant α1β3γ2 GABAA receptors

| Category | Observable(s) | How it is computed/interpreted | Where shown |
| --- | --- | --- | --- |
| Pore geometry | Pore radius surfaces (qualitative, HOLE) | HOLE-derived radius surfaces along the pore axis to visualize constrictions/widening at 20', 9', -2' positions. | Fig. 3A-C |
|  | Time-averaged pore radius (Å); Minimum pore radius per frame (Å) | HOLE radius profiles along the pore axis; minimum radius per frame used to track the tightest gate over time. | Fig. 3D-E; S3-S3 |
|  | Pore expansion frequency (PEF, % frames) | Fraction of trajectory frames with minimum radius >2.0 Å (conductive criterion) |  |
|  | Cross-sectional pore area (Å<sup>2</sup>) | Regular pentagon area from M2 Ca-Ca distances at axial positions (20', 9', -2' in main text and 20', 9', 6', 2', -2' in SI); larger area implies backbone widening. | Fig. 3G; S4-S5; S14-S15 |
| Hydration status | Water count near 9' | Number of water molecules within $\pm 7$ Å of 9' gate along the pore axis (7 Å above and 7 Å below (Wordom/MDAnalysis). Higher counts indicate wetted gate. | Fig. 4A-B; S17 |
|  | Water flux (events per 10 ps); Cumulative water flux | Direction-independent count of full transits between 20' and -2' (Wordom). Summed over time for cumulative flux. | Fig. 4C-D; S18-S20 |
| Cl<sup>-</sup> transport energetics (PMF) | Potential of mean force (PMF) long pore axis (kcal/mol); Barrier heights at 9', -2' | SMD → umbrella sampling windows (1 Å spacing, 20 ns/window) → WHAM. Lower barriers reflect facilitated permeation | Fig. 5A-B |
| Computational electrophysiology (CompEL) | Chloride permeation events; anion selectivity | Double bilayer with $\Delta q$ charge imbalance to generate $\Delta V$ ($\sim 0.5$-$1.1V$). Track $Cl^-$ and $Na^+$ crossings. | Fig. 6A-C |
|  | Conductance (pS) | Linear fit to the I-V relation from chloride permeation counts vs voltage. | Fig. 6D |
| Conformational Dynamics | Global twist angle ($\tau^\circ$) | Average angle between ECD and TMD $C_\alpha$ COM vectors projected onto the plane perpendicular to the symmetry axis (Wordom). Lower $\tau$ is associated with activation-like untwisting. | Fig. 7A |
| | M2 tilt (polar $\theta_p$, azimuthal $\theta_a^\circ$) | Helix axis from $C_\alpha$ least-squares; polar = polar radial tilt; azimuthal = tangential rotation; outward tilt accompanies opening. | Fig. 7B-C |
| | ECD β-sandwich tilts ($\theta_p$, $\theta_a^\circ$) | $C_\alpha$-based β-sandwich axes per subunit; reduced polar tilt indicates ECD compaction during activation. | Fig. 7E-F |
| | Orthosteric site closure proxy ($\text{\AA}$) | $\beta 3:Y205-\alpha 1:R120$ (specify atom pair in Methods); smaller distance indicates C-loop closure | Fig. 7G-I; S26-S27 |

### Changes in Pore Geometry

GaMD simulations revealed that all-subunit L9′T and L9′S mutants induce radial expansion of the pore at three key axial positions, the extracellular 20′ entry, the 9′ hydrophobic gate, and more modestly, the -2′ desensitization gate, relative to WT (**Figure 3A-C**). Time-averaged radius profiles along the pore axis show that the 9′ gate increases to ∼3.7-4.1 Å in the mutants, while the 20′ entry widens by ∼0.5-0.7 Å, and the -2′ gate fluctuates between 1.8 and 2.6 Å (**Figure 3D**; dynamic ranges are summarized in **Figure S2**). By contrast, the WT channel maintains a minimum pore radius <1.7 Å across the trajectory (**Figure 3E**), consistent with a non-conductive geometry. To assess the extent of pore accessibility, we computed a pore expansion frequency (PEF), the fraction of simulation frames with minimum radius > 2.0 Å, which reached ∼48% for both L9′T and L9′S but 0% for WT (**Figure 3F**).

**Figure 3.**
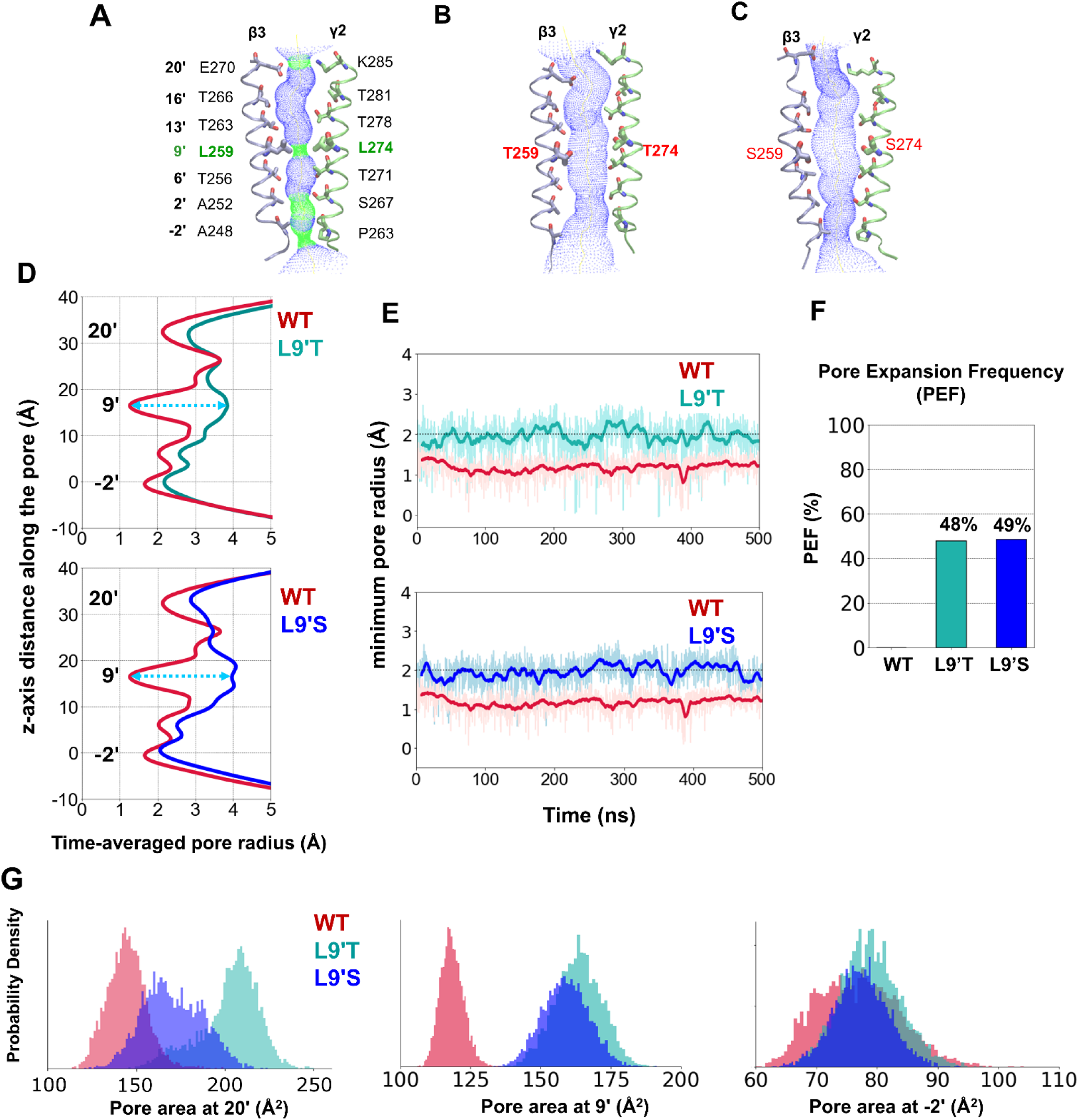
Structural dynamics of the channel pore in the WT, L9′T-and L9′S-mutant α_1_β_3_γ_2_ GABA_A_ receptors. (A-C) Membrane cross-sectional views of the channel pore formed by M2 helices from all five subunits. For clarity, only two opposing M2 helices (from β3 and γ2 subunits) are shown, with pore lining residues in licorice representation. Pore volumes were calculated using HOLE software, with blue surfaces indicating regions with radius >2.3 Å and green surfaces indicating 1.5 Å < r <2.3 Å. (D) Time-averaged pore radius profiles along the channel axis over 500 ns simulations, comparing WT (crimson) with L9′T (light seagreen) and L9′S (blue) systems. The significant differences in pore radii at 9’ upon mutations are highlighted in blue dotted lines. (E) Minimum pore radius values calculated per frame across the simulation trajectory. (F) Pore expansion frequency (PEF), defined as the percentage of frames with a minimum pore radius >2.0 Å, highlights the increased pore accessibility in mutant channels. (G) Density plots of cross-sectional pore areas at 20’, 9’, and -2’ positions computed from M2 Cα pentagons (see Methods). Data shows significant expansions at 20’ and 9’ positions, and moderate widening at -2’ gate in L9′T and L9′S mutants, where a larger area indicates greater backbone separation among M2 helices.

In addition, cross-sectional pore areas, computed from M2 Cα pentagons (mean nearest neighbor M2 Cα-Cα distance used as the side of a regular pentagon; see Methods) at axial positions. This backbone-based metric is sensitive to helical backbone separation (i.e., pore widening by backbone motion), complementary to HOLE radii, which can be influenced by sidechain rotamers. The results corroborate backbone-level expansion at 20′ and 9′, with a smaller increase at -2′ (**Figure 3G**). At the 20′ entry, the mean area increases from ∼145 Å^2^ in WT to ∼220 Å^2^ in L9′T and ∼170 Å^2^ in L9′S (expansion vs WT ≍ +75 Å^2^ and 25 Å^2^, respectively). At the -2′ gate, the area increase for both mutants is modest (∼ +18 Å^2^ vs WT), consistent with -2′ remaining the narrowest constriction. Full area distributions across 20′, 9′, 6′, 2′, and -2′ are provided in the Supporting Information (**Figure S4**), where the 6′ region typically shows an intermediate expansion in both mutants (∼ +35 Å^2^ vs WT). Together, the radius and area analyses indicate that L9′T and L9′S all-unit substitutions destabilize the 9′ hydrophobic gate and expand the 20′ entry, while leaving -2′ as the principal residual constriction.

### Hydration Status of the Channel Pore

Channel hydration is a critical determinant of ion permeability. A well-hydrated pore facilitates ion conduction, whereas a dewetted (dry) or occluded pore impedes it^39,40^. To assess hydration dynamics, we quantified two complementary readouts: (i) the number of water molecules near the 9′ hydrophobic gate and (ii) water flux across the pore. Water counts were measured within a 14 Å axial slab centered at the 9′ gate (i.e., 7 Å above and 7 Å below; see Methods). Flux was defined with Wordom as the number of complete water transits between the extracellular (20′) and intracellular (-2′) sides per unit time, independent of direction (the metric is direction-agnostic by design; as expected for equilibrium GaMD, net water displacement across the box was ∼0 over each trajectory).

In the WT receptor, the pore near 9′ was sparsely hydrated, with an average of ∼ 11.8±3.8 water molecules (**Figure 4A-B**). In contrast, the all-subunit L9′T and L9′S mutants displayed markedly higher water occupancy, ∼32.3 ± 5.9 and ∼33.5±5.4 molecules, respectively, consistent with the expanded radii and increased cross-sectional areas at 20′ and 9′.

**Figure 4.**
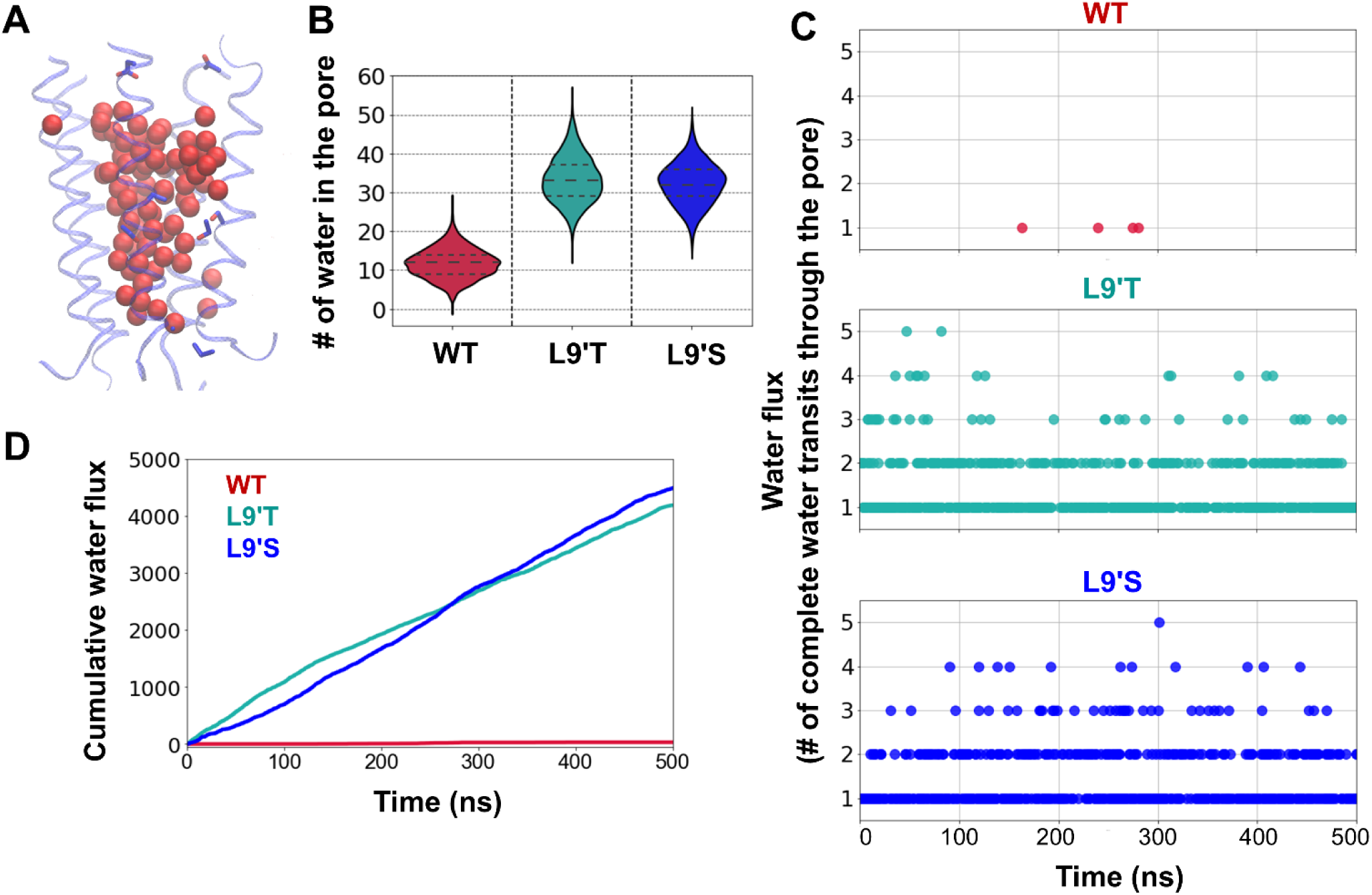
Hydration status of the channel pore in wild-type and L9′T/L9′S mutant α1β3γ2 GABA_A_ receptors. (A) Membrane view of the pore-lining M2 helices with water molecules shown as red spheres in the pore region. (B) Violin plots of water molecules within ±7 Å of the 9′ hydrophobic gate (7 Å above and 7 Å below; see Methods). Both L9′T and L9′S mutants exhibited significantly higher hydration than WT (t-test p <0.001). (C) Water flux vs. time (Wordom/MDAnalysis), defined as the direction-agnostic count of complete water transits between the extracellular (20′) and intracellular (-2′) sides per unit time. (D) Cumulative water flux (total number of complete transits) over the 500-ns trajectories. (Where indicated, distributions pool three independent GaMD replicates per system.)

Water flux increased commensurately with hydration. Over 500 ns of simulation, L9′T and L9′S each exhibited > 4000 complete transits, whereas the WT receptor showed < 200 (**Figure 4C-D**). Flux traces typically showed 2-3 water molecules in transit at any given time, with intermittent bursts of 4-5. These observations were reproducible across three independent GaMD replicates per system (pooled in the distributions). Together, these hydration metrics support a transition from a dewetted, non-conductive pore (WT) to wetted, conductive pores in the L9′T and L9′S all-subunit mutants, consistent with the observed pore expansion (**Figure 3**), reduced PMF barriers at 9′ and -2′ (**Figure 5**), and with the chloride conductance observed in CompEL (**Figure 6**).

**Figure 5.**
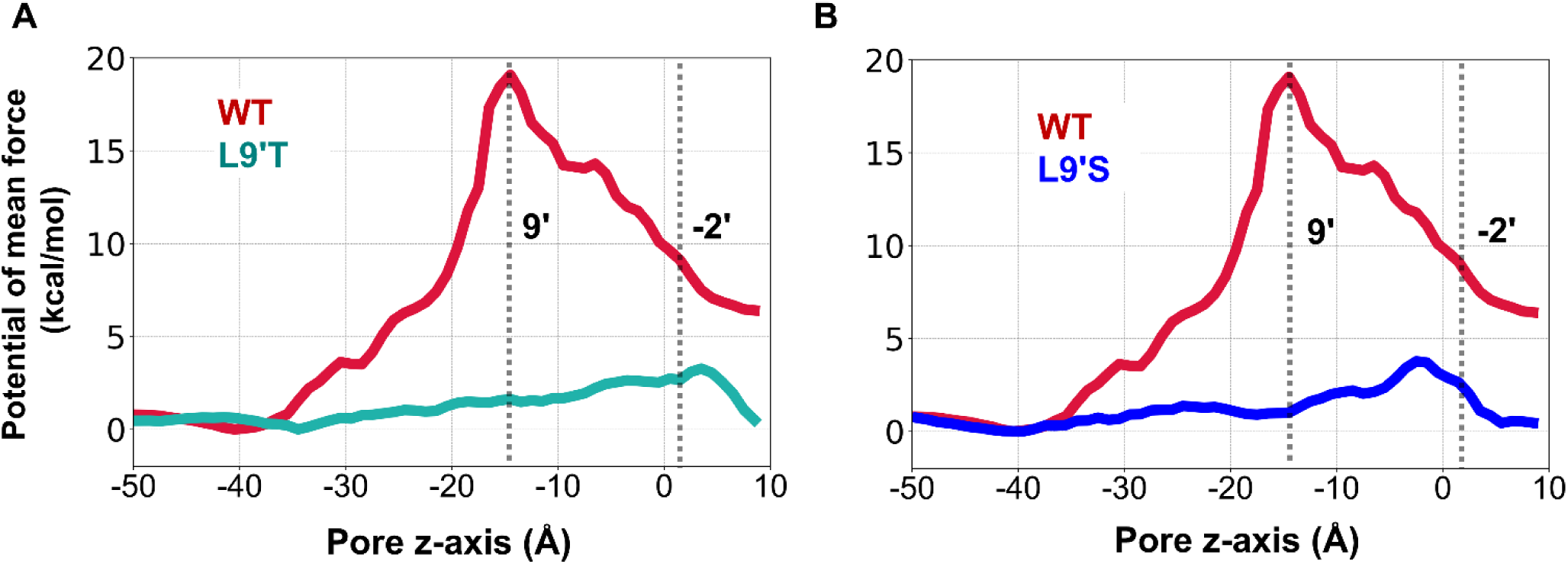
Free energy profiles (PMFs) for Cl^−^ permeation along the pore axis of wild-type and L9′T and L9′S mutant α1β3γ2 GABA_A_ receptors. (A) L9′T (light seagreen) compared with WT (red), (B) L9′S (blue) compared with WT (red). In both panels, the reaction coordinate is z along the pore axis (extracellular → intracellular), and vertical dotted lines mark the 9’ hydrophobic gate and -2’ desensitization gate positions. The WT PMF exhibits two dominant barriers, ∼19 kcal/mol at 9’ and ∼9 kcal/mol at -2’, whereas barriers are reduced in the mutants (L9′T: ∼1.6 and ∼3.0 kcal/mol; L9′S: ∼1.3 and 3.4 kcal/mol at 9’ and -2’, respectively). PMFs were obtained by umbrella sampling with 1 Å window spacing and 20 ns/window, reconstructed with WHAM (see Methods). Barriers are mean across replicas; replicate variability was ≤ ∼0.5 kcal/mol at barrier maxima (see Methods).

**Figure 6.**
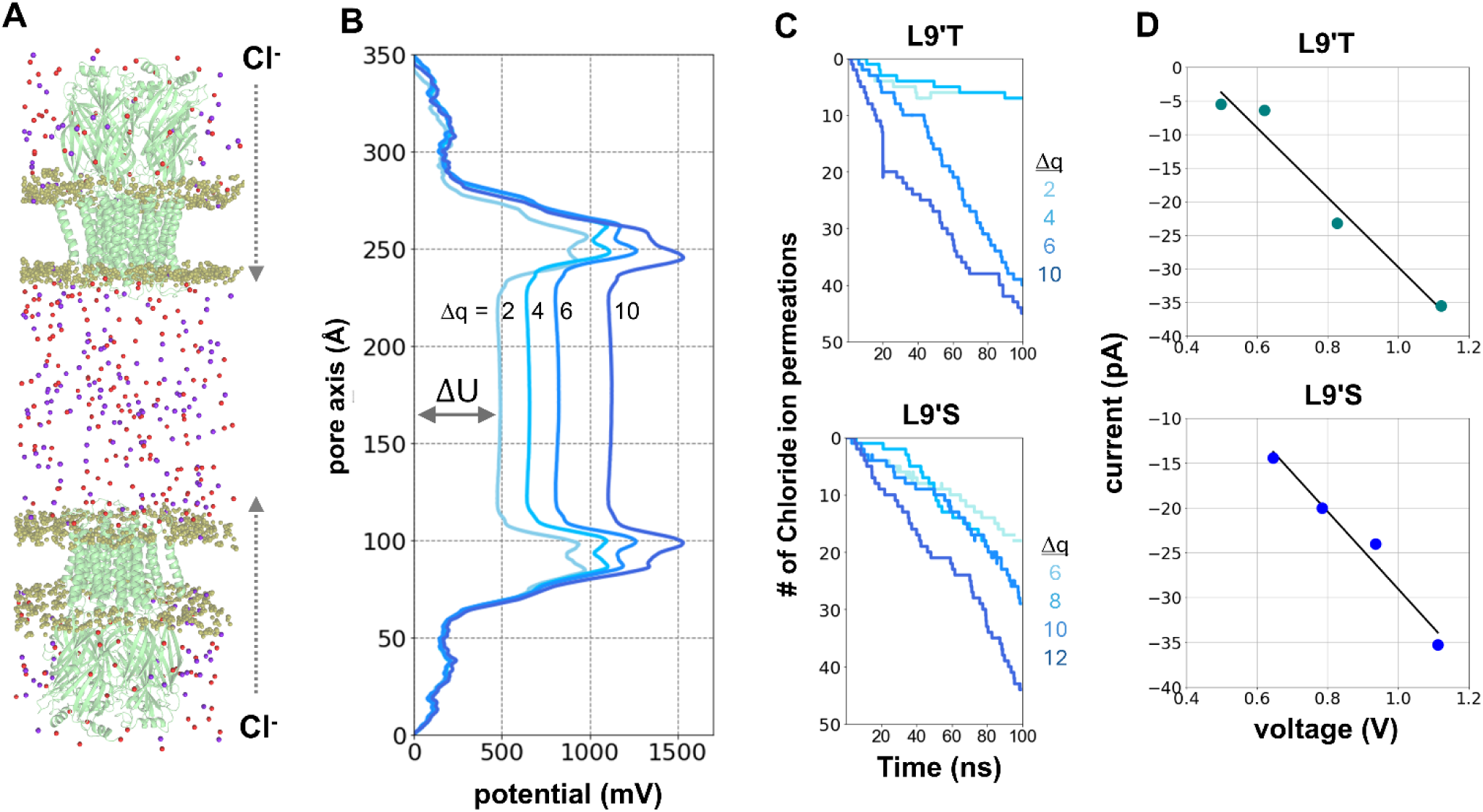
Computational electrophysiology (CompEL) of all-subunit L9′T and L9′S α1β3γ2 GABA_A_ receptors. **(A)** Schematic of the double bilayer CompEl setup built from GaMD-equilibrated snapshots. A charge imbalance (Δq) is imposed between compartments to generate a transmembrane potential (ΔU). **(B)** Electrostatic potential profiles (ΔU) along the z-axis for representative Δq values. **(C)** Time evolution of Cl^−^ permeation events for L9′T and L9′S. Na^+^ crossings were not observed under any simulated conditions. Swaps were restricted to a 1.0 nm radius cylinder at the 9’ plane to track channel permeation. (D) Current-voltage relations with linear fits, yielding conductance of ∼10-30 pS (L9′T) and ∼22-32 pS (L9′S) over the simulated voltage range. Conductance values are consistent with single-channel measurements in experimental systems.

### Energetics of Chloride Permeation

We characterized the free-energy landscape for chloride ion transport using steered MD (SMD) followed by umbrella sampling (US) simulations. A chloride ion was pulled along the pore axis (z-axis) from the extracellular vestibule to the intracellular exit, generating a series of conformational windows, spaced 1 Å apart (**Movie M1**). Each window was then subjected to 20 ns of restrained sampling to capture local conformational fluctuations (∼1.2 µs per system). Subsequently, the potential of mean force (PMF) was reconstructed using the Weighted Histogram Analysis Method (WHAM) to yield free energy profiles^41^.

In the WT closed-state receptor, the PMF reveals two major energy barriers: ∼19 kcal/mol at the 9’ hydrophobic gate and ∼9 kcal/mol at the -2’desensitization gate **(Figure 5A)**. These barriers effectively prevent chloride permeation and are consistent with the dewetted, constricted geometry observed for WT.

In the all-subunit mutants, both barriers are dramatically reduced **(Figure 5A-B).** For L9′T, the energy barrier falls to ∼1.6 kcal/mol at the 9’ gate and to ∼3 kcal/mol at the -2’ gate; for L9′S, the barriers fall to ∼1.3 kcal/mol (9’) and to ∼3.4 kcal/mol (-2’). These reductions reflect the increased pore radius and hydration at the corresponding axial positions (**Figures 3-4**) and explain the chloride conductance observed in CompEL (**Figure 6**). Across replicate PMFs, the barrier heights were reproducible (variation ≤ ∼0.3-0.5 kcal/mol at the barrier maxima; see Methods/Errors), indicating robust convergence of the dominant features.

Overall, the L9′T/L9′S substitutions destabilize hydrophobic gating at 9’ and partially lower the -2’ constriction, reshaping the energy landscape from non-conductive (WT) to conductive with a residual bottleneck at -2’.

### Computational Electrophysiology (CompEL)

To assess conductive behavior directly, we performed CompEL simulations on the all-subunit L9′T and L9′S mutant α1β3γ2 GABA_A_ receptors (see Methods)^42,43^. Starting structures were taken from well-equilibrated GaMD snapshots (see Methods), duplicated along z to build a double-bilayer system, and equilibrated prior to production (**Figure 6A**). Ion swaps were restricted to a 1.0 nm radius cylindrical region centered on the pore at the 9’ plane so that counted events reflect channel permeation rather than bulk exchanges.

A set of charge imbalances (Δq = 2, 4, 6, 8, and 10) was applied across the double-bilayer system to generate transmembrane potentials (ΔU) spanning ∼500-1100 mV across the tested conditions (**Figure 6B**; exact Δq → ΔU mapping and potential profiles are in Methods). Under these voltages, both mutants exhibited robust Cl^−^ permeation that increased with ΔU (**Figure 6A-C, Movie M2**), while no Na^+^ permeation events were observed under any simulated condition, indicating strong anion-selectivity (**Figure 6C**). The resulting current-voltage (I-V) relations were approximately linear (ohmic), yielding conductance of ∼10-30 pS for L9′T and ∼22-32 pS for L9′S (**Figure 6D**). These values are in good agreement with single-channel conductance measured in in vitro and in vivo preparations of functional GABA_A_Rs^44,45^, and they are consistent with the hydrated, expanded pores seen in GaMD (**Figure 3-4**) and lower PMF barriers at 9’ and -2’ (**Figure 5**). Results were reproducible across independent CompEL runs initiated from equilibrated GaMD structures (see Methods); linear fits report mean slopes with small variability across replicates.

### Conformational dynamics

Hydrophilic L9′T and L9′S mutations induce significant changes in pore geometry, hydration, and conductance, accompanied by conformational isomerization of the receptor’s quaternary structure. We quantified global twisting of the receptor and local tilts of the pore-lining M2 helices and ECD β-sandwiches using Cα-based least-square axes (Wordom; see Methods for residue ranges). By convention, positive polar tilt denotes outward (radial) tilt from the channel axis, and positive azimuthal tilt denotes clockwise rotation when viewed from the extracellular side; global twist (τ) is the angle between projected ECD and TMD center-of-mass vectors on a reference plane perpendicular to the receptor’s symmetry axis^46^.

The WT closed-state receptor exhibited a mean global twist angle of ∼21.5° (SD = ±1.1°), whereas the L9′T and L9′S all-subunit mutants showed reduced twist of ∼18.9° (SD = ±0.9°) and 19.4° (SD = ±1.0°), respectively **(Figure 7A)**, consistent with the untwisting typically associated with activation in pLGICs^46–49^. Mutants also showed increased M2 helix tilt relative to WT: L9′T displayed a ∼+2° increase in both polar and azimuthal components, while L9′S primarily increased azimuthal tilt (**Figure 7B-C**). In the ECD, β-sandwich polar tilts decreased mostly in both mutants, indicating compaction of the extracellular domain (**Figure 7E-F**), a motion that is associated with channel activation^46^.

**Figure 7.**
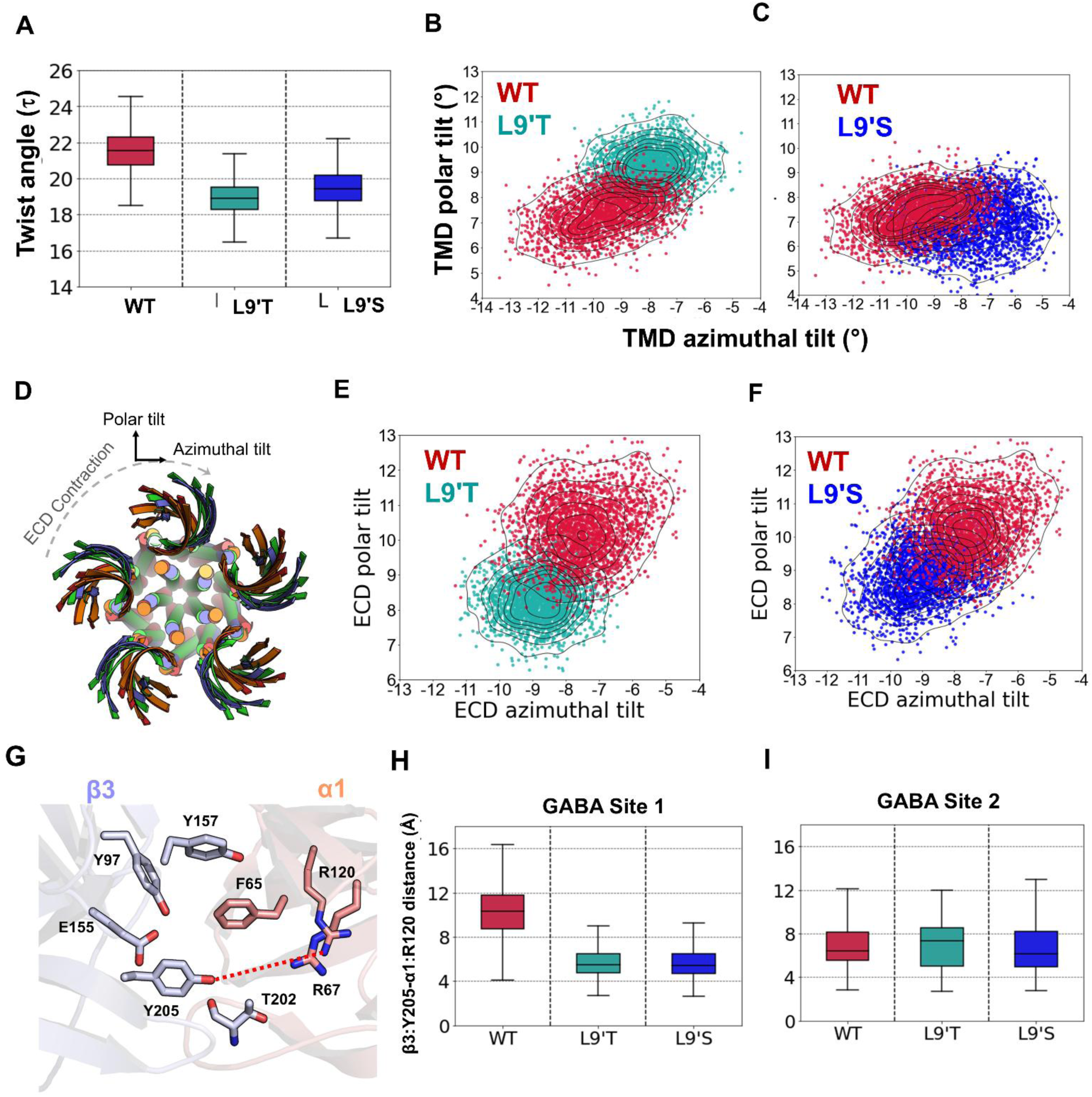
Global and local conformational dynamics of WT and L9′T/L9′S α1β3γ2 GABA_A_ receptors. **(A)** Box plots show the distribution of global twist angle for WT (red), L9′T (light seagreen), and L9′S (blue) receptors over 500 ns GaMD trajectories. **(B-C)** 2D scatter plots of polar versus azimuthal tilt angles of the pore-lining M2 helices for WT vs L9′T and WT vs L9′S; each point represents a simulation frame. **(D)** Schematic diagram of the top view of the α1β3γ2 GABA_A_ receptor showing the β-sandwiches of the extracellular domain and helices of the transmembrane domain defining the sign conventions used for polar (outward) and azimuthal (clockwise, extracellular view) tilts and for the global twist angle. (**E-F**) Scatter plots of ECD β-sandwich polar versus azimuthal tilt angles for WT vs L9′T (E) and WT vs L9′S (F). (**G**) Cartoon representation of the orthosteric binding interface, highlighting the distance between β3:Y205 and α1:R120 used as a proxy for C-loop closure. (**H-I**) Box plots showing the distribution of the β3:Y205-α1:R120 inter-residue distances at two GABA-binding orthosteric sites 1 (H) and 2 (I) in WT (red), L9′T (light seagreen), and L9′S (blue) receptors. All angles were calculated from Cα-based least-squares axes using Wordom (see Methods for residue ranges and atom definitions). Positive polar tilt denotes outward motion from the receptor axis; positive azimuthal tilt denotes clockwise rotation viewed from the extracellular side. Boxes indicate the median and interquartile range, with whiskers denoting the full data range unless stated otherwise. Distributions pool independent GaMD replicates.

At the orthosteric interfaces, the β3:Y205 (oxygen atom from the sidechain -OH)-α1:R120 (amino group from the sidechain guanidine) distance for one of the binding sites decreased in both mutants relative to WT (**Figure 7G-I**), consistent with C-loop closure over the agonist binding sites that accompanies activation. Across independent replicas, angle distributions and orthosteric distances were stable (variability typically within ∼0.3-0.6° for angle medians and ∼0.2-0.4Å for distance medians), supporting convergence of these conformational signatures.

### Single- and Double-Subunit Variants

To assess whether partial 9′ hydrophilic substitution captures intermediate states, we performed the same analyses for single- and double-subunit L9′T/L9′S mutants (Supporting Information, **Figures S9-S27**). In brief, single-subunit variants generally maintained sub-conductive minima (low PEF) with reduced hydration and higher PMF barriers relative to all-subunit mutants, while selected double-subunit combinations exhibited intermediate pore expansion, sustained water flux, and partial barrier reduction. Conformational signatures (reduced twist, increased M2 tilt, orthosteric tightening) were directionally consistent but of smaller magnitude than in all-subunit L9′T/L9′S. These trends support a graded stabilization of open-like ensembles with increasing 9′ hydrophilicity (see **Figures S14-S20** for hydration/flux; **S11, S16** for PEF/PMF; **S21-S27** for angles and orthosteric distances).

### Effects of Positive Allosteric Modulation on L9’T

Hydrophilic L9′T substitutions stabilize an open-like pore that provides a permissive background for probing positive allosteric modulation. We therefore examined the effects of ganaxolone (GAN) and diazepam (DZP) on the all-subunit L9′T α1β3γ2 GABA_A_ receptor. Ganaxolone binds within the transmembrane domain^50,51^ at a neurosteroid site formed primarily by the β3 M3 helix and neighboring α1 M1/M4 helices (**Figure 8A**), approximately ∼10 Å from the -2′ gate. In the presence of GAN, the L9′T receptor exhibited a higher pore expansion frequency (∼69%), compared to the apo L9′T system (**Figure 8C-D**). This increase was accompanied by a selective expansion at the -2′ desensitization gate, where the mean radius increased to ∼2.5 Å (range ∼2.2–2.7 Å), while the 9′ gate remained largely unchanged, consistent with the gate already being destabilized by the L9′T substitutions (**Figure 8B**).

**Figure 8.**
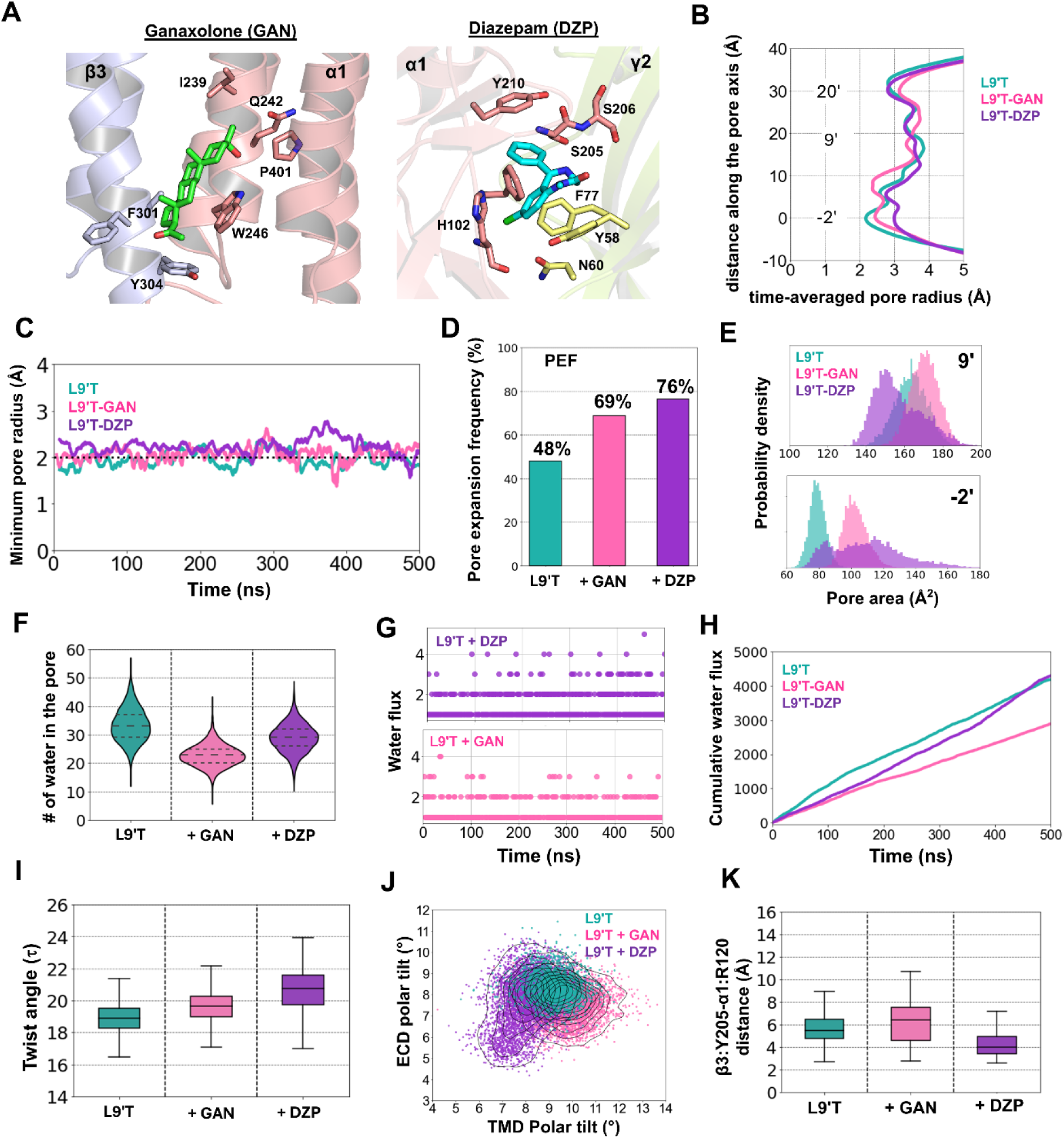
Positive allosteric modulation of all-subunit L9′T α1β3γ2 GABA_A_ receptors. (**A**) Cartoon representations of PAM binding sites: diazepam (DZP, cyan) at the α1/γ2 interface and ganaxolone (GAN, green) at the transmembrane neurosteroid site in licorice representation. Important binding site residues at α1 (salmon), β3 (light blue), and γ2 (lime) subunits are shown. (**B**) Time-averaged pore radius profiles along the channel axis for apo L9′T (light seagreen), L9′T+DZP (purple), and L9′T+GAN (pink) are shown. (**C**) Time series plots of the minimum pore radius for the three systems. (**D**) Bar plot of pore expansion frequency (PEF), defined as the fraction of simulation frames with minimum pore radius >2.0 Å. (**E**) Density distributions of pore cross-section areas at 9′ and -2′ positions, highlighting PAM-induced expansion. (**F**) Violin plots of the total number of water molecules within 7 Å of the 9′ gate, both sides. (**G**) Time series plots of water flux (direction-agnostic complete transits; Wordom). (**H**) Cumulative water flux over 500 ns. (**I**) Box plots of the global twist angle. Both PAMs show modest increases in twist yet remain lower than the WT closed-state value, consistent with stabilization of an open-like ensemble. (**J**) 2D Scatter plots of TMD polar tilt (x-axis) versus ECD polar tilt (y-axis) angles. In this representation, activation corresponds to higher TMD polar tilt (→right) and lower ECD polar tilt (↓ down), i.e., a shift toward the lower-right region. Both PAMs maintain or slightly accentuate this activated-like coupling relative to apo L9′T. (**K**) Box plots of the β3:Y205-α1:R120 distance, reporting orthosteric C-loop closure. All quantities were computed from GaMD trajectories of the all-subunit L9′T receptor. Plot conventions, angle sign definitions, and statistical representations are described in Figure 7 and Methods.

Cross-sectional area analysis corroborated this effect, showing an ∼20 Å^2^ increase at the -2′ position, indicative of backbone-level widening rather than side-chain rearrangement.

Diazepam binds distally at the α1/γ2 benzodiazepine site in the extracellular domain^52,53^ (**Figure 8A**), yet it produced effects similar to GAN. DZP increased the pore expansion frequency to ∼72% and promoted enhanced opening at the 2′ desensitization gate, with a mean radius approaching ∼3 Å and increased variability across frames (**Figure 8D-E**).

Both PAMs maintained a hydrated pore relative to WT but displayed comparable or slightly reduced water flux and cumulative counts compared to the apo L9T system (**Figure 8F-H**). Despite enhanced pore expansion frequency and increased accessibility at the -2′ gate, GAN- and DZP-bound receptors displayed reduced stochastic water exchange, suggesting that PAM binding stabilizes an already open-like pore while dampening large-amplitude fluctuations in water permeation. This behavior is consistent with PAMs biasing the conformational ensemble toward more ordered open states, rather than increasing overall pore wetting.

To assess, at a global level, how PAMs bias activation-linked quaternary motions, we quantified the ECD global twist angle. Relative to apo-L9′T, GAN and DZP induced modest increases in the global twist angle (∼0.6 and 1.3, respectively; **Figure 8I**) while maintaining values below those of the WT receptor (**Figure 8I**). These small shifts are consistent with fine-tuning of an already open-like ensemble rather than a transition toward a closed or desensitized conformation. In addition, we plotted TMD polar tilt (x-axis) versus ECD polar tilt (y-axis) for the all-subunit L9′T receptor in apo and GAN-and DZP-bound states (**Figure 8J**). In our system, activation corresponds to increased TMD polar tilt (outward tilt of M2) and decreased ECD polar tilt (β-sandwich compaction), i.e., a shift toward the lower-right region of the plot. Relative to apo-L9′T, both GAN and DZP maintain or modestly accentuate this activated-like coupling (→ right in TMD polar; → down in ECD polar), consistent with stabilization of an open-like ensemble rather than a distinct opening mechanism. (Angle sign conventions follow Figure 7D; positive polar = outward). Finally, both ligands stabilized orthosteric C-loop closure, as reflected by sustained β3:Y205- α1:R120 distances (**Figure 8K**).

### Effects of Negative Allosteric Modulation on L9’T

We next examined how the orthosteric antagonist bicuculline (BIC) and pore blocker picrotoxin (PTX) influence the all-subunit L9’T α1β3γ2 receptor, using the same set of pore, hydration, and conformational observables as in the PAM section (cf. **Figure 8**).

Relative to apo-L9’T, BIC^28,54^ reduces pore expansion frequency (PEF) and re-constricts the pore at both 9’ and -2’ (**Figures 9B-E**). In time-averaged radius profiles, 9’ narrows substantially from the L9’T expanded state, and the -2’ gate shifts toward smaller radii with increased variability (**Figure S3**).

**Figure 9.**
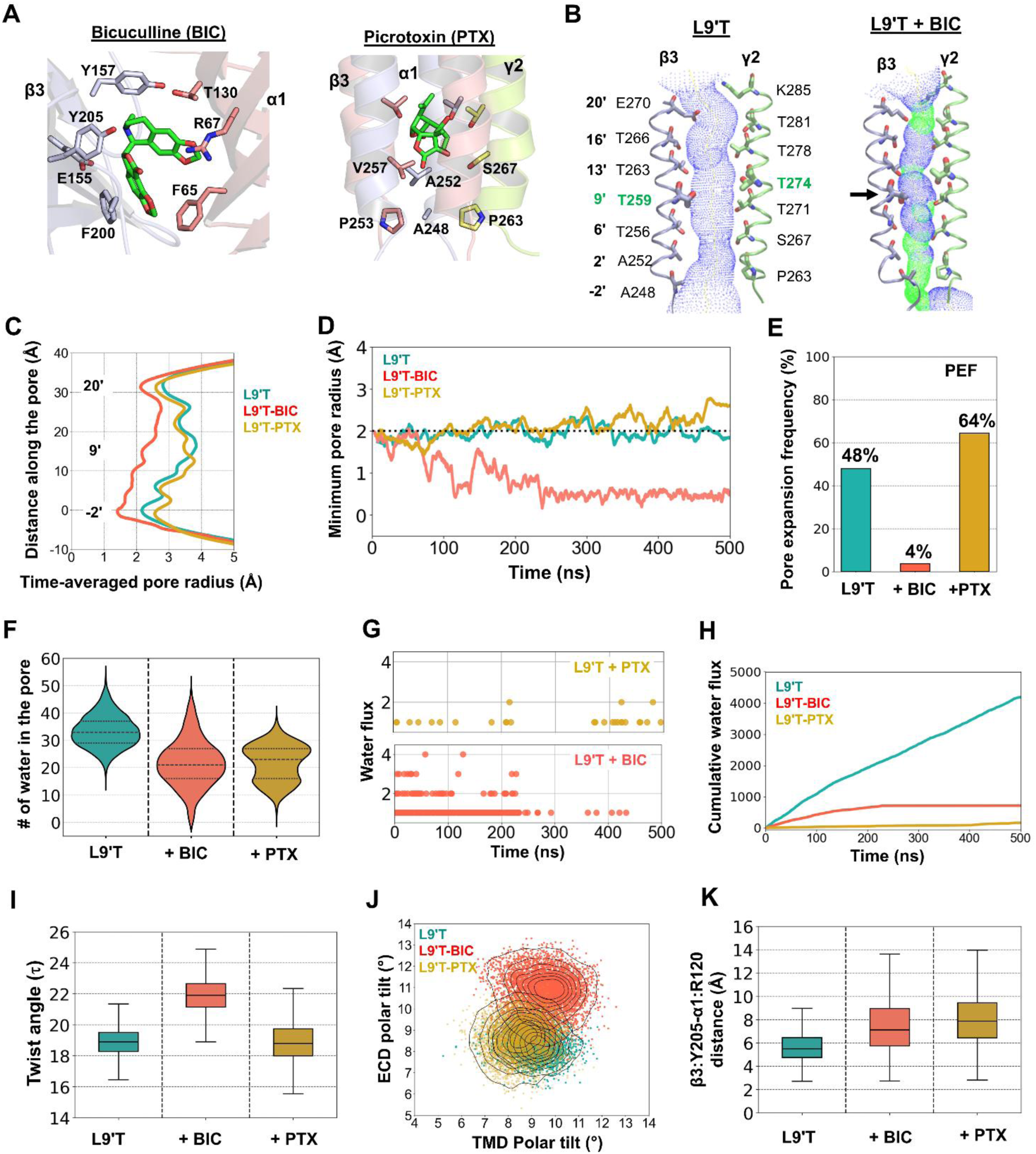
Effects of negative allosteric modulators on all-subunit L9′T α1β3γ2 GABA_A_ receptors. (**A**) Cartoon representations of the receptor showing bicuculline (BIC) at the orthosteric β3/α1 interface and picrotoxin (PTX) lodged within the pore between 9’ and -2’ positions. Ligands are in licorice representation (green). (**B**) Pore volume renderings (HOLE surfaces) for apo-L9’T and L9’T+BIC, analogous to Figure 3B-C. Blue surfaces indicate regions with radius > 2.3 Å, green indicates 1.5 Å < r <2.3 Å, and red (if present) r <1.5 Å. The BIC-bound structure shows visible constrictions at the 9’ and - 2’ relative to apo-L9’T. (**C**) Time-averaged pore-radius profiles along the channel axis for apo-L9′T (light seagreen), L9′T+ BIC (orange), and L9′T+PTX (gold). (**D**) Time series of the minimum pore radius per frame. For PTX, the minimum radius can be as large as or slightly larger than apo-L9′T because the ligand’s physical presence locally expands the geometric radius even though the pore is functionally occluded. (**E**) Bar plot of pore expansion frequency (PEF), fraction of simulation frames with minimum pore radius > 2.0 Å. PTX shows a high PEF (∼64%) due to ligand-induced geometric expansion rather than a conductive state; BIC lowers PEF by re-constricting the pore. (PEF is a radius-based metric and does not account for occlusion by PTX). (**F**) Violin plots of the number of water molecules within ±7 Å of the 9’ gate along the pore axis (see Methods for details). (**G**) Time series of water flux (direction-agnostic complete transits between 20’ and -2’; Wordom). (**H**) Cumulative water flux over 500 ns, illustrating reduced flux with BIC and near-zero flux with PTX relative to apo- L9′T. (**I**) Box plots of the global twist angle (apo-L9’T vs BIC vs PTX). (**J**) 2D scatter plot of TMD polar tilt (x-axis) versus ECD polar tilt (y-axis). In this representation, activation corresponds to higher TMD polar tilt (→ right) and lower ECD polar tilt (↓ down) (see Figure 7D for sign conventions). BIC shifts the ensemble toward lower TMD polar and higher ECD polar (closed-like coupling), whereas PTX clusters near apo-L9’T, consistent with pore block without major quaternary reorientation. (**K**) Box plots of β3:Y205-α1:R120 inter-residue distance (orthosteric C-loop proxy). In BIC, this distance is already high at the start and remains elevated, consistent with steric prevention of C-loop closure rather than a time-dependent “opening”; PTX produced a similar increase.

Cross-sectional area distributions corroborate backbone-level contraction across the lower pore (**Figure 9C-E**). Hydration metrics reflect these geometric changes: water counts near 9’ decrease and water flux and cumulative counts are reduced relative to apo-L9’T (**Figure 9F-H**). In the ECD-TMD coupling view (**Figure 9J**; see axis details below), BIC shifts the ensemble away from the activated-like quadrant (lower ECD polar, higher TMD polar) toward higher ECD polar and lower TMD polar, consistent with a partial return to a closed-like ECD-TMD coupling. Global twist increases modestly toward the WT-closed value (**Figure 9I**), further supporting a closing bias.

### Relation to antagonist-bound cryo-EM structures

The bicuculline effects we observe, including reduced PEF, contraction at 9’ and -2’, lower hydration/flux, increased global twist, and an ECD-TMD coupling shift toward higher ECD polar tilt and lower TMD polar tilt, are consistent with antagonist-bound cryo-EM conformations of heteropentameric GABA_A_Rs that exhibit inhibited/closed-like extracellular arrangements and less permissive pores at the lower TMD interface^52,55^. In particular, orthosteric antagonists prevent C-loop closure^15^, and we likewise find larger β3:Y205-α1:R120 distances with bicuculline than in apo-L9′T (**Figure 9K**), paralleling those structural signatures. While our L9′T background is an open-like ensemble and cryo-EM structures are typically obtained in WT receptors, the direction of the bicuculline-induced changes aligns with the closed-bias seen in antagonist-bound maps, supporting a model in which orthosteric inhibition destabilizes ECD-TMD coupling and re-closes the residual -2’ constriction that remains after 9’ hydrophobic gating is relieved.

Picrotoxin (pore blocker)^52,53^. PTX produces near-complete suppression of water flux and cumulative transit counts with minimal changes in global quaternary metrics (**Figure 9F-I**). Because PTX is physically lodged within the pore, the time-averaged radius at the constriction can be as large as or slightly larger than apo-L9’T, and the pore expansion frequency (PEF) reaches ∼64% (**Figure 9D-E**). This reflects a geometric expansion caused by the ligand’s presence rather than a conductive state, i.e., the radius-based PEF does not account for occlusion, which explains why flux remains suppressed despite an apparently large minimum radius. Consistently, the apparent increases in geometric radius/area between 9’ and -2’ reflect steric occlusion of the pore by PTX rather than a conductive configuration. In the ECD-TMD polar-tilt plot (TMD polar on x-axis, ECD polar on y-axis; **Figure 9J**), PTX points cluster near apo-L9’T, supporting block without major quaternary rearrangement. The orthosteric C-loop metric (β3:Y205-α1:R120) shows small shifts relative to apo-L9’T (**Figure 9K**), far smaller than those seen with BIC, consistent with pore block rather than orthosteric antagonism.

In summary, BIC reduces accessibility and hydration, drives a shift toward closed-like ECD-TMD polarity, and modestly increases twist, consistent with re-closing the open-like L9’T ensemble. PTX primarily halts permeation via steric occlusion with minimal global conformational change.

### Bicuculline-Induced Closing Sequence in L9’T

The bicuculline-bound L9′T simulations reveal a stepwise, bottom-to-top closing sequence that progresses from an open-like to a closed-like ensemble (**Figure 10A-C**). For context, panel C compares each time-binned metric for L9′T+BIC against reference distributions from the closed WT (PDB ID: 7QNE)^13^ and a desensitized-like WT (PDB ID: 6HUO) ensemble, as well as the apo-L9′T baseline.

**Figure 10.**
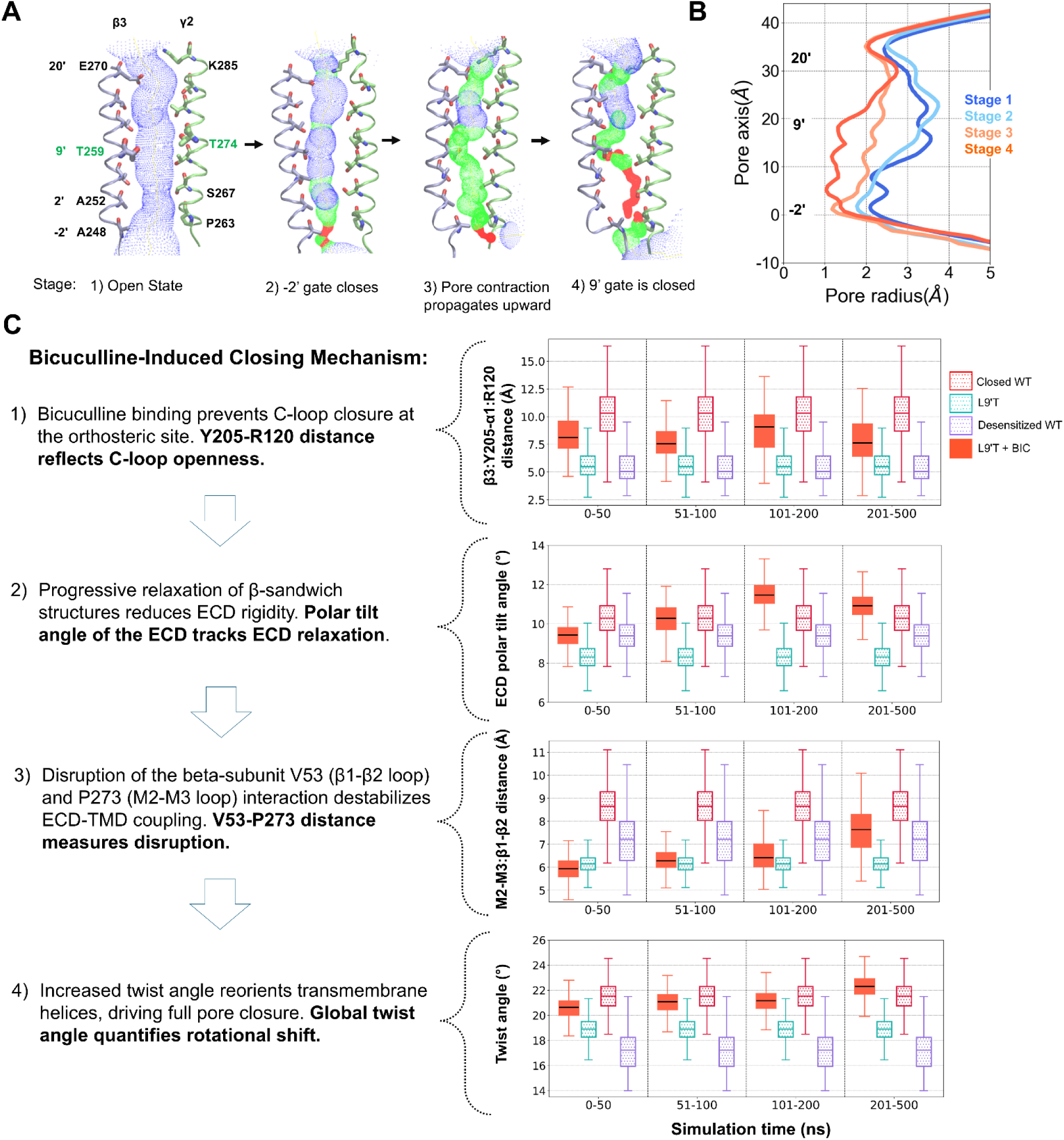
Bicuculline-induced closing mechanism of the open-state all-subunit L9′T α_1_β_3_γ_2_ GABA_A_ receptor with reference state comparisons. (**A**) Pore volume renderings (HOLE) of representative Stage 1-4 snapshots under bicuculline (apo-like → closed-like). Blue, green, and red surfaces indicate pore radius >2.3 Å, 1.5 Å<r<2.3 Å, and r<1.5 Å, respectively. Stage 2 shows selective -2’ constriction with 9’/20’ still expanded; stage 3-4 show propagation of constriction to 9’ and 20’. (**B**) Time-averaged pore radius profiles along the z-axis for each stage of the bicuculline sequence. Open-like stages are shown in blue (Stage 1: dark; Stage 2: light) and closed-like stages in orange-red (Stage 3: light; Stage 4: dark). The curves illustrate the gradual migration of the pore constriction from -2’ toward 9’/20’, culminating in Stage 4, where the minimum radius between -2′ and 9′ falls <1.5Å. (**C**) Time-binned boxplots (0-50, 51-100, 101-200, 201-500 ns) under L9’T + BIC (orange-filled), plotted alongside closed WT (red-dotted), desensitized-like WT (purple-dotted), and apo-L9’T (light-seagreen-dotted) references for four structural observables. Panels: (1) Orthosteric C-loop proxy (β3:Y205-α1:R120 inter-residue distance); BIC sits near closed WT, above apo-L9’T; desensitized-like is intermediate or slightly above apo-L9’T; larger values indicate C-loop cannot close under bicuculline, (2) ECD polar tilt; BIC > apo-L9’T and desensitized-like, approaching closed WT consistent with β-sandwich relaxation, which reduces ECD rigidity, (3) V53 (β1-β3)-P273 (M2-M3) distance; BIC > apo-L9’T and approaching closed WT, trending toward increase indicates loosening of ECD-TMD coupling, (4) Global twist angle; increase by ∼2° after 200 ns, approaching closed WT and remaining above apo-L9’T and desensitized-like, consolidates a closed-biased quaternary geometry. Angle sign conventions and atom/axis definitions are as in Methods (see Figure 7D for tilt/twist signs). Reference boxes for closed WT and desensitized-like WT provide state landmarks for each metric; apo-L9’T shows the open-like baseline from which bicuculline drives the bottom-to-top closure (-2’ first, then 9’/20’, then twist). All metrics are Cα-based unless noted; box plots show median, IQR, and whiskers to full range. Colors/line styles match the legend used in panel C.

Across all time bins, L9′T + BIC aligns most closely with the closed WT landmark for the β3:Y205-α1:R120 orthosteric distance (C-loop cannot close under BIC), while apo-L9′T is distinctly lower; the desensitized-like WT lies intermediate or modestly above apo-L9′T depending on site. ECD polar tilt under BIC increases relative to apo-L9′T and approaches the closed WT band (typically remaining at or just above the closed WT reference), consistent with β-sandwich relaxation. The P273-V53 separation (ECD-TMD coupling) grows beyond apo-L9′T and approaches the closed WT landmark, trending toward desensitized-like values as the interface loosens. Finally, global twist under BIC rises by ∼2° after ∼200 ns, approaching (and slightly exceeding) closed WT, while remaining distinct from apo-L9′T. Together, these comparisons indicate that bicuculline biases the L9′T ensemble toward a closed-like configuration while sampling desensitized-like coupling for the ECD-TMD interface, i.e., a state distinct from both the open-like apo-L9′T and any single WT reference state.

**Stage 1 (≍ 0-50 ns; open-like baseline).** Pore geometry closely resembles apo-L9′T, with time-averaged radii > 2 Å across most of the channel axis and the minimum radius typically above the conductive threshold (**Figure 10B**).

**Stage 2 (onset ≍ 50-100 ns; -2’ constriction first).** The first major event is selective constriction at the -2′ gate, while 9’ and 20’ gates remain expanded, a desensitized-like configuration in which -2’ is the primary constriction and the 9’ hydrophobic gate remains permissive (**Figure 10A-B**). Beginning ∼50-100 ns, the ECD polar tilt starts to increase (β-sandwich relaxation), marking early ECD changes that accompany the lower-pore constriction (**Figure 10C, ECD polar-tilt panel**).

**Stage 3 (≍ 100-200 ns; upward propagation).** Pore closure propagates upward from the intracellular end: constrictions appear at 9’ and -20’, and the minimum-radius location migrates from -2’ toward 9’/20’ (**Figure 10 A-B**). Around ∼200 ns, we observe a discernible increase in the P273(M2-M3)-V53(β1-β2) separation, indicating loosening of ECD-TMD coupling across the interface (**Figure 10C, P273-V53 panel**).

**Stage 4 (>200 ns; closed-like).** The minimum radius between -2’ and 9’ falls <1.5 Å, consistent with a closed-like pore (**Figure 10A-B, Stage 4**). After ∼200 ns, the global twist increases by ∼2° relative to early frames, consolidating the shift toward a closed-biased quaternary geometry (**Figure 10C, global-twist panel**).

### Asymmetric M2 collapse and implications for pore metrics

During bicuculline-driven closure, we observed a characteristic asymmetric collapse of the M2 bundle: three pore-lining helices converge while two are displaced laterally (**Movie M3**). This geometry reduces the HOLE-defined radius at -2′ first and then at 9′/20′ **(Figure 10B)**, eliminates water flux, yet can inflate the M2-Cα pentagon area because the two displaced helices increase the mean nearest-neighbor spacing used by the regular-pentagon approximation. To avoid this artifact, we recalculated the backbone cross-section by excluding the two laterally displaced M2 helices and averaging the three-helix distances from the convergent side (Methods), which yielded a monotonic area decrease that correctly tracks pore closure (**Figure 11**). Notably, the observed asymmetry is consistent with time-resolved cryo-EM reports of short-lived asymmetric intermediates^56^ (open, closed, desensitized) featuring subset-specific M2 translations/rotations and lipid access to the pore; orthosteric antagonism (bicuculline) appears to bias the L9′T open-like ensemble into such an asymmetric TMD collapse that precedes/quasi-coincides with -2′ first closure, and then propagates to 9′/20′, consolidating a closed-biased quaternary geometry.

**Figure 11.**
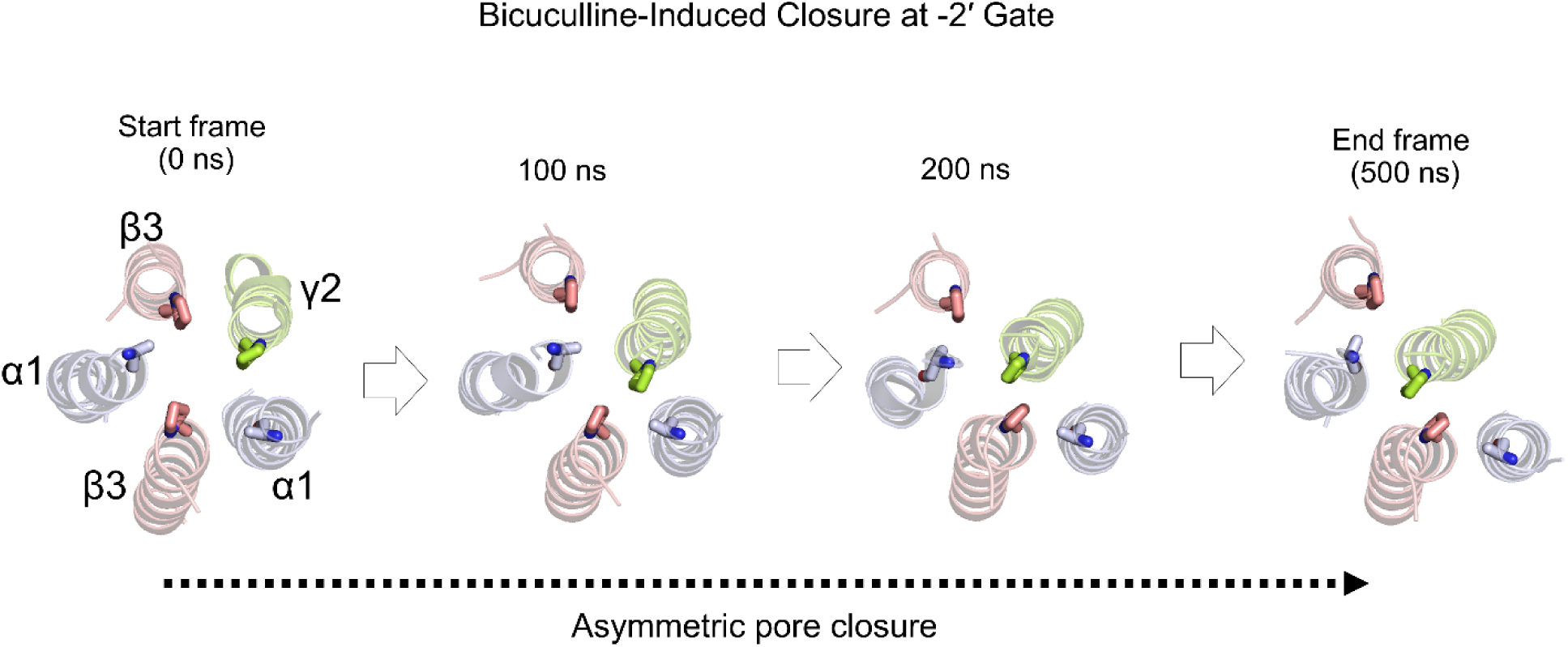
Asymmetric M2 collapse during bicuculline-induced closure at the -2′ gate (L9′T background). Bottom (Intracellular) view snapshots of the pore-lining M2 helices at the -2′ plane show a time-ordered collapse in the presence of bicuculline from start (0 ns) to end (500 ns) with two intermediates at 100 and 200 ns timepoints. Three M2 helices α1(1), β3(2), and γ2 converge toward the pore axis, while two helices β3(1), α1(2) are displaced laterally, producing an asymmetric constriction. This geometry accompanies the -2′ first closure that precedes narrowing at -0 and 20 (See Figure 10B for staged radii) and is consistent with loss of hydration and water flux at this plane. Sunit identifies are indicated (α1, β3, γ2). The panel illustrates structure-only changes; quantitative radius, hydration, and cross-section metrics are presented in Figure 9. Also, see Movie M1.

### Conformational signatures (Figure 10C panels)

**Orthosteric site (C-loop):** β3:Y205-α1:R120 distance is already elevated at t=0 and remains high under BIC (C-loop cannot close).

**ECD relaxation:** ECD polar tilt begins to increase at ∼50-100 ns (β-sandwich expansion) **(Figure 10C, panel 2).**

**ECD-TMD decoupling:** the P273-V53 distance (of the β3 subunit) increases around ∼200 ns, indicating loosening at the ECD-TMD interface **(Figure 10C, panel 3).**

**Global twist:** increases after ∼200 ns by ∼2° relative to the initial window, signaling a shift toward a closed-like conformation distinct from the initial open-state L9′T model and the desensitized state.

Viewed in reverse, the sequence suggests a plausible opening path (-2′ relief following early 9′ wetting), but the earliest steps collapse into our sampling window when 9′ is made hydrophilic (L9′T/L9′S), consistent with rapid early events below our GaMD timescale (Figure 2).

## DISCUSSION

By stabilizing open-like ensembles via hydrophilic substitutions at the 9′ hydrophobic gate (L9′T/L9′S) in the heteropentameric α1β3γ2 GABA_A_ receptor, we link local gate hydration/expansion to lowered permeation barriers and anion conductance. This relationship is internally consistent across observables and externally aligned with canonical pLGIC activation motifs. Mutants exhibit radial expansion at 9′ and 20′ with a residual -2′ bottleneck (HOLE radii; M2 Cα-pentagon areas), increased water occupancy/transits near 9′ (∼3-4 fold higher than WT), PMF barrier reductions at 9′ (∼1.3– 1.6 kcal/mol) and -2′ (∼3.0-3.4 kcal/mol) relative to WT, and ohmic conductance (∼10-32 pS) in CompEL that agree with known single-channel ranges for GABA_A_Rs. Conformationally, L9′T/L9′S display reduced global twist, increased M2 tilt, modest ECD compaction, and shorter β3:Y205-α1:R120 distances (C-loop closure), i.e., activation-like signatures described across pLGICs^1,46,47^. Together, these observations support a framework in which hydrophilic destabilization of the 9′ gate lowers the energetic cost of conduction while biasing the quaternary motions associated with opening.

On the L9′T background, PAMs (DZP, GAN) preferentially affect the -2′ gate (larger radius/area there, no further 9′ expansion), increase PEF and slightly dampen water-flux variability (relative to apo-L9′T), and nudge twist/tilts in the activation-like direction, consistent with stabilizing a more ordered open-like ensemble rather than creating an alternative opening mechanism. NAMs behave distinctly: BIC (orthosteric antagonist) re-constricts 9′/-2′, reduces hydration/flux, increases twist, and shifts ECD-TMD coupling toward a closed-like quadrant. In contrast, PTX (pore blocker) results in near-zero flux and little quaternary shift. Notably, PTX can make the geometric minimum radius appear large (and PEF high) because the lodged ligand increases the apparent cross-section; this reflects physical occlusion of the pore rather than true permeation accessibility.

The bicuculline-induced, bottom-to-top closure in L9′T includes a distinct asymmetric M2 collapse at -2′, three helices converge while two displace laterally (**Figure 11**), which precedes/quasi-coincides with the -2′-first narrowing and propagates to 9′/20′ with a modest twist increase (Figure 10). Notably, time-resolved cryo-EM has visualized short-lived asymmetric intermediates (closed, open, and desensitized) across β3, α1β3, α1β3γ2 GABA_A_Rs within ≤10 ms of agonist exposure, marked by subunit-specific M2 translations/rotations, branched or linear gating paths, and transient lipid access/occlusion in the pore and interfaces. These data provide an experimental framework consistent with our asymmetric TMD collapse and the staged -2′ → 9′/20′ closure inferred from radii, hydration, flux, and coupling metrics.

Our findings also agree with emerging evidence that gating involves multiple structural drivers beyond pore hydration. For example, β-subunit M2-M3 backbone interactions can stabilize the closed state and weaken during activation, contributing measurably to the energetic cost of opening^57^. Similarly, computational models of native receptors predict open-state ensembles with expanded gates and rearrangements at the αβ interface, consistent with the quaternary motions and hydration signatures we report^58^. Taken together with established evidence for hydrophobic gating and for ECD-TMD twist/tilt coupling across pLGICs^46,47^, these observations support a unified view in which local chemistry at the 9’ hydrophobic gate, backbone flexibility in coupling loops, and interfacial contacts collectively define the gating coordinate. This view is also compatible with the pore-occluding behavior of picrotoxin (PTX) observed structurally and functionally.

Although L9′T/S variants are not clinically prevalent, recent human genetic findings have identified pathogenic variants at the same 9’ position in the *GABRB3* gene (β3 subunit), including L9’P and L9’R, associated with gain-of-function phenotypes manifesting as early-onset epilepsy within the first month of life, movement disorders, and Lennox–Gastaut syndrome^59^. These substitutions, like threonine and serine, are less bulky or polar and disrupt hydrophobic gating. In contrast, the hydrophobic L9′M variant did not elicit significant functional changes, reinforcing the link between polar substitutions at the 9′ gate and spontaneous channel activation^59^.

Experimental mutagenesis and electrophysiology indicate that introducing L9’T into a single subunit (α, β, or γ) is often sufficient to yield a spontaneously active receptor, with substantial basal unliganded *P*_0_ (e.g., α1(L9’T) or γ2(L9’T) ∼0.3-0.4 and a high *P*_0_ for β3(L9′T) ∼0.8) (see **Figure 1 and S1**)^30^. In silico, single- or double-subunit substitutions produce limited pore expansion and partial barrier reductions, whereas only the all-subunit L9′T/S configuration consistently stabilizes open-like states. The apparent discrepancy likely reflects timescale and context: microsecond atomistic simulations versus millisecond-to-second experimental observations, together with cell-level and subunit-specific factors that can amplify gating in the oocyte system. Consistently, single- and double-subunit simulation trajectories reveal how partial gate hydration redistributes energetics without fully stabilizing the open ensemble: single substitutions typically maintain a constricted 9’ (<2 Å) and only partially lower PMF barriers, consistent with smaller hydration increases, while double substitutions show intermediate dilation (PEFs ∼10-21%) and steady water flux with occasional conductance-relevant openings (see Supporting Information). Functionally, the elevated basal *P*_0_ in experimental recordings, especially the β3(L9′T) case, despite limited pore opening on microsecond scales in MD, underscores the value of integrating electrophysiology with simulations and cautions that fast wetting/rare openings may be under sampled in unbiased trajectories. Together with all-subunit results, these observations support a graded view in which increasing 9’ hydrophilicity moves the receptor along the gating coordinate from partially wetted, energetically softened pores to stable open-like ensembles.

These results have practical implications for structural biology and pharmacology. First, they provide open-state templates that better reflect efficacy-linked geometries, hydrated 9’ and flexible -2’ for structure-guided docking of PAMs and for designing ligands that tune hydration or interface contacts rather than simply occupancy. Second, the bicuculline sequence highlights structural elements where orthosteric binding destabilizes ECD-TMD coupling and increases twist, suggesting state-dependent antagonism that could be leveraged to reduce desensitization bias. Third, because the observed motions (untwisting of ECD, tilt of M2 helices) and interfacial tightening mirror features reported across pLGICs, these principles should generalize beyond GABA_A_ receptors. Taken together, our data imply that opening may proceed as the reverse of the bicuculline sequence, rapid 9′ wetting followed by -2′ relief under tightened ECD-TMD coupling, and that PAMs primarily tune the residual -2′ constriction.

In summary, hydrating the 9’ gate with L9′T or L9′S mutations ties local chemistry to global activation, while subtle backbone and interface rearrangements reported in recent studies provide complementary native drivers for open-state stabilization. Diazepam and ganaxolone stabilize open-like states.

Picrotoxin blocks conduction with minimal structural change, while bicuculline induces a bottom-to-top, stepwise closure that likely mirrors opening in reverse. Together, these insights provide a high-resolution view of GABA_A_ receptor gating and identify actionable structural sites for rationally designing state-selective modulators along the activation pathway.

#### BOX 2. STUDY LIMITATIONS

1. **Mutant-stabilized open-like ensembles.** Our open-like states are stabilized by all-subunit L9′T/L9′S substitutions at the 9’ hydrophobic gate. While this approach exposes the geometry-hydration-energetics-conformational linkage clearly, it may not capture every aspect of native WT ligand-triggered gating in heteropentameric channels.
2. **Sampling window and accelerated dynamics.** GaMD provides efficient barrier crossing on the sub-microsecond to microsecond scale, but millisecond-second gating/desensitization kinetics and rare substates remain beyond direct resolution. In our L9′T/L9′S systems the earliest wetting events at 9’ occur rapidly and are not temporally resolved.
3. **CompEL boundary conditions.** Computational electrophysiology used charge imbalances that yielded ∼0.5-1.1 V transmembrane potential to obtain sufficient permeation statistics in finite trajectories. Such supraphysiological ΔU are standard in MD but can amplify field-sensitive phenomena; therefore, reported current-voltage slopes are best interpreted comparatively and in the context of hydration/PMF trends rather than as exact physiological values.
4. **Force fields and water model dependence.** We used CHARMM36m/CGenFF with TIP3P water. Hydration and hydrophobic gating in nanoconfined pores can depend on electronic polarization; polarizable models often increase wetting relative to fixed-charge models and can alter anion-pore interactions. Conclusions are robust across the observables, but absolute wetting propensities may vary with model choice.
5. **Metric-specific caveat under pore blockers.** PEF (frames with minimum radius > 2.0 Å) is radius-threshold-based and does not account for occupancy. With picrotoxin (PTX) lodged in the pore, the apparent radius (and thus PEF) can be high despite functional block; accordingly, we rely on flux (near-zero) to reflect non-conducive state under PTX.
6. **Membrane composition.** Simulations used a simplified POPC:cholesterol bilayer. Anionic lipids (e.g., PIP2) are known to bind α1 subunits in α1β3γ2 and may influence gating and coupling; these interactions are not explicitly modeled here.
7. **Stoichiometry/subtype generalization.** Results presented here pertain to α1β3γ2. Structural and pharmacological features can differ across GABAA subtypes, so quantitative aspects may not generalize without qualification, even if qualitative gating motifs are conserved.

## CONCLUSIONS

This work provides an atomistic, internally consistent view of open-like ensembles in the heteropentameric α1β3γ2 GABA_A_ receptor, obtained by stabilizing the 9′ hydrophobic gate with L9′T/L9′S substitutions. Across geometry, hydration, energetics, conductance, and quaternary motions, the mutant-stabilized open-like states show a unified signature: pore expansion at 9′/20′ with a residual -2′ bottleneck, increased wetting, reduced PMF barriers at 9′ and -2′, ohmic anion conductance, and activation-like conformational changes (reduced twist, outward M2 tilt, modest ECD compaction, C-loop closure). Together, these findings support a framework in which hydrophilic destabilization of the 9′ gate lowers the energetic cost of permeation while biasing the activation-linked quaternary rearrangements characteristic of pLGICs.

By probing positive (diazepam, ganaxolone) and negative (bicuculline, picrotoxin) modulators on this open-like background, we further show that PAMs primarily tune the residual -2′ constriction, whereas bicuculline drives a bottom-to-top, stepwise closure and picrotoxin occludes the pore with minimal global reorientation. These ligand-dependent responses, together with the energy profiles for chloride permeation, establish a tractable structural platform for interrogating state-dependent modulation and for structure-based design of compounds that stabilize specific points along the GABA_A_ receptor activation pathway.

## METHODS

### In vitro Studies

#### Mutagenesis and in vitro transcription

DNA for wild-type and mutant GABA_A_R human α1, β3, and γ2 subunits was subcloned in the pUNIV vector^60^. Mutations were introduced by GenScript and confirmed by sequencing of the entire subunit.

Complementary RNA (cRNA) for each construct was generated (mMessage mMachine T7, Ambion) and quantified (Qubit, ThermoFisher Scientific) prior to injection in *Xenopus laevis* oocytes.

#### Two-Electrode Voltage Clamp (TEVC) recording in oocytes

Defolliculated *Xenopus laevis* oocytes were obtained from Ecocyte. Oocytes were injected with 12 ng of total cRNA for α1, β3, and γ2 subunits (wild-type or mutants) in a 1:1:10 ratio (Nanoject, Drummond Scientific). Oocytes were incubated in a sterile incubation solution (88 mM NaCl, 1mM KCl, 2.4 mM NaHCO_3_, 19 mM HEPES, 0.82 mM MgSO_4_, 0.33 mM Ca(NO_3_)_2_, 0.91 mM CaCl_2_, 10,000 units/L penicillin, 50 mg/L gentamicin, 90 mg/L theophylline, and 220 mg/L sodium pyruvate, pH=7.5) at 16 °C. Currents from expressed channels 1-2 days post-injection were recorded in two-electrode voltage clamp (Oocyte Clamp OC-725C, Warner Instruments) and digitized using a PowerLab 4/30 system (ADInstruments). Data was obtained from at least two different batches of oocytes for each experimental group.

Oocytes were held at -70 mV and perfused continuously (2 mL/min) with ND96 buffer (96 mM NaCl, 2 mM KCl, 1 mM CaCl_2_, 1 mM MgCl_2_, 5 mM HEPES, pH 7.5) or ND96 buffer containing picrotoxin (PTX) or GABA. PTX was diluted from a 0.5 M stock solution in DMSO. GABA was diluted from 1 M stock solution in water. The recording protocol was as follows: a 10 s pulse of PTX was followed by a series of 20-40 s pulses of increasing concentrations of GABA, and a final 10 s pulse of PTX. Pulses were sufficiently long to resolve the peak response and inter-pulse intervals were 5–15 min to allow washout with buffer and currents to return to baseline. Representative traces are shown in **Figure X**. Current traces were analyzed with custom scripts in MATLAB (Mathworks). GABA concentration-response curves (CRCs) were fit with the Hill equation:

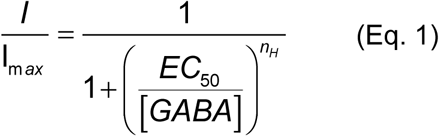

where *I* is the magnitude of the ligand-elicited current, [*ligand*] is GABA concentration, *EC*_50_ is the concentration eliciting a half-maximal response, and *n*_*H*_ is the Hill slope.

Basal unliganded open probability (*P*_*o*_) was calculated from the ratio of PTX-sensitive to total current amplitude (*I*_*PTX*_⁄*I*_*Total*_, see **Figure S1**). The total current amplitude was obtained by adding the spontaneous (PTX-sensitive) current to the maximal current obtained when GABA was applied. This is approximately equivalent to going from fully closed receptors (blocked by PTX) to fully activated (maximal response to GABA). Rundown correction for spontaneously active mutants was as described previously^37^.

### Statistical analysis

Summary data was analyzed using Prism 10 (GraphPad). Symbols and error bars are mean ± SEM, and plots of individual values show median and interquartile intervals. To compare multiple conditions, we applied One-way Brown-Forsythe ANOVA (as the standard deviations were not equal among groups) followed by Dunnett’s T3 multiple comparisons test.

### In silico studies

#### Protein and ligand structure preparation

The cryo-EM structure of heteropentameric α1β3γ2 GABA_A_ receptor in complex with the antagonist Ro15-4513 (PDB ID: 7QNE, 2.70Å)^13^ was used as the starting model; all co-purified ligands were removed prior to simulation. Protein structure preparation in MOE^61^ (QuickPrep) assigned protonation states at pH 7.4, added hydrogens, relieved local clashes by minimization, and preserved the original backbone geometry. In-silico mutagenesis introduced L9’T or L9’S at the 9’ (M2) positions in the indicated subunits (all-subunit, single or double). In 7QNE numbering, 9’ corresponds to α1:L264, β3:L259, γ2: L274 (see **Figure 2C**).

For ligand-bound simulations, ligand coordinates were taken from experimental structures where available (DZP at the α1/γ2 site (PDB ID: 6HUP); BIC at β/α orthosteric sites (PDB ID: 6HUK); PTX within the pore (PDB ID: 6HUJ)^15^ or from PubChem (GAN) and prepared in MOE. Ligand protonation states were assigned at pH 7.4. CHARMM General Force Field (CGenFF)^62^ parameters and charges were generated with CHARMM-GUI Ligand Reader ^63–65^ and penalty scores were inspected to confirm acceptable analog coverage. Docking poses (MOE) were selected to match experimental binding modes. CHARMM36m/CGenFF parameter and topology files^66^ were produced in engine-specific formats for AMBER (GaMD/SMD/US) and for GROMACS (CompEL) without manual alteration. Initial receptor and ligand coordinates are provided in SI (Dataset uploaded to Zenodo).

#### Gaussian-Accelerated MD Simulations

Receptors (apo and ligand-bound) were oriented in the membrane using the OPM server^67^ and embedded in a model bilayer consisting of 16:0/18:1 phosphatidylcholine (POPC) and cholesterol in 10:1 ratio (∼135-140 POPC and 13-15 cholesterol per leaflet)^68^, solvated with TIP3P (water padding up to 23 Å on both sides of the bilayer), and neutralized at 0.15 M NaCl using CHARMM-GUI^63–65^.

Following energy minimization and staged equilibration (following a six-step CHARMM-GUI equilibration protocol) at 310K temperature and 1 bar pressure conditions (2-fs time step, LINCS/SETTLE hydrogen constraints, PME electrostatics^69^, and 12 Å cut-off) each system underwent 20 ns unbiased MD, 65 ns GaMD equilibration, and 500 ns GaMD production per replicate. Three independent GaMD replicates were run per system from distinct equilibrated snapshots (distinct seeds). GaMD used a dual-boost scheme (dihedral + total potential) protocol; boost statistics and σ0 settings are available in log files (Zenodo). Reweighting employed a second-order cumulant expansion^70^. All GaMD runs were performed in AMBER 2021^71^. Further analyses of the simulation trajectory were implemented using the Amber cpptraj package, and visualization was carried out using VMD and PyMOL^72–74^.

#### Steered MD and Umbrella Sampling

We performed steered MD and umbrella sampling (US) simulations to obtain the free energy profile of chloride ion permeation through the GABA_A_ receptor transmembrane channel pore^75^. All the simulations were carried out using the Amber 2021 package. The initial structures for steered MD were obtained from well-equilibrated snapshots after the GaMD simulations. A single Cl^−^ ion was placed in the extracellular vestibule ∼50 Å above the -2’ desensitization gate (measured along the channel axis) and steered along the pore axis (z) from ECD through the pore to the intracellular bulk at 1 Å/ns using a harmonic spring constant k = 2.5 kcal.mol^−1^.Å^−2^ (1 fs time step). We then extracted ∼60 windows at 1 Å spacing along z and performed 20 ns restrained sampling per window with a positional restraint of 5 kcal.mol^−1^.Å^−2^ on the ion’s z-projection. The z-axis reaction coordinate follows the channel symmetry axis used for pore geometry and aligns with the axial positions of the 9’ and -2’ gates^76^. Simulation of all 60 windows resulted in a cumulative simulation time of ∼1.2 µs for each of the studied GABA_A_ receptor systems. The obtained probability distributions were reweighted using the weighted histogram analysis method (WHAM) to derive an unbiased Potential Mean Force (PMF) ^41^. Barrier heights reported in the Results are means across replicates, and uncertainties reflect bootstrap variability (See **SI Table S2**).

#### Computational Electrophysiology

Computational electrophysiology (CompEL) was performed in GROMACS using the double-bilayer systems generated by duplicating equilibrated GaMD snapshots along z. We utilized the protocol proposed to generate a transmembrane potential by maintaining a charge imbalance Δq across the membrane, which leads to a differential potential, ΔU, based on the membrane’s capacitance^42,43^.

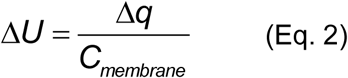

Snapshots of the GABA_A_ receptor system with all-subunit L9′T and L9′S were used as starting structures (Compartment A). The structures were subjected to further equilibration runs as previously described in the GaMD section. A duplicate copy (Compartment B) of the equilibrated simulation box along the z-axis of the membrane was generated using the Gromacs *gmx editconf* command. Also, the index file was duplicated with the (gmx make_ndx -twin). The compEL protocol was enabled by adding the keyword, *swapcoords = Z,* in the *.mdp* input file. We defined the pore position in compartments A and B by a cylinder of 1.0 nm radius at the M2 9’ position, which extends by 1.5 nm in the direction of the ECD and downwards to the intracellular region. Different transmembrane potentials were created by varying the chloride ion concentrations (Δq = 2, 4, 6, 8, 10 e) in both compartments while the concentration of sodium ions kept constant. The ion *swap frequency* was set to 100, *coupl-steps* to 10, and the *threshold* to 1. The simulations were thereafter subjected to a production run of 100 ns^77^. The potential difference due to each charge imbalance (Δq) was calculated using the *gmx potential* command^78^. Also, details of ion movement between the compartments and time were recorded in *swapions.xvg* file.

Based on the number of permeation events, transmembrane potential, and time, we computed the channel conductance using the equation:

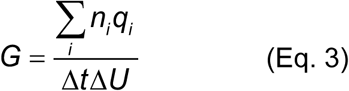

In Eq 3, G is the computed conductance, which is obtained from the number of permeations (n), due to a charge imbalance (q) that generated a potential difference (U) during a time interval (t).

#### Channel Pore Radius

Pore radius profiles and surfaces were computed by the HOLE program along the channel axis on full production trajectories via the MDAnalysis-Hole Interface^79,80^. HOLE defines a series of points along the pore where a sphere can be placed without overlapping any atoms based on their van der Waals radii. From the output files, gate positions (20’, 9’, -2’) were identified from residue z-coordinates. Time-averaged profiles and per-frame minima were used to visualize constrictions and to compute pore expansion frequency (PEF = fraction of frames with minimum radius > 2.0 Å).

#### Cross-sectional Pore Area from M2 Cα Pentagons

To quantify backbone-level widening of the channel, we computed a cross-sectional area at specific axial positions (20′, 9′, 6′, 2′, −2′,) along the pore in VMD using an in-house Tcl script. For each simulation frame, we identified the five pore-lining M2 helices and extracted the Cα coordinates of the residue whose z-coordinates match the target axial position within a small tolerance (±0.5 Å). We then calculated the five nearest-neighbor Cα-Cα distances around the ring (i.e., β→α→β→α→γ order matching the pentameric arrangement). The mean nearest-neighbor distance *ā* was used as the side length of an ideal regular pentagon that approximate the pore cross-sectional area at that position. The area *A* of the regular pentagon was computed using Eq. 4 given below.

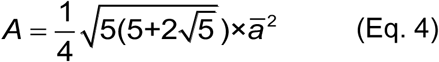

Where *ā* is the average of the five nearest-neighbor distances for that frame and position. This Cα-based area metric emphasizes backbone separation among M2 helices and is therefore complementary to HOLE-based pore radii, which are more sensitive to sidechain rotamers and local packing. For each system, we report time-averaged areas and distributions over the trajectory. Gate positions (20′, 9′, −2′) follow the standard pLGIC numbering; the 6′ and 2′ planes are provided in SI to illustrate intermediate axial changes. were computed for each frame of the entire trajectory in VMD using an in-house Tcl script.

#### Handling asymmetric TMD collapse

In bicuculline-bound trajectories, asymmetric M2 displacements (three helices convergent, two displaced laterally) can inflate the five-helix regular-pentagon area despite pore closure. To obtain an area readout consistent with radius/flux, we computed a three-helix cross-section from the convergent side only: for each frame/plane we selected the three nearest M2 Cαs (one per converging subunit), averaged their pairwise distances, and reported the area of the isosceles-triangle envelope (converted to an equivalent pentagon-normalized area for comparability across panels; see Figure 11). This adjustment preserves backbone-spacing sensitivity while preventing artificial area inflation under asymmetric collapse. The asymmetric-state rationale is supported by time-resolved cryo-EM evidence for asymmetric gating intermediates in GABA_A_Rs.

#### Pore hydration and water flux

We quantified hydration near the hydrophobic gate by counting water molecules within ±7 Å of the 9’ gate along the pore axis (i.e., a 14 Å slab centered at 9’, spanning 7 Å above and 7 Å below). This window captures the gate region with tolerance for local backbone/side-chain motions. Sensitivity tests using ±6-8 Å produced the same qualitative conclusions. We quantified the number of water molecules passing through the channel pore (flux) using the Flux module in Wordom^81^. Water flux was computed as direction-agnostic counts of complete transits between the extracellular 20’ and intracellular -2’ sides per unit time (10 ps); cumulative flux was obtained by summation over the trajectory.

#### Global twist, tilts (polar and azimuthal), and orthosteric distance

The global conformational dynamics of GABA_A_R across the various systems were tracked over time using two structural observables, twist (τ) and tilt angles (θ), analyzed with Wordom software^81,82^. It is well established that ion-channel activation involves radial reorientation, outward tilting of the M2 helices, and tangential movements. These angles capture the global conformational changes associated with GABA_A_R gating and ungating mechanisms. Twist angle (τ) was defined as the average angle between the projections of the center of mass (COM) vectors of the extracellular domain (ECD) and transmembrane domain (TMD) regions of individual subunits onto the plane perpendicular to the receptor’s symmetry axis. To minimize noise, only the Cα atoms of the subunits were included in τ calculations. The symmetry axis direction was determined by performing a 3D least-squares fitting on the Cartesian coordinates of the Cα atoms. The following residue ranges were used: α-subunit ECD 13-223, TMD 224-418; β-subunit: ECD 8-217, TMD 218-447; γ-subunit ECD 26-233, TMD 234-436.

Similarly, the tilt angle of the channel’s pore-lining M2 helices was derived by measuring the angle between each helix axis and the receptor axis and then averaging these values across all subunits. The helix and receptor axes were identified through 3D least-squares fitting of the Cα atom coordinates. The following M2 residue ranges were used to calculate the tilt angle θ: residues 246-271 for the β3 subunit, 251-276 for the α1 subunit, and 262-285 for the γ2 subunit. The orthosteric C-loop proxy was measured as β3:Y205-α1:R120 inter-residue distance using the oxygen atom of the sidechain -OH group of Y205 and the amino nitrogen of the sidechain guanidine of R120.

##### Reproducibility and uncertainties

Unless stated otherwise, each system was stimulated in triplicate from distinct equilibrated starting structures with unique random seeds. GaMD boost statistics and reweighting diagnostics are available in log files (Zenodo). WHAM bootstrap uncertainties are reported in Table S2.

## FUNDING/ACKNOWLEDGEMENTS

This research was supported by NIH/NIGMS grants R01GM148591 to MPG-O and R01GM137022 to SN.

## AUTHOR CONTRIBUTIONS

AD, SN, and MPG-O conceived and designed this work. AD carried out and SN supervised all simulations and analysis thereof. CMB carried out and MPG-O supervised all TEVC recordings and analysis thereof. AD, CMB, MPG-O, and SN wrote the manuscript.

## ADDITIONAL INFORMATION

**Supplementary information.** A pdf document “Supporting_Information_Diyaolu.pdf” is attached.

**Supporting Movie: M1** – Energetics of chloride permeation through the pore determined using steered MD and umbrella sampling

**Supporting Movie: M2** – Chloride permeation through the pore during CompEL

**Supporting Movie: M3** – Asymmetric collapse of M2 helices upon bicuculline binding

## DATA AVAILABILITY

All input structural data and parameters for unbiased MD, GaMD, SMD, umbrella sampling, computational electrophysiology simulations, and sample trajectories are available at 10.5281/zenodo.18447712

