## Supporting Information for "Open-State Dynamics and Allosteric Modulation of the α1β3γ2 GABA_A_ Receptor Stabilized by L9′T/S Substitutions"

**Table S1. Electrophysiology data of single, double, and all-subunit L9'T and L9'S α1β2/3γ2 $GABA_A$ receptors from the literature.**

| $GABA_A$ receptor subtype | Mutation | Mutant $EC_{50}$ (μM) | Fold shift vs Wildtype | References |
|---|---|---|---|---|
| α1β3γ2 | γ2(L274S) | 0.29 | ~37 | (Bianchi *et al.*, 2001)[1] |
| α1β1γ2 | α1(L264T) | 5.0 | ~2 | (Dalziel *et al.*, 2000)[2] |
| α1β2γ2 | α1(L264T) | $0.30 \pm 0.02$ | ~83 | (Scheller & Forman 2002, Nors et al. 2021, Nors et al. 2024)[3-5] |
| α1β2γ2 | γ2(L274S) | $1.04 \pm 0.05$ | ~44 | (Chang & Weiss, 1999)[6] |
| α1β2γ2 | α1(L264S) | $0.22 \pm 0.02$ | ~208 | |
| α1β2γ2 | β2(L259S) | $0.052 \pm 0.005$ | ~881 | |
| α1β2γ2 | α1(L264S) γ2(L274S) | $0.095 \pm 0.003$ | ~482 | |
| α1β2γ2 | β2(L259S) γ2(L274S) | $0.07 \pm 0.01$ | ~664 | |
| α1β2γ2 | α1(L264S) β2(L259S) | $0.038 \pm 0.007$ | ~1205 | |
| α1β2γ2 | α1(L264S) β2(L259S) γ2(L274S) | $0.066 \pm 0.002$ | ~694 | |

**Table S2: Chloride inflow potential of mean force error estimation**

| | WT | | L9'T | | L9'S | |
|---|---|---|---|---|---|---|
| Window | Mean (kcal/mol) | ±SD (kcal/mol) | Mean (kcal/mol) | ±SD (kcal/mol) | Mean (kcal/mol) | ±SD (kcal/mol) |
| -50.0 | 0.000000 | 0.000000 | 0.000000 | 0.000000 | 0.000000 | 0.000000 |
| -49.0 | 0.787513 | 0.064976 | 0.464062 | 0.087344 | 0.725061 | 0.062225 |
| -48.0 | 0.772946 | 0.061732 | 0.467323 | 0.082220 | 0.630863 | 0.055995 |
| -47.0 | 0.726220 | 0.052317 | 0.442669 | 0.078728 | 0.485064 | 0.067985 |
| -46.0 | 0.641251 | 0.050645 | 0.505462 | 0.084444 | 0.438427 | 0.061096 |
| -45.0 | 0.560008 | 0.046845 | 0.563277 | 0.077133 | 0.334903 | 0.059379 |
| -44.0 | 0.488678 | 0.042721 | 0.610010 | 0.067804 | 0.232630 | 0.062409 |
| -43.0 | 0.364074 | 0.036979 | 0.639530 | 0.069596 | 0.169930 | 0.055653 |
| -42.0 | 0.254437 | 0.028444 | 0.627193 | 0.061180 | 0.080014 | 0.056038 |
| -41.0 | 0.089921 | 0.022103 | 0.648139 | 0.049678 | 0.014791 | 0.055405 |
| -40.0 | 0.000000 | 0.027158 | 0.621102 | 0.048155 | 0.006218 | 0.048075 |
| -39.0 | 0.051719 | 0.031271 | 0.532412 | 0.042690 | 0.000000 | 0.048280 |
| -38.0 | 0.127990 | 0.040382 | 0.447507 | 0.048476 | 0.124600 | 0.048168 |
| -37.0 | 0.282394 | 0.039831 | 0.304184 | 0.047262 | 0.311781 | 0.051882 |
| -36.0 | 0.576793 | 0.039221 | 0.282156 | 0.045267 | 0.301790 | 0.053688 |
| -35.0 | 0.819254 | 0.040725 | 0.209630 | 0.035232 | 0.342290 | 0.047323 |
| -34.0 | 1.543803 | 0.057066 | 0.000000 | 0.038274 | 0.572558 | 0.054650 |
| -33.0 | 2.203648 | 0.069532 | 0.138372 | 0.041505 | 0.582166 | 0.066756 |
| -32.0 | 2.571460 | 0.076218 | 0.311599 | 0.035394 | 0.735623 | 0.058941 |
| -31.0 | 3.125892 | 0.085160 | 0.478551 | 0.052791 | 0.618228 | 0.065138 |
| -30.0 | 3.626434 | 0.081415 | 0.618128 | 0.059843 | 0.672339 | 0.071416 |
| -29.0 | 3.527391 | 0.081709 | 0.583478 | 0.057469 | 0.907695 | 0.072388 |
| -28.0 | 3.506237 | 0.092284 | 0.756891 | 0.062693 | 0.922195 | 0.061984 |
| -27.0 | 4.148379 | 0.109122 | 0.786297 | 0.056737 | 1.128345 | 0.056144 |
| -26.0 | 5.115588 | 0.112340 | 0.857714 | 0.060815 | 1.152821 | 0.053036 |
| -25.0 | 5.888539 | 0.116813 | 0.930283 | 0.071616 | 1.164998 | 0.060073 |
| -24.0 | 6.265865 | 0.118022 | 1.036347 | 0.078634 | 1.369936 | 0.072782 |
| -23.0 | 6.512317 | 0.120674 | 1.015393 | 0.084293 | 1.324272 | 0.083458 |
| -22.0 | 6.826054 | 0.121471 | 0.955037 | 0.094025 | 1.277494 | 0.092676 |
| -21.0 | 7.398250 | 0.130448 | 0.993090 | 0.099407 | 1.313291 | 0.101682 |
| -20.0 | 8.318685 | 0.147314 | 1.247535 | 0.101712 | 1.199815 | 0.103294 |

| | | | | | | |
|---|---|---|---|---|---|---|
| -19.0 | 9.762346 | 0.128203 | 1.422028 | 0.096761 | 1.059062 | 0.104742 |
| -18.0 | 11.763881 | 0.174538 | 1.493423 | 0.099864 | 0.934458 | 0.101157 |
| -17.0 | 13.011714 | 0.187419 | 1.457677 | 0.094627 | 0.898796 | 0.086723 |
| -16.0 | 17.325632 | 0.192296 | 1.436707 | 0.099104 | 0.964088 | 0.077016 |
| -15.0 | 18.436617 | 0.211465 | 1.558902 | 0.106943 | 0.980032 | 0.088808 |
| -14.0 | 19.081897 | 0.212237 | 1.625447 | 0.105034 | 1.005702 | 0.086133 |
| -13.0 | 18.119499 | 0.209917 | 1.473226 | 0.097317 | 1.313184 | 0.085455 |
| -12.0 | 16.510624 | 0.226765 | 1.560859 | 0.100962 | 1.608852 | 0.075329 |
| -11.0 | 15.905184 | 0.223123 | 1.507827 | 0.106089 | 1.763146 | 0.088731 |
| -10.0 | 15.410503 | 0.218708 | 1.672026 | 0.106603 | 2.004767 | 0.091104 |
| -9.0 | 14.222497 | 0.205602 | 1.677951 | 0.111046 | 2.108176 | 0.095911 |
| -8.0 | 14.139212 | 0.205778 | 1.805061 | 0.119514 | 2.194510 | 0.097178 |
| -7.0 | 14.047906 | 0.212289 | 1.940729 | 0.111774 | 2.029765 | 0.108045 |
| -6.0 | 14.305663 | 0.214784 | 2.223460 | 0.126589 | 2.117235 | 0.105868 |
| -5.0 | 13.757523 | 0.217146 | 2.447162 | 0.123346 | 2.343362 | 0.109252 |
| -4.0 | 12.571684 | 0.216471 | 2.535715 | 0.133415 | 2.821981 | 0.112908 |
| -3.0 | 11.998064 | 0.234975 | 2.640566 | 0.139701 | 3.409744 | 0.114705 |
| -2.0 | 11.771169 | 0.235853 | 2.612001 | 0.140311 | 3.772735 | 0.111112 |
| -1.0 | 11.088642 | 0.240918 | 2.559060 | 0.139160 | 3.703875 | 0.115460 |
| 0.0 | 10.101609 | 0.259400 | 2.524517 | 0.143143 | 3.166799 | 0.121372 |
| 1.0 | 9.634788 | 0.263471 | 2.729491 | 0.152351 | 2.870030 | 0.132081 |
| 2.0 | 9.108915 | 0.264163 | 2.667568 | 0.159431 | 2.630183 | 0.133514 |
| 3.0 | 8.262407 | 0.262866 | 3.093355 | 0.168558 | 1.995987 | 0.147498 |
| 4.0 | 7.503622 | 0.261514 | 3.267920 | 0.167314 | 1.104618 | 0.157715 |
| 5.0 | 7.071476 | 0.258849 | 3.042923 | 0.165749 | 0.847123 | 0.152451 |
| 6.0 | 6.851728 | 0.265861 | 2.528268 | 0.169572 | 0.442977 | 0.147890 |
| 7.0 | 6.677323 | 0.261016 | 1.916331 | 0.165699 | 0.546789 | 0.149083 |
| 8.0 | 6.466235 | 0.266841 | 1.060212 | 0.173780 | 0.529539 | 0.159551 |
| 9.0 | 6.406962 | 0.269347 | 0.508437 | 0.181451 | 0.455373 | 0.150570 |

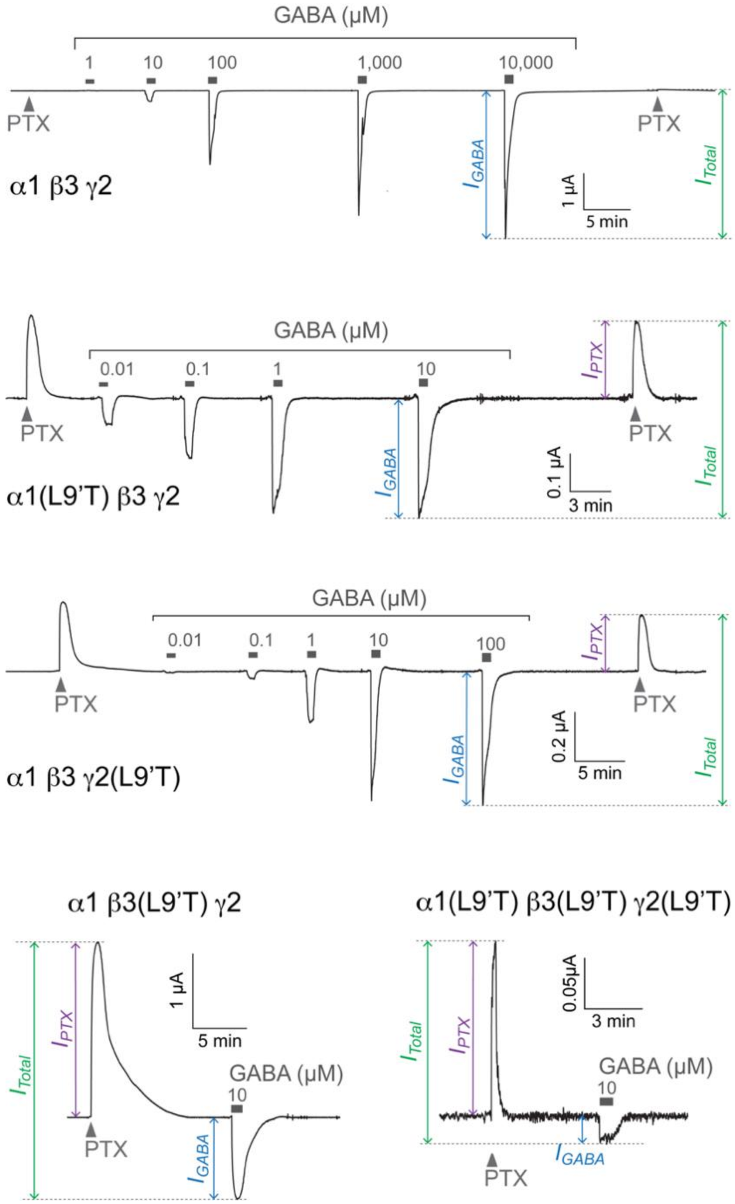


**Figure S1. Representative current traces of responses to brief applications of GABA and PTX in α1β3γ2 $GABA_A$ receptors without and with the gain-of-function L9'T substitution in individual or all subunits.** PTX applications (1 mM, 10 s) are indicated with triangles. GABA applications (20-40 s) are indicated by bars. Downward current in response to GABA reflects opening of the channel pore, whereas upward current in response to the pore blocker PTX reflects block of a spontaneous unliganded standing current conferred by the gain-of-function L9'T substitutions. The total current amplitude ($\boldsymbol{I_{Total}}$) is the sum of the spontaneous ($\boldsymbol{I_{PTX}}$) and the maximal GABA-evoked ($\boldsymbol{I_{GABA}}$) current amplitudes.

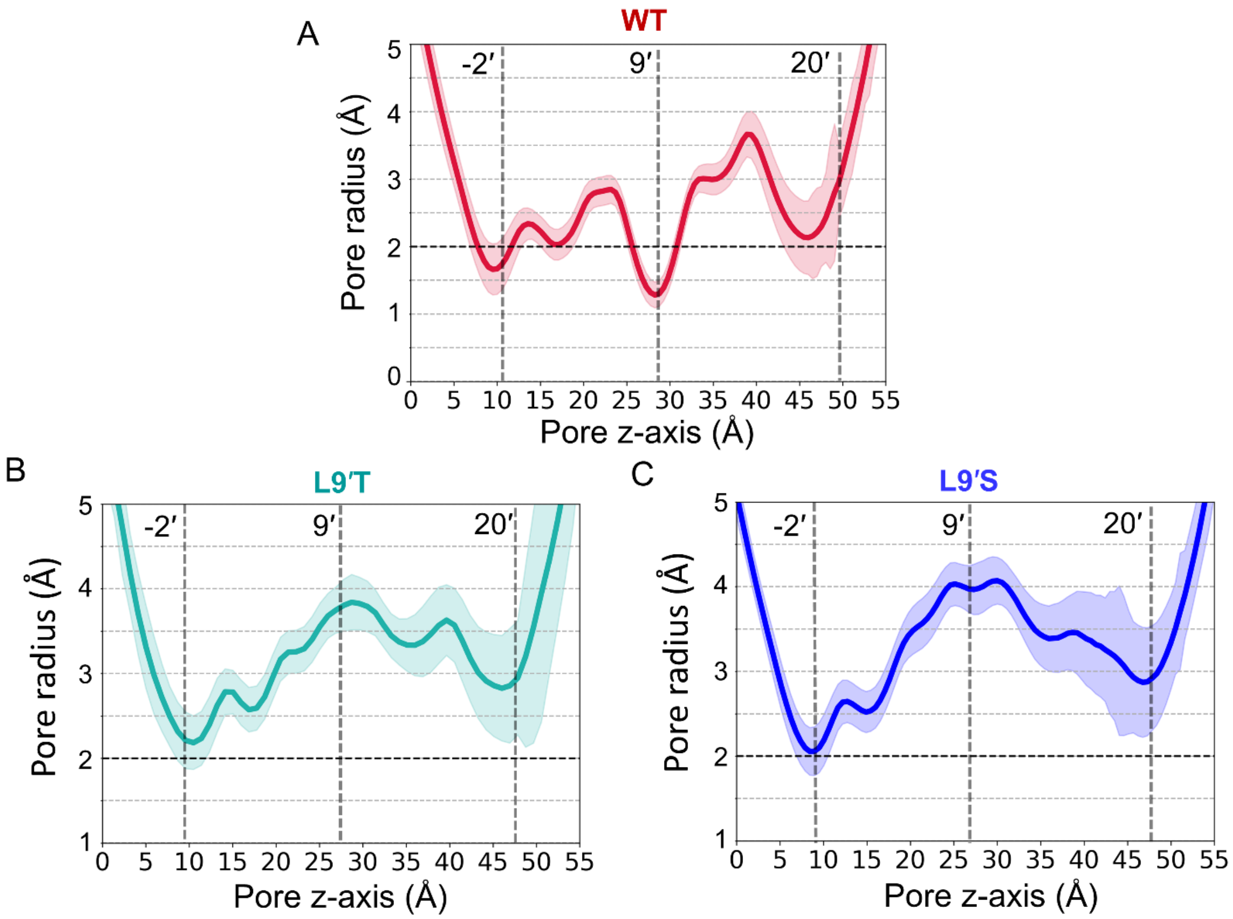


**Figure S2. Pore radius dynamics in WT, all-subunit L9'T and L9'S mutants.** Time-averaged HOLE radius profiles along the channel axis over 500 ns GaMD for WT (A), L9'T (B) and L9'S (C). Mutants show clear expansion at 9' (hydrophobic gate) and 20' (pore entry) relative to WT; -2' widens modestly and intermittently exceeds 2.0 Å. Vertical reference lines mark 20'/9'/-2' gate planes.

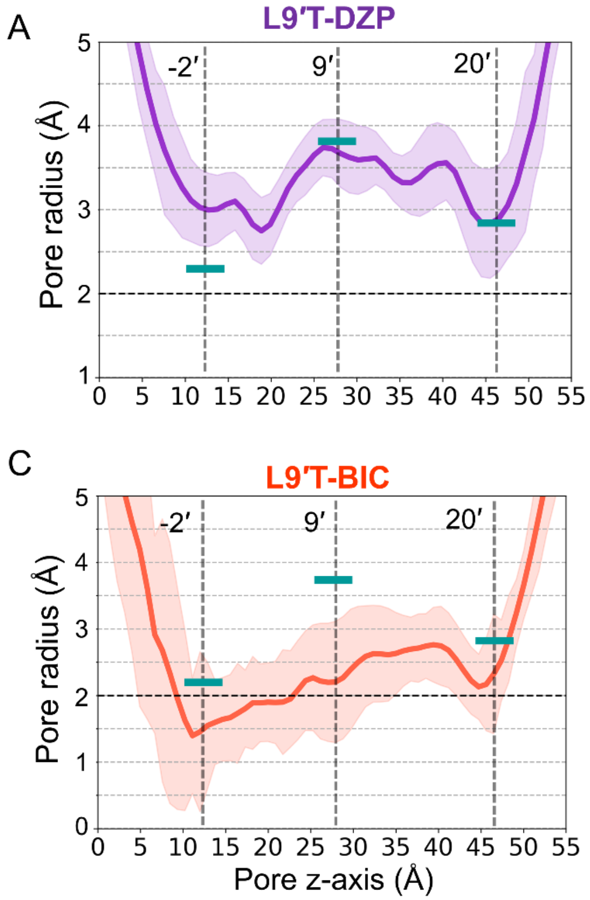


**Figure S3. Pore radius ranges in L9'T with modulators**. Per-frame HOLE radius ranges at -20', 9', and -2' gates for L9'T bound to diazepam (A), ganaxolone (B), bicuculline (C), and picrotoxin (D). Thick lines (light seagreen) indicate the apo- L9'T mean at each gate for reference. These panels summarize how PAMs/NAMs bias gate access relative to the apo open-like background.

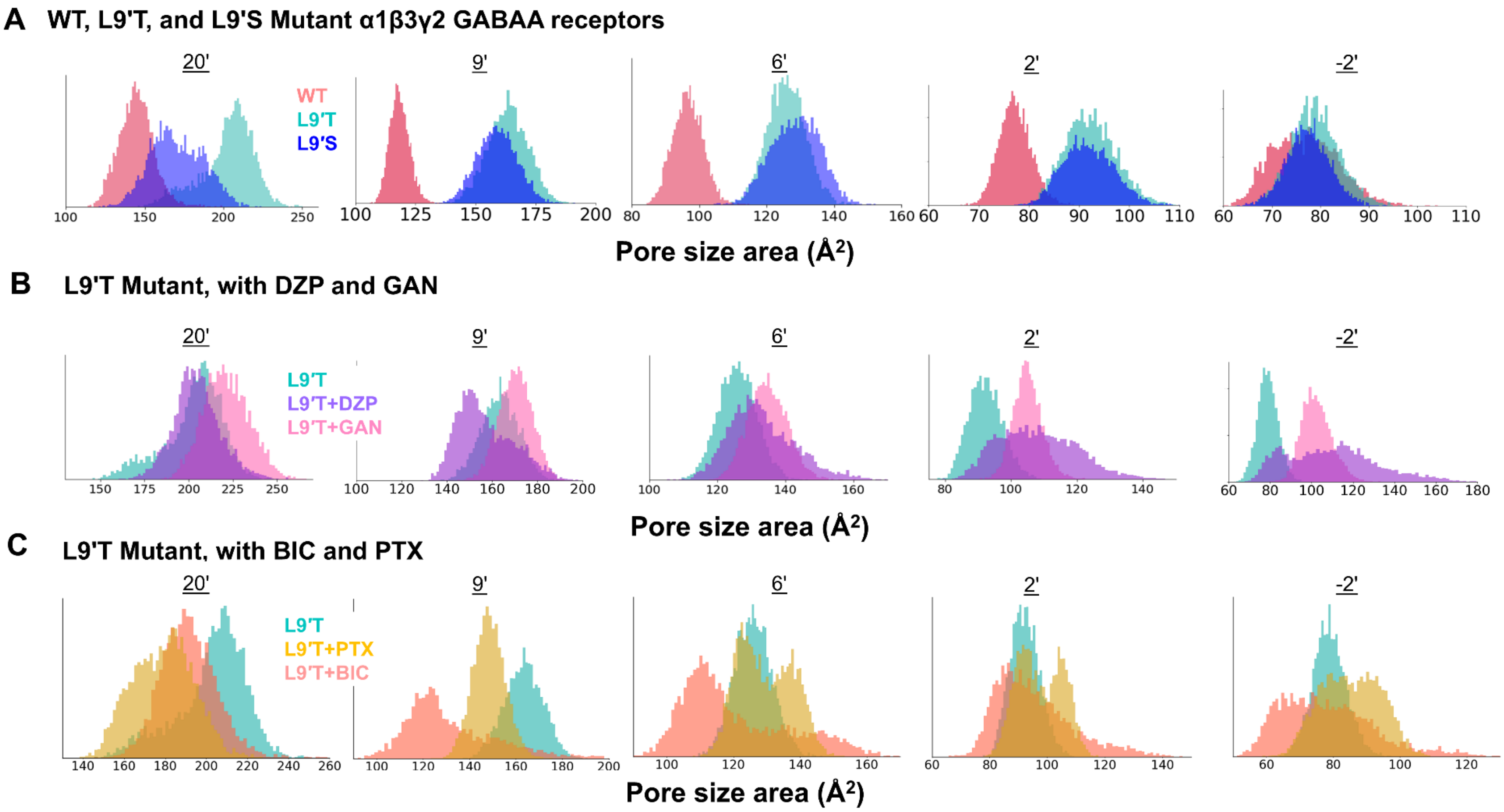


**Figure S4. Cross-sectional pore area distributions at five axial positions in WT, L9'T, L9'S, and modulator-bound $\alpha_1\beta_3\gamma_2$ GABA$_A$ receptors. (A)** Density plots of cross-sectional pore areas at 20', 9', 2', and -2' positions for WT, L9'T, L9'S receptors. (B) Same axial positions for the L9'T receptor in the presence of diazepam (DZP) and ganaxolone (GAN). (C) Same axial positions for the L9'T receptor in the presence of bicuculline (BIC) and picrotoxin (PTX). At each axial position, the pore cross-section was computed from M2 Cα pentagons. For each simulation frame, the five nearest-neighbor M2 Cα-Cα distances were computed, their mean was used as the side length of a regular pentagon, and the area of that pentagon was reported (see Methods). This backbone-based metric reports M2 helix separation and complements HOLE-based pore radius calculations. Each density plot shows the distribution over the full GaMD trajectory for the indicated system.

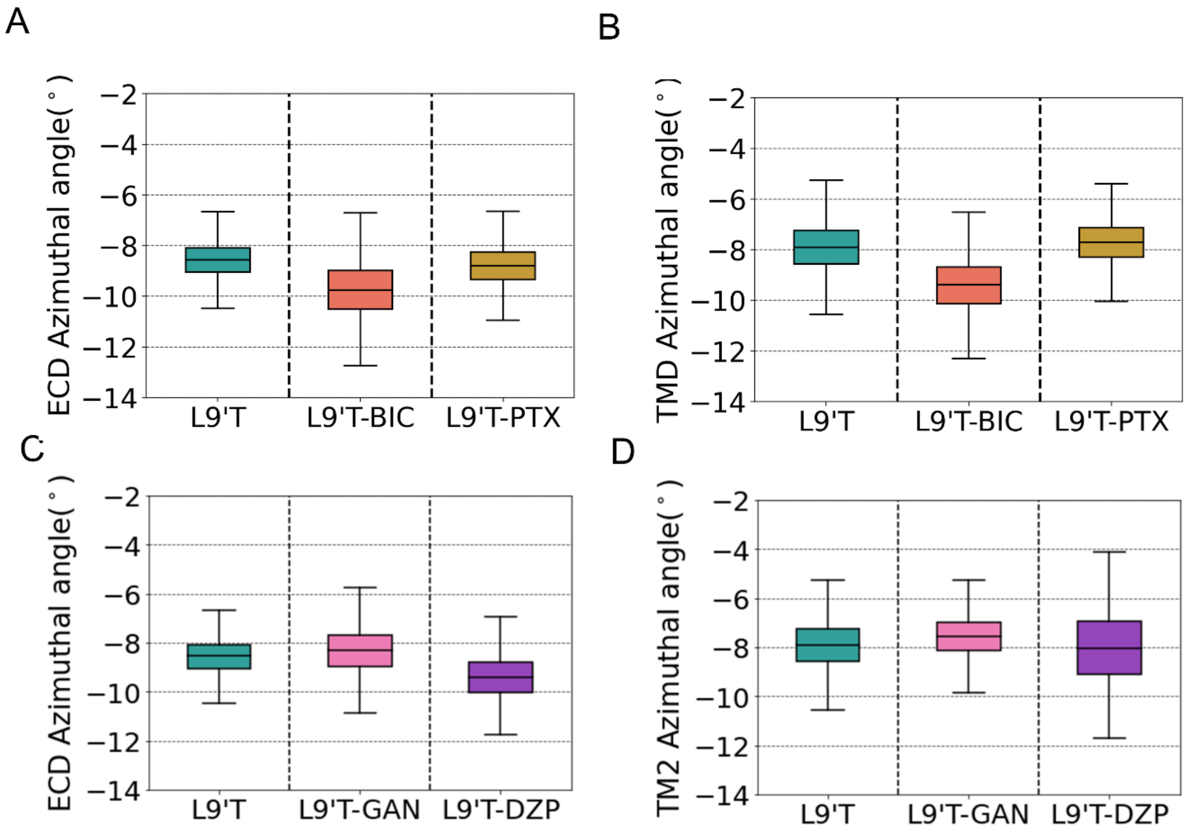


**Figure S5. Azimuthal angles (ECD β-sandwich and M2 helices) in L9'T with modulators.** (A, C) Boxplots showing ECD β-sandwich azimuthal angles in the presence of NAMs (A) and PAMs (C). (B, D) Boxplots showing M2 helix azimuthal angles in the presence of NAMs (B) and PAMs (D). Positive azimuthal denotes clockwise rotation viewed from the ECD (signs as in Methods). Boxes show median/IQR; whiskers span the full sampled range.

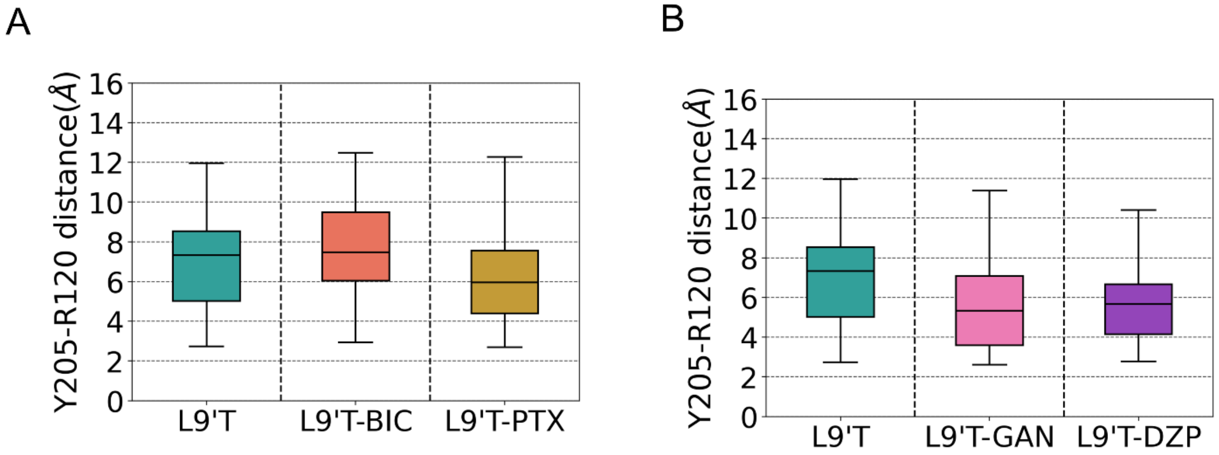


**Figure S6. Orthosteric (site 2) C-loop proxy in L9'T with modulators. (A)** β3:Y205-α1:R120 distance at orthosteric site 2 for L9'T + NAMs bicuculline (BIC) and picrotoxin (PTX) over GaMD trajectories. (B) distances for L9'T + PAMs ganaxolone (GAN) and diazepam (DZP). Smaller values indicate C-loop closure. Atom definitions as in Methods.

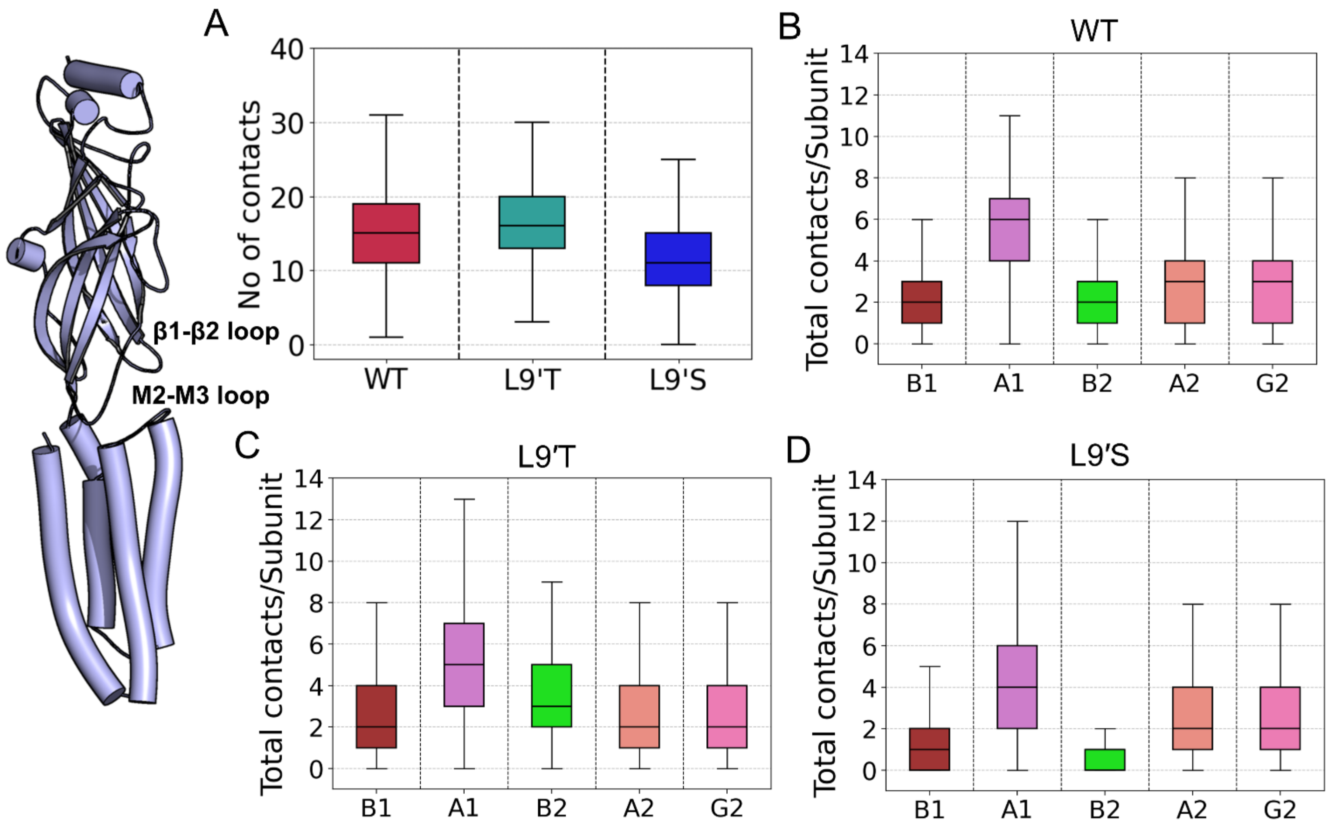


**Figure S7. ECD-TMD loop contacts for WT vs all-subunit mutants.** Number of contacts between TMD M2-M3 and ECD β1-β2 loops in WT and all-subunit L9'T and L9'S: pooled distributions (A) and pre-subunit breakdowns for WT (B), L9'T (C), and L9'S (D). Subunits: A1= α1(1); A2 = α1(2); B1 = β3(1); B2 = β3(2); G = γ2. The residue selection for each subunit is as follows: α1subunit: β1-β2 loop 53-60, M2-M3 loop 276-285; β3 subunit: β1-β2 50-57, M2-M3 271-280; γ2 subunit: β1-β2 65-72, M2M3 286-295.

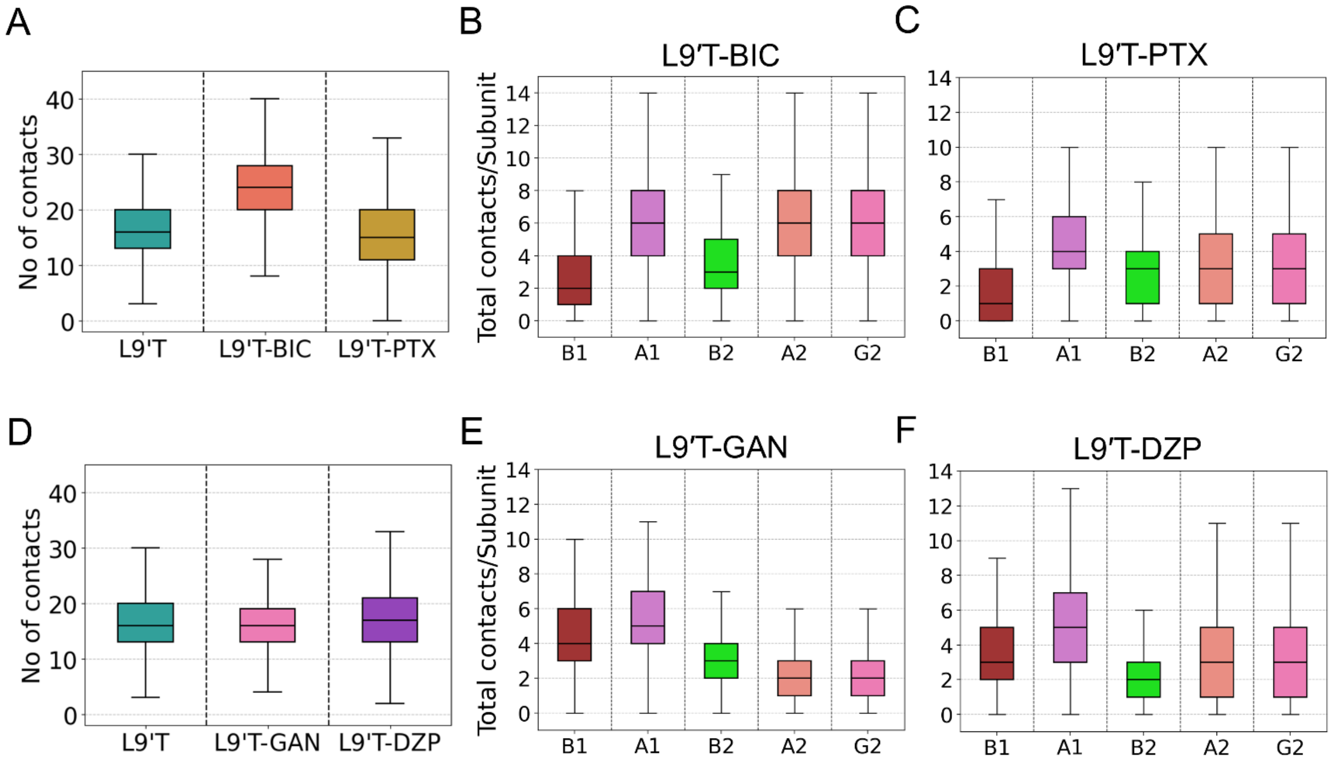


**Figure S8. ECD-TMD loop contacts in all-subunit L9'T bound to modulators.** Boxplots show the distribution of the number of contacts between TMD M2-M3 and ECD β1-β2 loops in all-subunit L9'T with NAMs: pooled distribution (A), per-subunit distribution with apo-L9′T, L9′T + bicuculline (B), and L9'T+picrotoxin (C). Number of contacts in all-subunit L9'T with PAMs: pooled distribution (D), per-subunit distribution with ganaxolone (E) and diazepam (F). Subunits: A1= α1(1); A2 = α1(2); B1 = β3(1); B2 = β3(2); G: γ2. ECD and TMD residue contact definitions as in Methods.

**Single- and Double-Subunit L9'T/L9'S Variants**

To probe how partial 9′ hydrophilic substitution shapes the gating landscape, we performed the same geometric, hydration, energetic, and conformational analyses on single- and double-subunit L9′T/L9′S variants as for the all-subunit mutants. These constructs test whether incremental wetting at the 9′ hydrophobic gate yields graded stabilization of open-like features, and whether limited substitution can reproduce aspects of the all-subunit phenotype within the same simulation and analysis framework. All systems were built and simulated with identical force fields, equilibration, GaMD production, SMD/US protocols, and measurement conventions as the main text (see Methods), ensuring that trends are directly comparable across stoichiometries.

For each variant, we report (i) pore geometry (HOLE radius profiles; per-frame minima; PEF), (ii) backbone-level widening (M2 Cα-pentagon cross-sectional areas at 20′, 9′, 6′, 2′, -2′), (iii) hydration and water flux within ±7 Å of 9′ and direction-agnostic complete transits, (iv) $Cl^-$ permeation energetics (US/WHAM PMFs and barrier heights at 9′ and -2′), and (v) conformational metrics (global twist; M2 and ECD β-sandwich tilts; β3:Y205-α1:R120 as a C-loop proxy). As summarized in the Discussion, single-subunit substitutions typically retain a constricted 9′ (< 2 Å) and partially reduce PMF barriers with lower hydration, whereas selected double-subunit combinations exhibit intermediate dilation (PEFs ~10-21%), steady water flux, and partial barrier lowering, with activation-like tilt/twist changes of smaller magnitude than in the all-subunit L9′T/L9′S background. Detailed distributions and time series for each observable are provided in Figures S9-S27, with system compositions, run lengths, and analysis information in Methods.

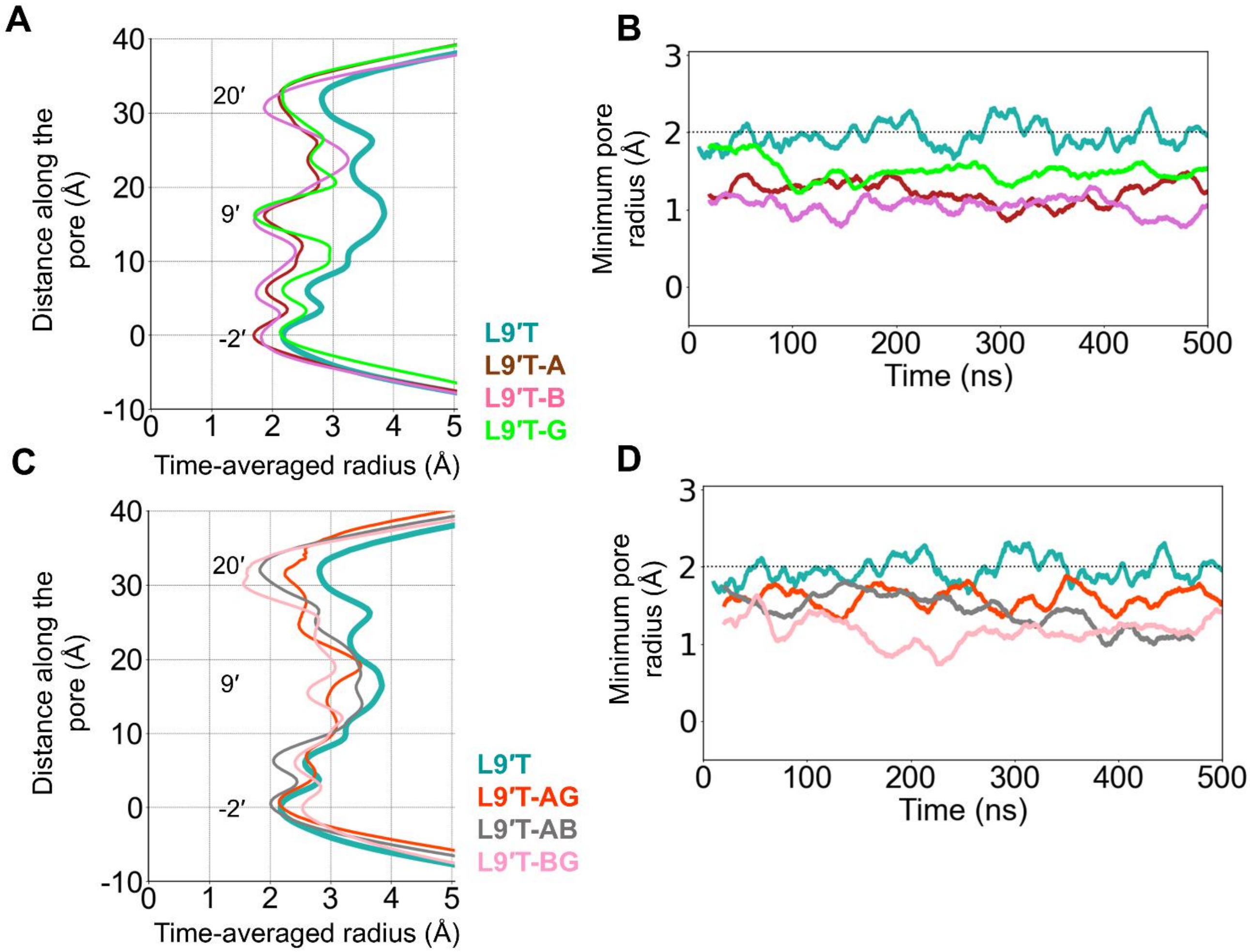


**Figure S9. Pore size dynamics across single- and double subunit L9'T α1β3γ2 $GABA_A$ receptors.** (A, C) Time-averaged pore radius profiles along the z axis for single- (A) and double-subunit (C) L9'T variants over 500 ns GaMD. Gate planes 20', 9', -2' indicated as references. (B, D) Per-frame minimum radius time series for the corresponding systems. Colors and labels match the figure legend. Subunit labels: A = α1; B = β3; G: γ2.

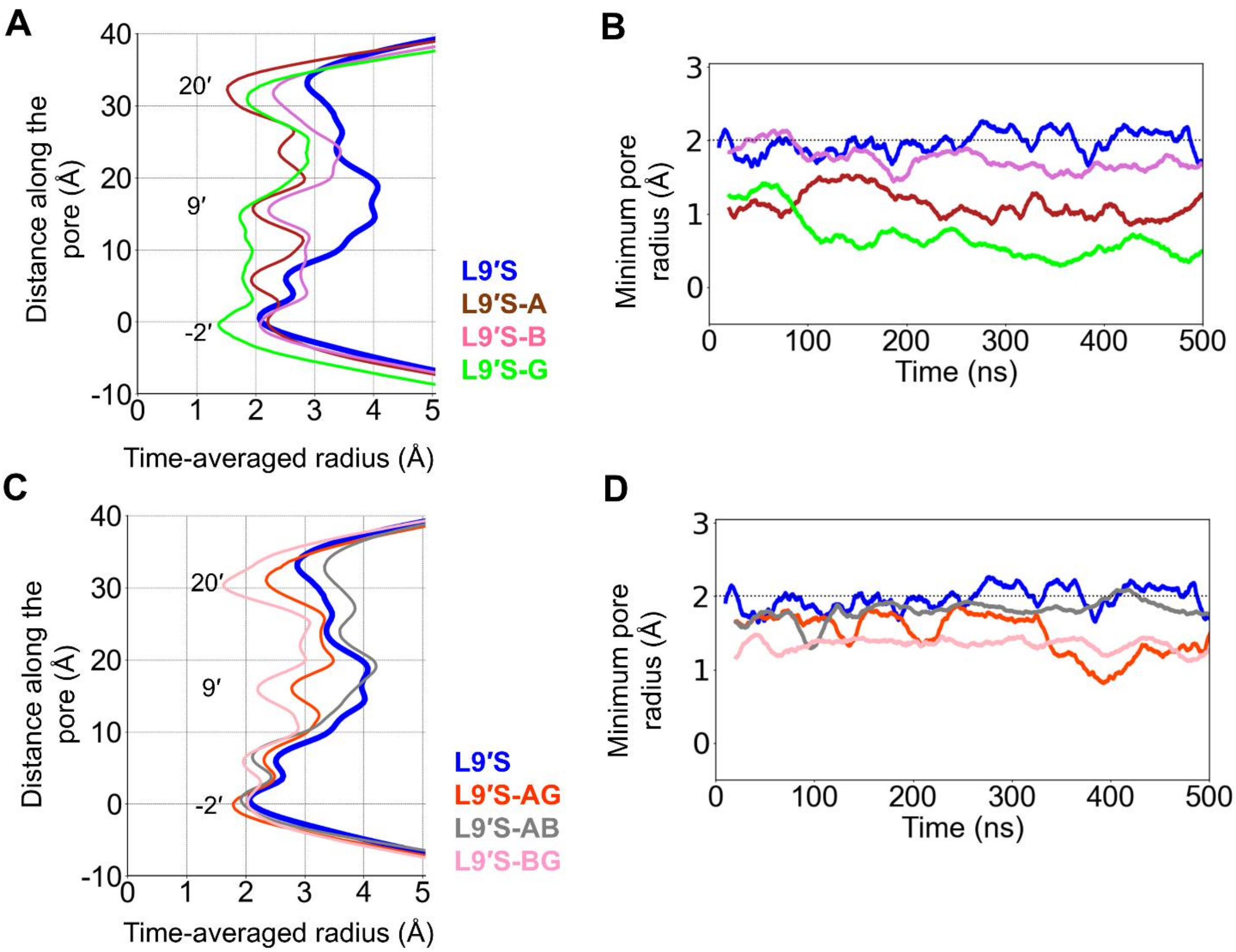


**Figure S10. Pore size dynamics across single- and double subunit L9'S α1β3γ2 GABA$_A$ receptors.** (A, C) Time-averaged pore radius profiles along the z axis for single- (A) and double-subunit (C) L9'S variants over 500 ns GaMD. Gate planes 20', 9', -2' indicated as references. (B, D) Per-frame minimum radius time series for the corresponding systems. Colors and labels match the figure legend. Subunit labels: A = α1; B = β3; G: γ2.

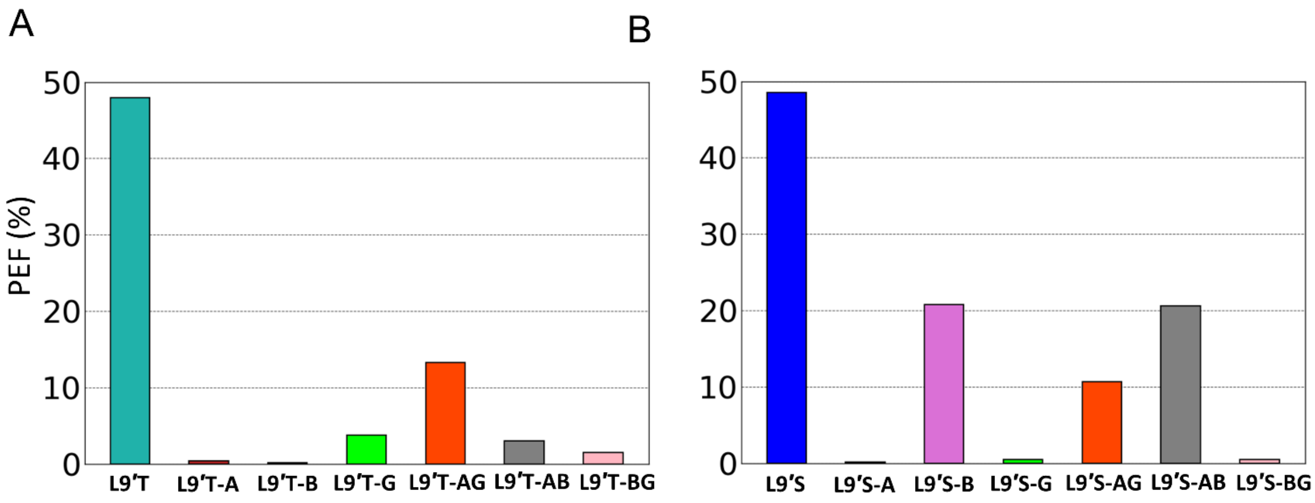


**Figure S11. Pore expansion frequency (PEF) across single- and double-subunit L9'T and L9'S variants.** (A) L9'T and (B) L9'S single- and double-subunit variants. PEF = fraction of simulation frames with minimum pore radius > 2.0 Å over 500 ns GaMD. Subunit codes: A = α1, B = β3, G = γ2. Subunit colors are consistent across panels.

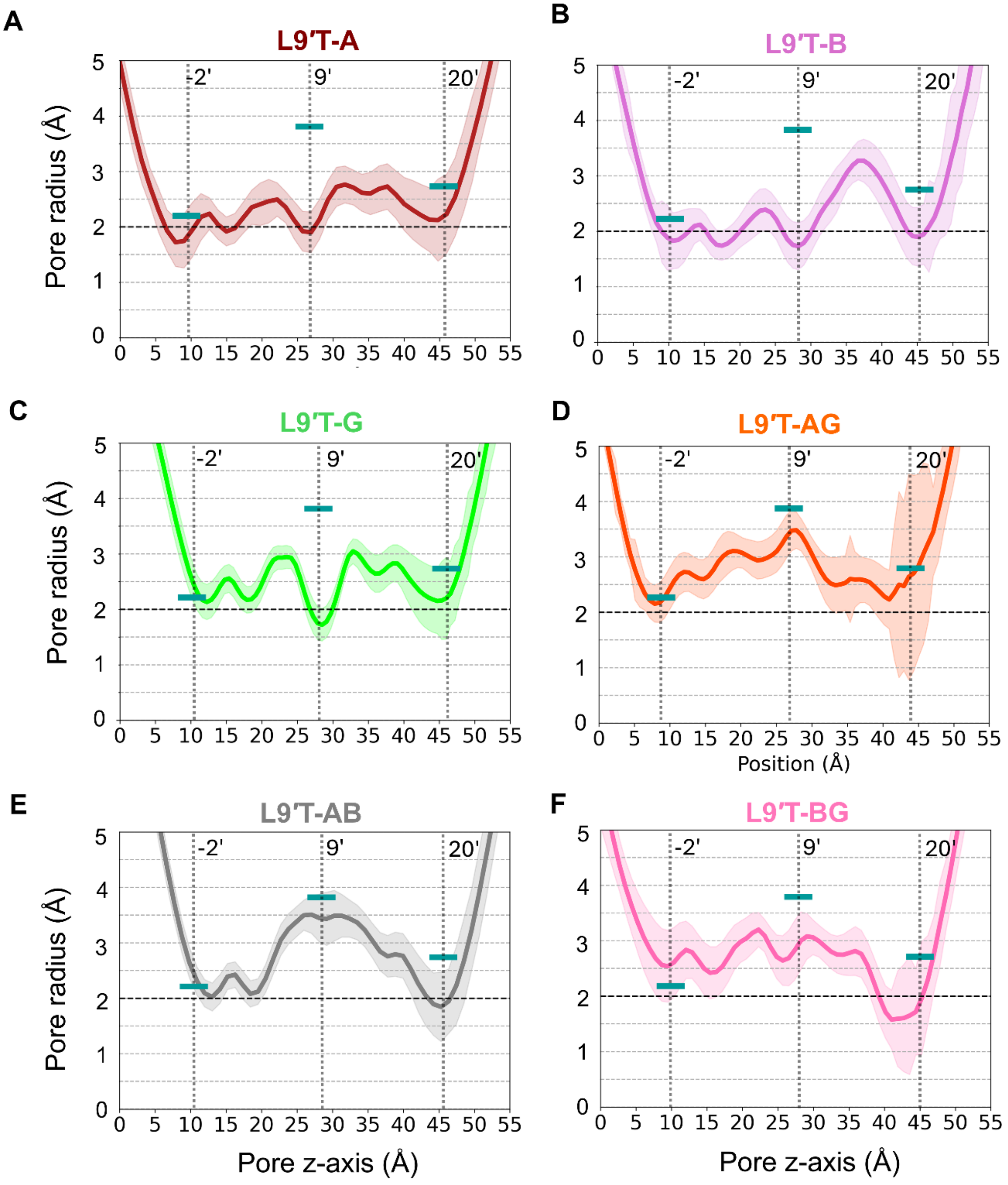


**Figure S12. Pore radius profiles in α1β3γ2 GABA$_A$R single- and double-subunit L9'T mutants.** Time-averaged HOLE radius profiles along the channel axis over 500 ns GaMD. Thick lines (light seagreen) at 20', 9', -2' gates denote all-subunit L9'T means for reference. (A-F) show individual constructs as labeled. Subunit codes: A = α1, B = β3, G = γ2.

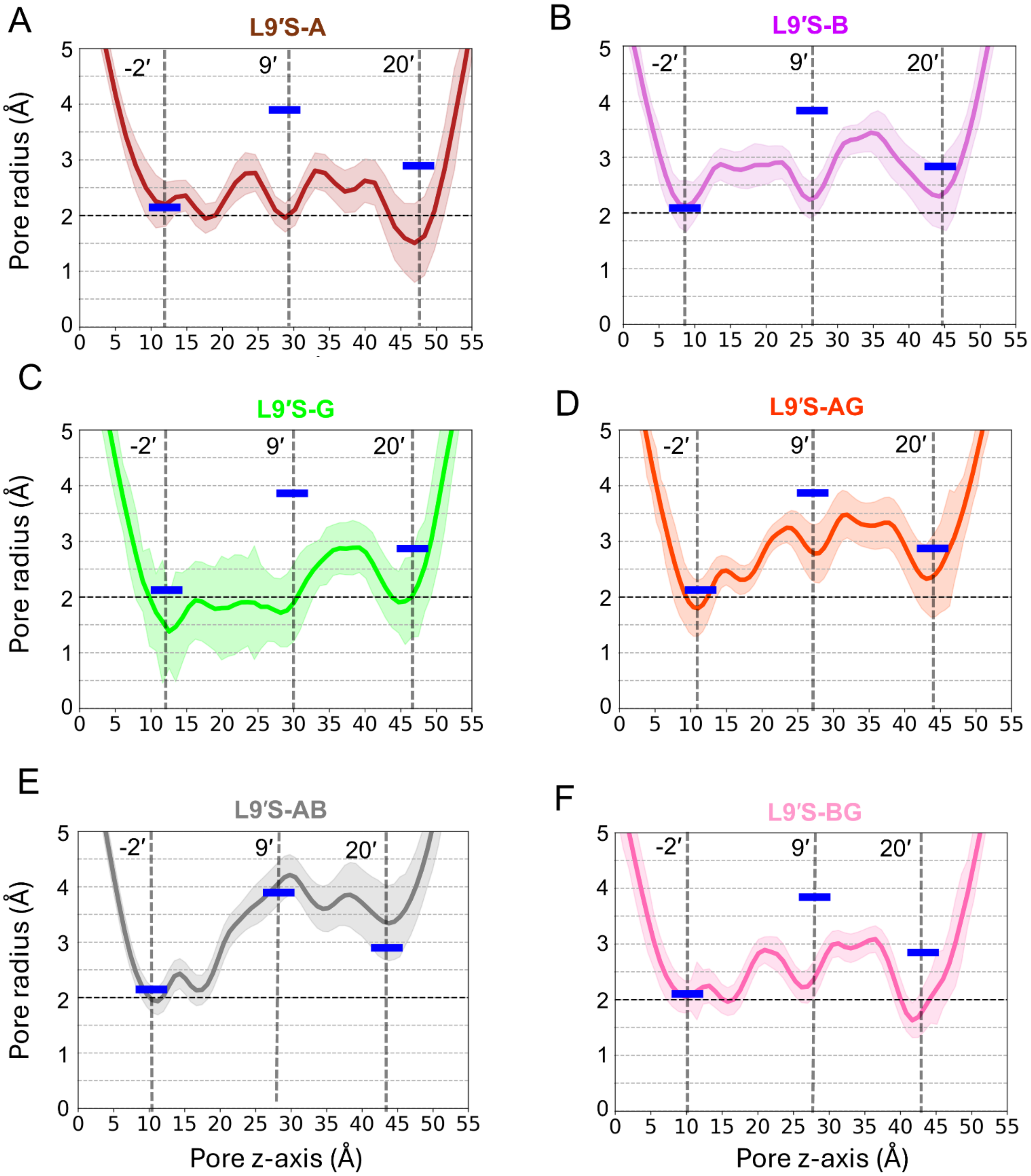


**Figure S13. Pore radius ranges in α1β3γ2 $GABA_AR$ single- and double-subunit L9'S mutants.** Time-averaged HOLE radius profiles along the channel axis over 500 ns GaMD. Thick lines (blue) at 20', 9', -2' gates denote all-subunit L9'S means for reference. (**A-F**) show individual constructs as labeled. Subunit codes: A = α1, B = β3, G = γ2.

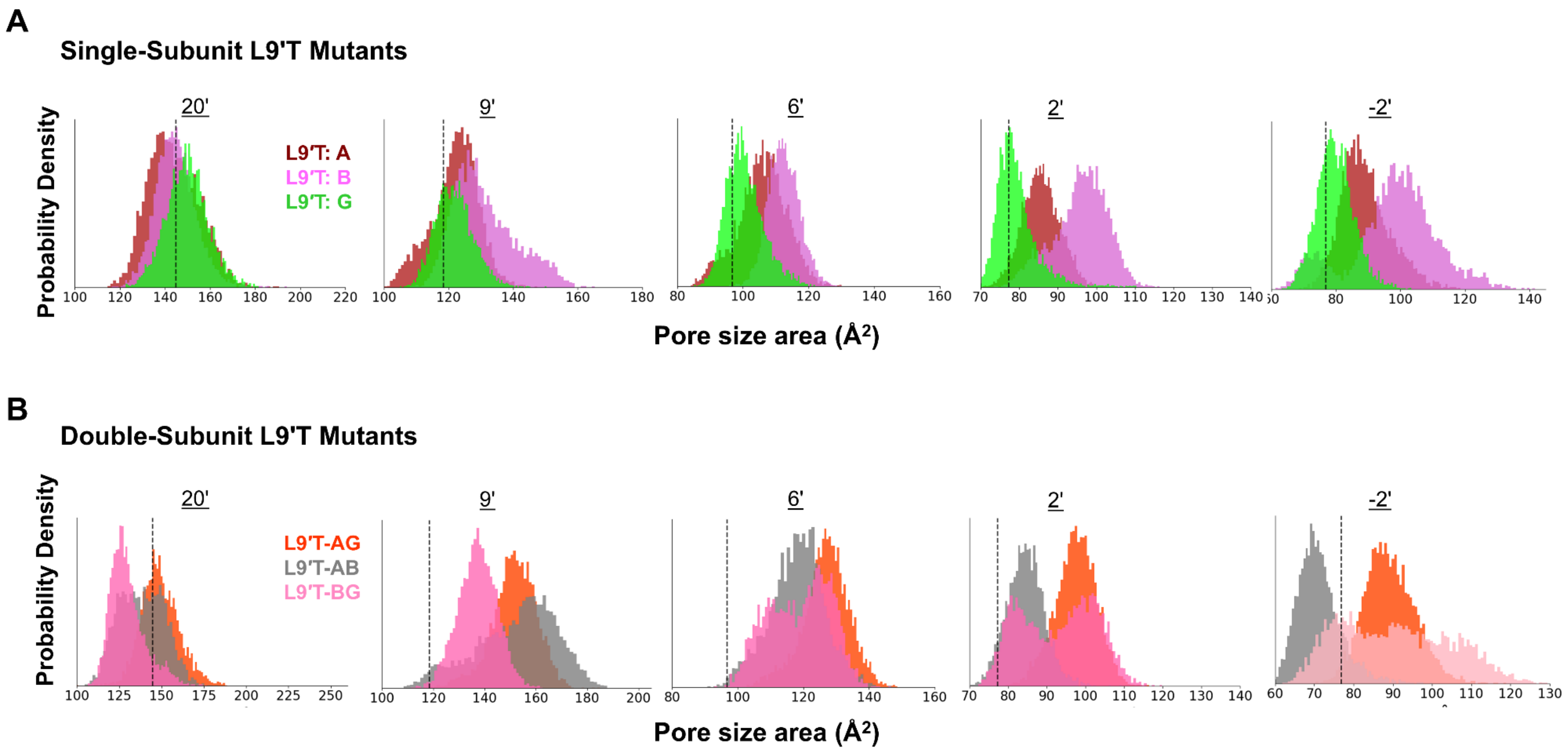


**Figure S14. Cross-sectional pore area distributions for single- and double-subunit L9′T mutants of the α1β3γ2 $GABA_A$ receptor.** Probability density distributions of pore cross-sectional area are shown at five axial positions along the channel pore: 20′, 9′, 6′, 2′, and −2′. **(A)** Single-subunit L9′T mutants: Area distributions for receptors bearing the L9′T substitution in the α (A), β (B), or γ (G) subunit. **(B)** Double-subunit L9′T mutants: Area distributions for L9′T-AG, L9′T-AB, and L9′T-BG combinations. At each axial position, pore area was computed from M2 Cα pentagons. For every simulation frame, the five nearest-neighbor M2 Cα–Cα distances were measured, their mean value was used as the side length of a regular pentagon, and the area of this pentagon was reported (see Methods). This metric captures backbone-level separation among M2 helices and complements HOLE pore-radius measurements. Each density plot represents the full GaMD trajectory of the corresponding mutant. The WT mean is indicated by a black dotted vertical line for reference. Colors are consistent across panels and chosen to distinguish mutant subunit combinations clearly.

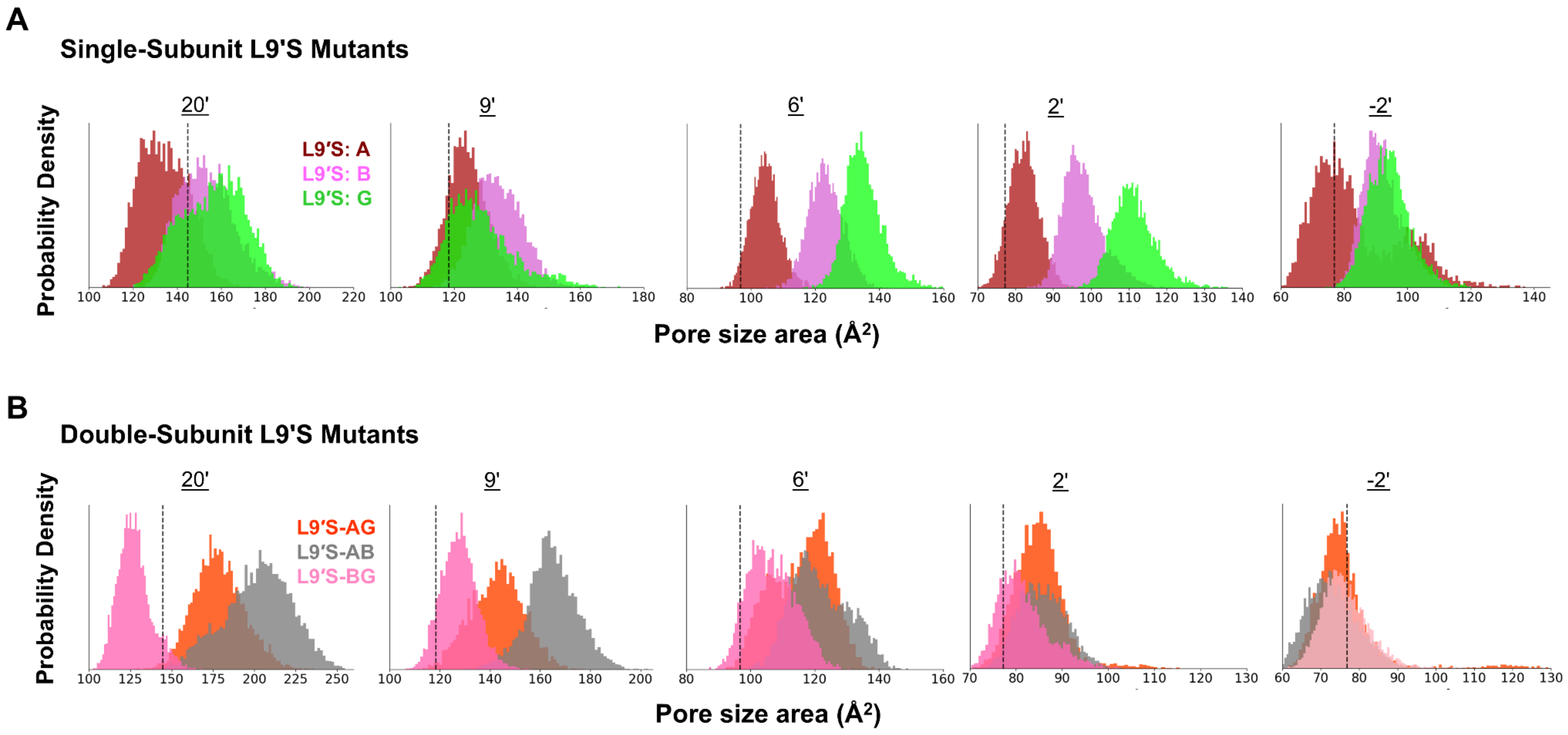


**Figure S15. Cross-sectional pore area distributions for single- and double-subunit L9′S mutants of the α1β3γ2 $GABA_A$ receptor.** Probability density distributions of pore cross-sectional area are shown at five axial positions along the channel pore: 20′, 9′, 6′, 2′, and −2′. **(A)** Single-subunit L9′S mutants: Area distributions for receptors bearing the L9′S substitution in the α (A), β (B), or γ (G) subunit. **(B)** Double-subunit L9′S mutants: Area distributions for L9′S-AG, L9′S-AB, and L9′S-BG combinations. Each density plot represents the full GaMD trajectory of the corresponding mutant. The WT mean is indicated by a black dotted vertical line for reference. Colors are consistent across panels and chosen to distinguish mutant subunit combinations clearly.

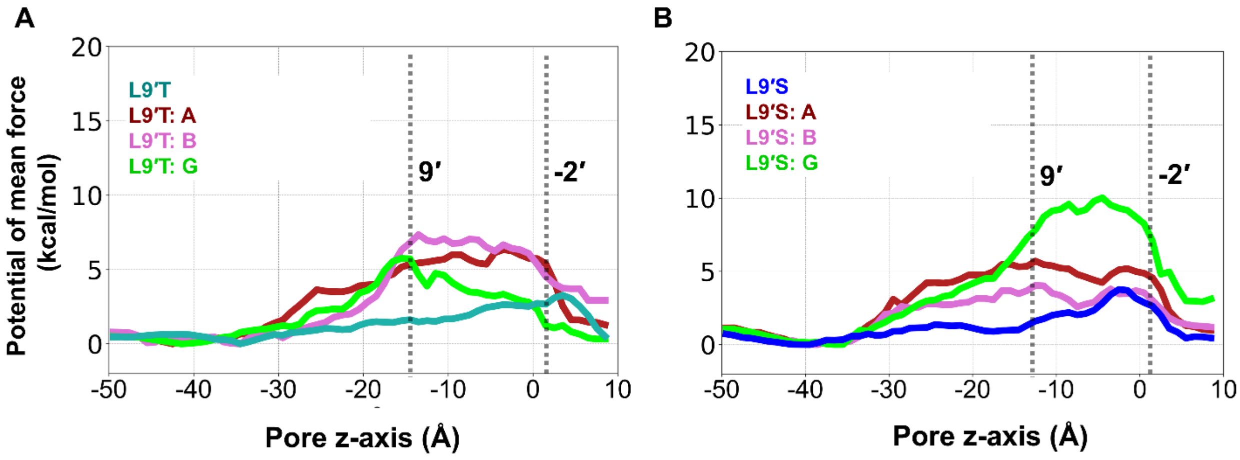


**Figure S16. Free energy profile for $Cl^-$ transport along the pore axis of single-subunit L9'T and L9'S α1β3γ2 $GABA_A$ receptors.** (A) Potential of mean force (PMF) plots show the energetic barrier to $Cl^-$ permeation in all-subunit L9'T (lightseagreen) and single-subunit α1(L264T)β3γ2, α1β3(L259T)γ2, α1β3γ2(L274T) mutant receptors (brown, magenta, and green, respectively). (B) Similar plots are shown for all-subunit (blue) and single-subunit L9'S mutant receptors. Prominent energy barriers at the 9' and -2' positions are indicated by dotted vertical lines. Subunit codes: A = α1, B = β3, G = γ2.

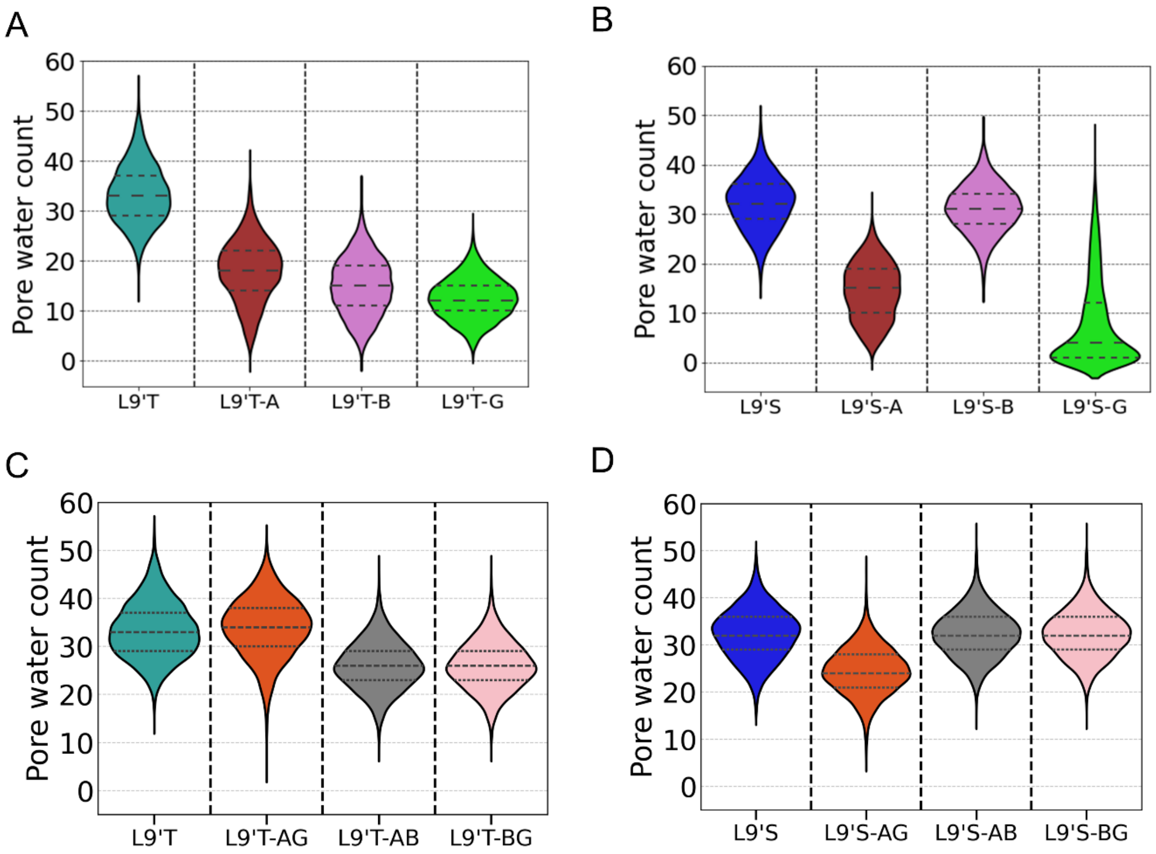


**Figure S17. Hydration near the 9' hydrophobic gate in single- and double-subunit L9'T and L9'S α1β3γ2 GABA$_A$R receptors.** (A-D) Violin plots show the distribution of water molecules within ±7 Å of the 9' gate along the pore axis (7 Å above and 7 Å below) in various single- and double-subunit mutant systems; distributions over 500 ns GaMD trajectories (see the Methods section for details). All-subunit L9'T (light-seagreen) and L9'S (blue) are shown as reference. Subunit codes: A = α1, B = β3, G = γ2.

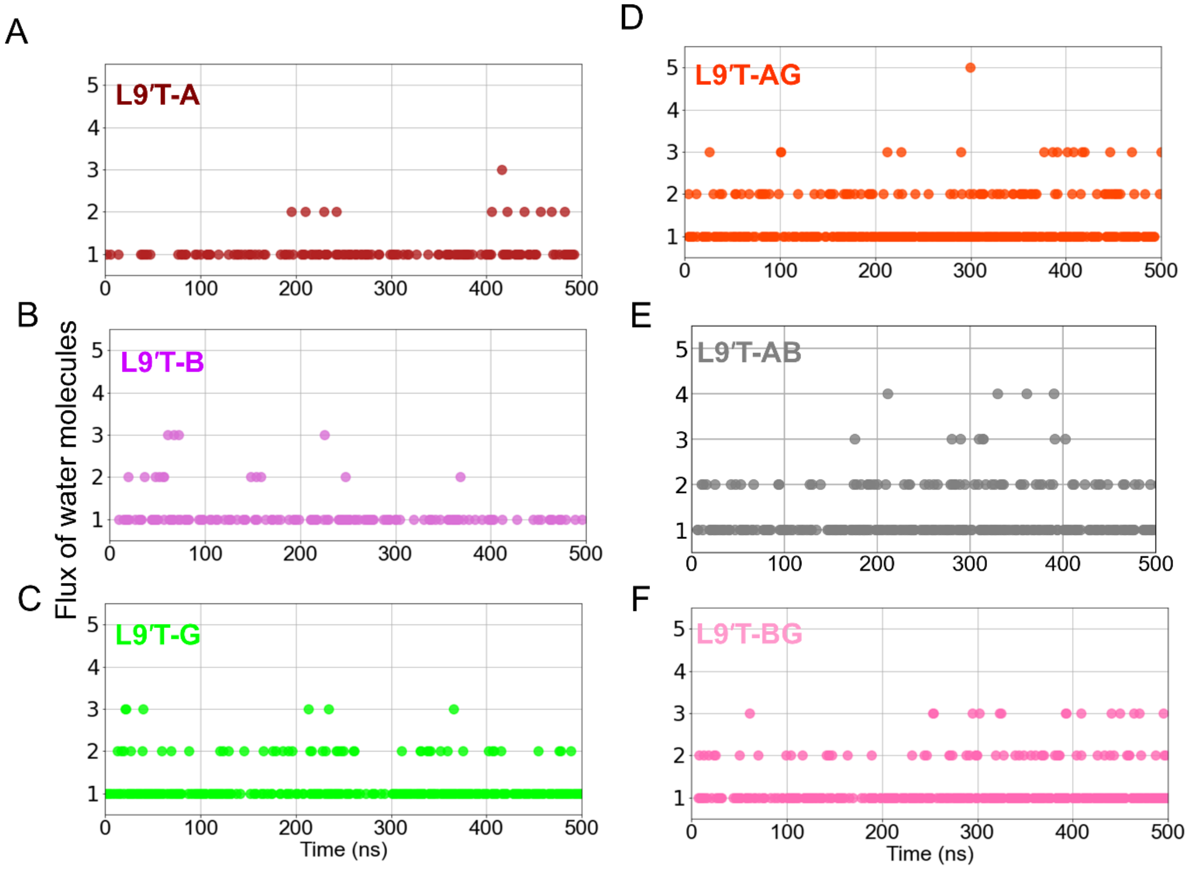


**Figure S18. Water flux time series for single- and double-subunit L9'T α1β3γ2 GABA$_A$ receptors.** Time evolution of water flux (direction-agnostic complete transits between 20' and -2', measured using Wordom) for single-subunit L9'T (A-C) and double-subunit L9'T (D-F) over 500 ns. Single-subunit constructs generally show lower flux than double-subunit constructs. Subunit codes: A = α1, B = β3, G = γ2.

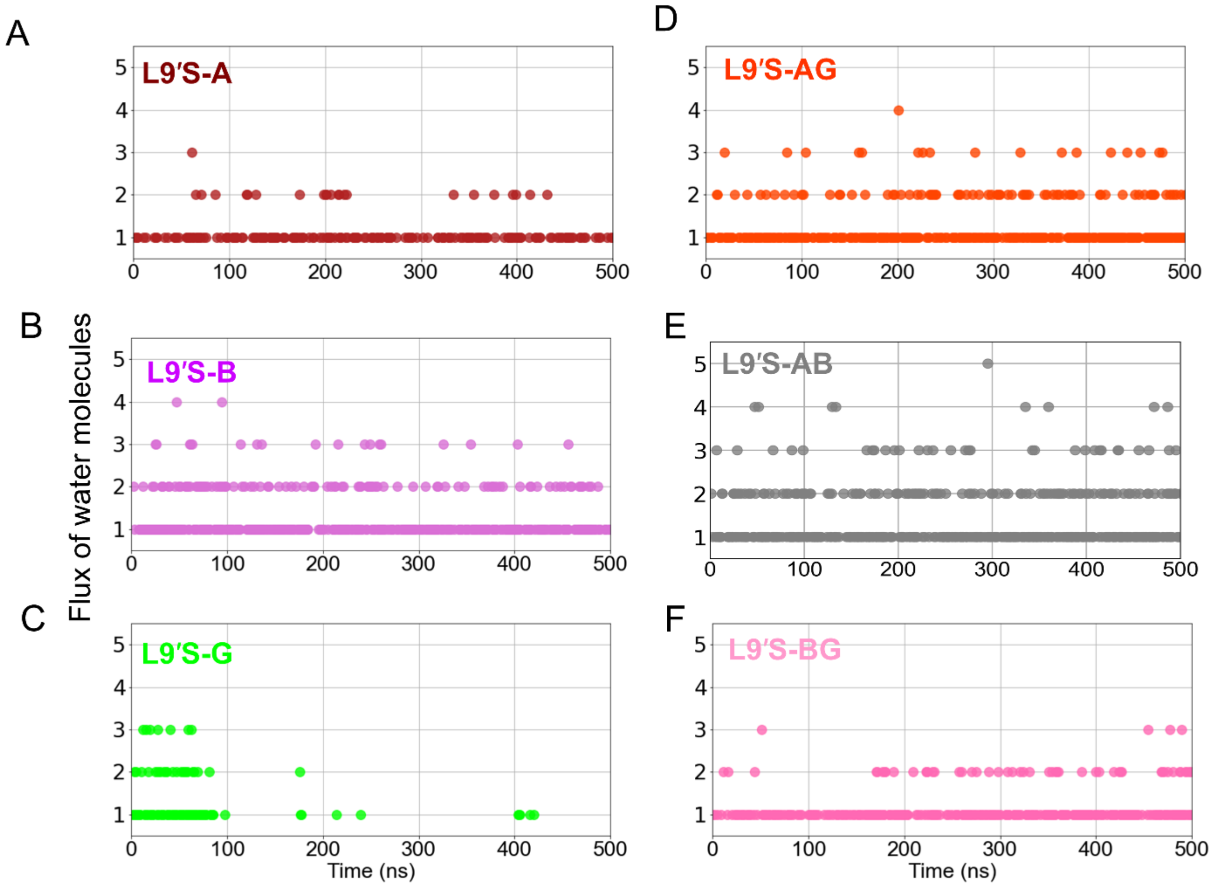


**Figure S19. Water flux time series for single- and double-subunit L9'S α1β3γ2 GABA$_A$ receptors.** Time evolution of water flux (direction-agnostic complete transits between 20' and -2', measured using Wordom) for single-subunit L9'S (A-C) and double-subunit L9'S (D-F) over 500 ns. Single-subunit constructs generally show lower flux than double-subunit constructs. Subunit codes: A = α1, B = β3, G = γ2.

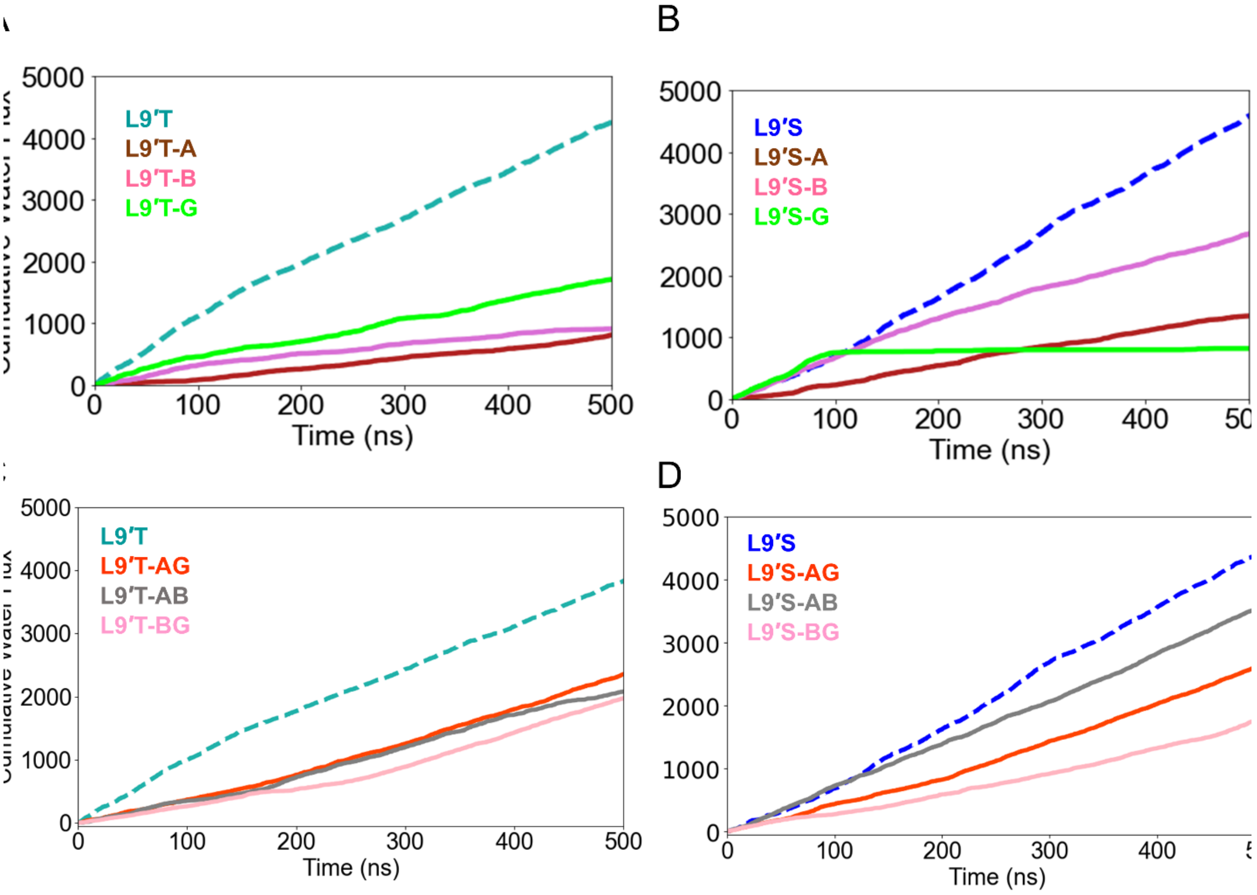


**Figure S20. Cumulative water flux in single- and double-subunit L9'T and L9'S α1β3γ2 GABA$_A$ receptors.** Cumulative count of complete water transits over 500 ns. (A, C) Single- and double-subunit L9'T variants with apo- L9'T traced as a broken light-seagreen reference line. (B, D) Single- and double-subunit L9'S variants with apo-L9'S traced as a broken blue reference line. Subunit codes: A = α1, B = β3, G = γ2.

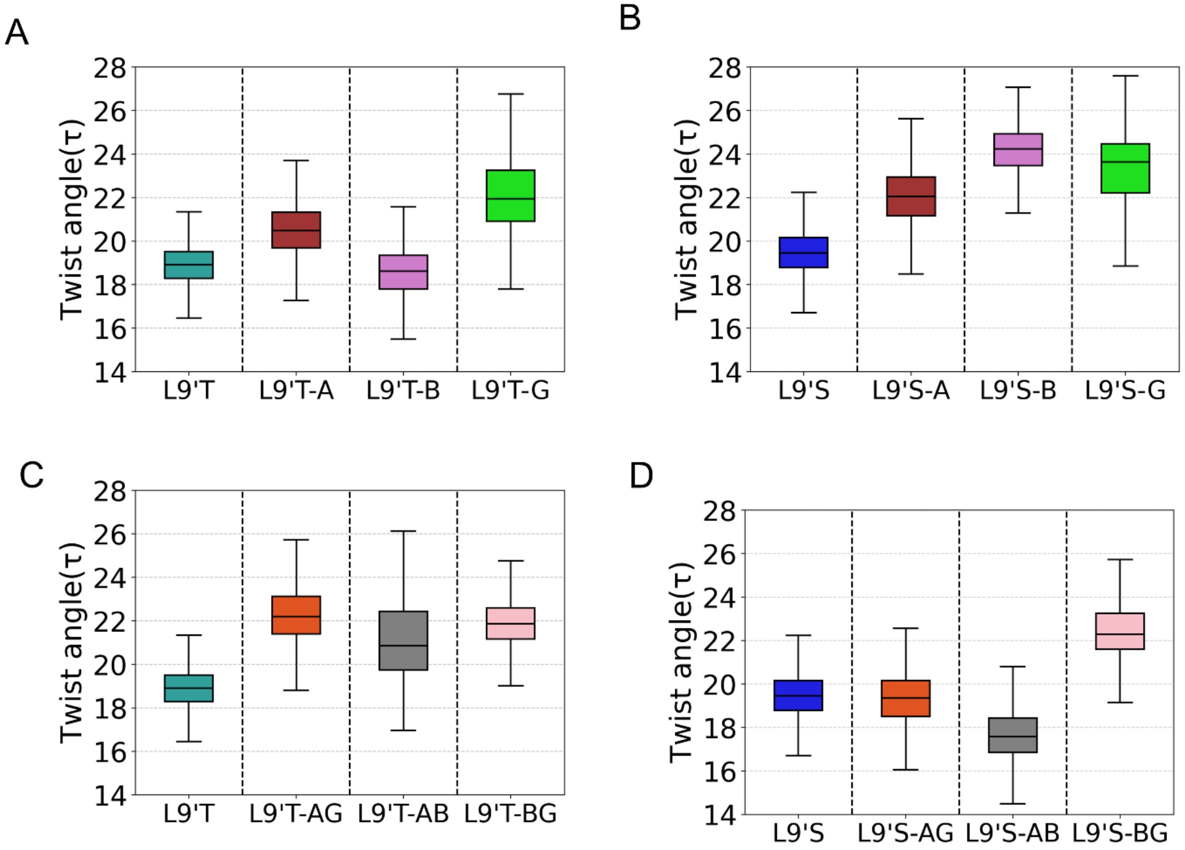


**Figure S21. Global twist profiles for single- and double-subunit L9'T and L9'S α1β3γ2 $GABA_A$ receptors.** Box plots show the twist angle distributions for L9'T and L9'S single -subunit (A, B) and double-subunit (C,D) variants over 500 ns. All-subunit L9'T (light-seagreen) and L9'S (blue) are shown as reference. Lower twist angles align with activation-like untwisting. Angle definitions as in Methods. Subunit codes: A = α1, B = β3, G = γ2.

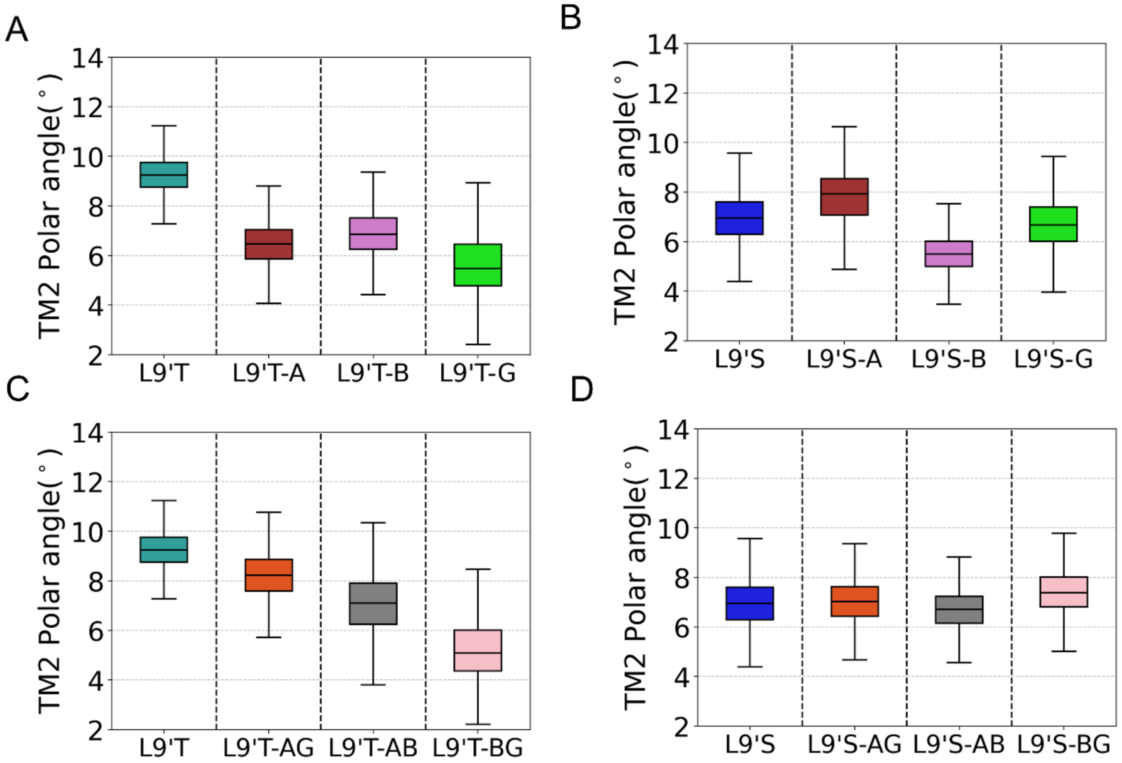


**Figure S22. TMD M2 helix polar tilt profiles for single- and double-subunit L9'T and L9'S α1β3γ2 GABA$_A$ receptors.** Box plots of M2 polar tilt angles for single-subunit L9'T and L9'S (A, B) and double-subunit L9'T and L9'S (C, D) over 500 ns. All-subunit L9'T (light-seagreen) and L9'S (blue) are shown as reference. All single- and double-subunit L9'T variants show lower values than all-subunit L9'T whereas L9'S variants show mixed profiles. Positive polar tilt denotes outward (radial) motion. Angle definitions as in Methods. Subunit codes: A = α1, B = β3, G = γ2.

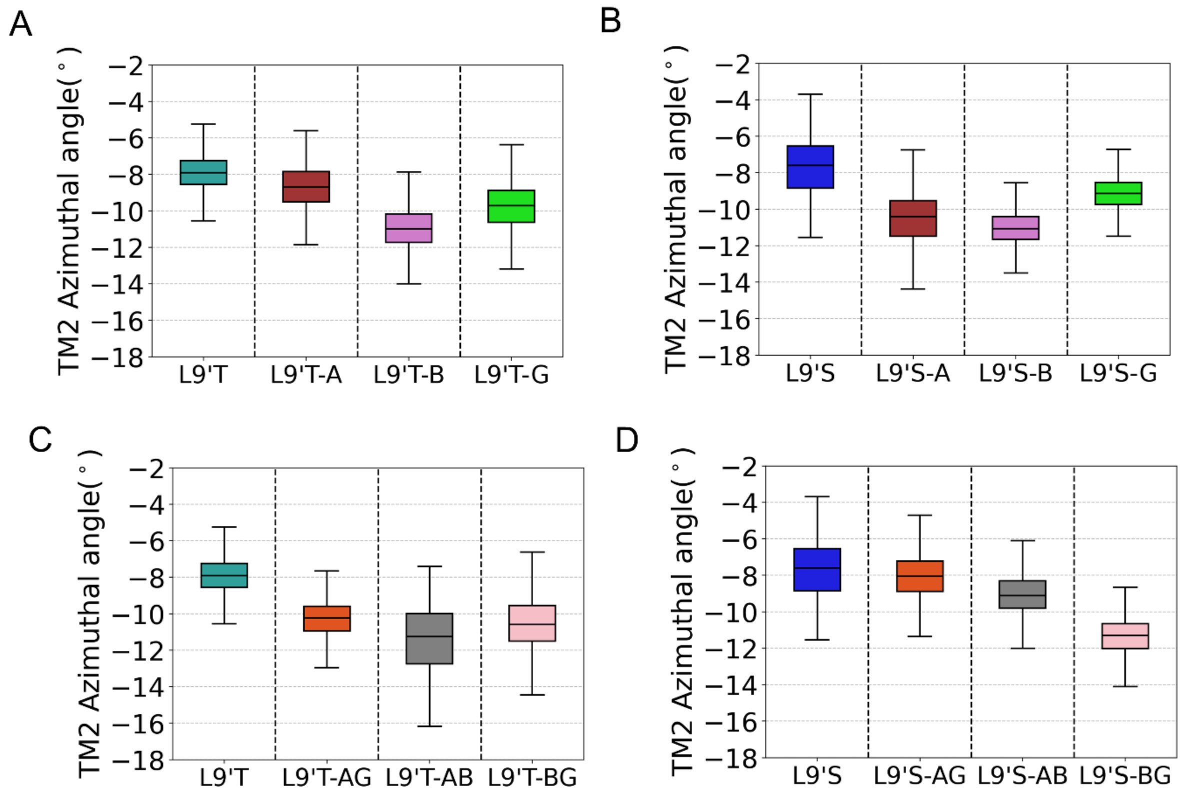


**Figure S23. TMD M2 helix azimuthal tilt profiles for single- and double-subunit L9'T and L9'S α1β3γ2 GABA$_A$ receptors.** Box plots of M2 azimuthal tilt angles for single-subunit L9’T and L9’S (A, B) and double-subunit L9'T and L9'S (C, D) over 500 ns. All-subunit L9'T (light-seagreen) and L9'S (blue) are shown as reference. All single- and double-subunit L9'T and L9'S variants show lower values than their respective all-subunit references. Positive azimuthal tilt denotes clockwise (ECD view) motion. Angle definitions as in Methods. Subunit codes: A = α1, B = β3, G = γ2.

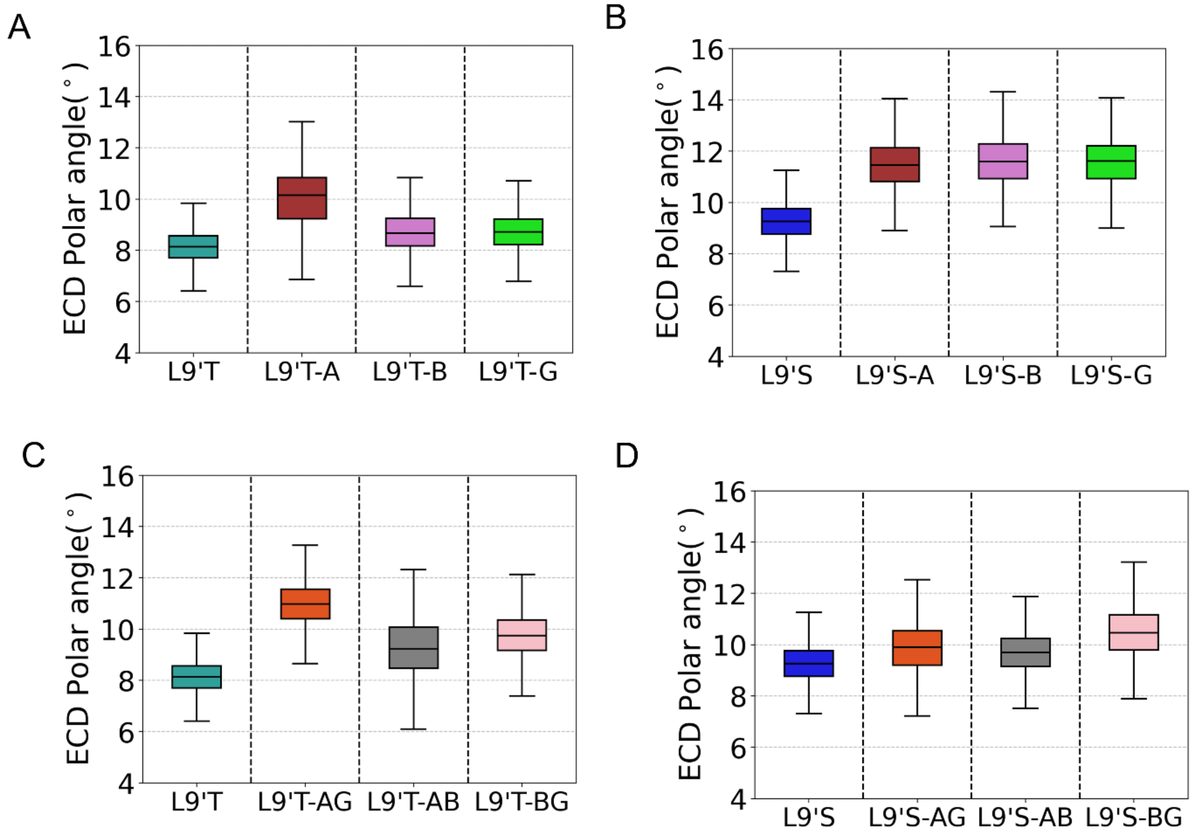


**Figure S24. ECD β-sandwich polar tilt profiles for single- and double-subunit L9'T and L9'S variants.** Polar tilt angle distributions for the ECD β-sandwich in single-subunit L9'T and L9'T (A, B) and double-subunit L9'T and L9'S variants (C, D). All-subunit L9'T (light-seagreen) and L9'S (blue) are shown as reference. Reduced polar tilt indicates ECD compaction. Angle definitions as in Methods. Subunit codes: A = α1, B = β3, G = γ2.

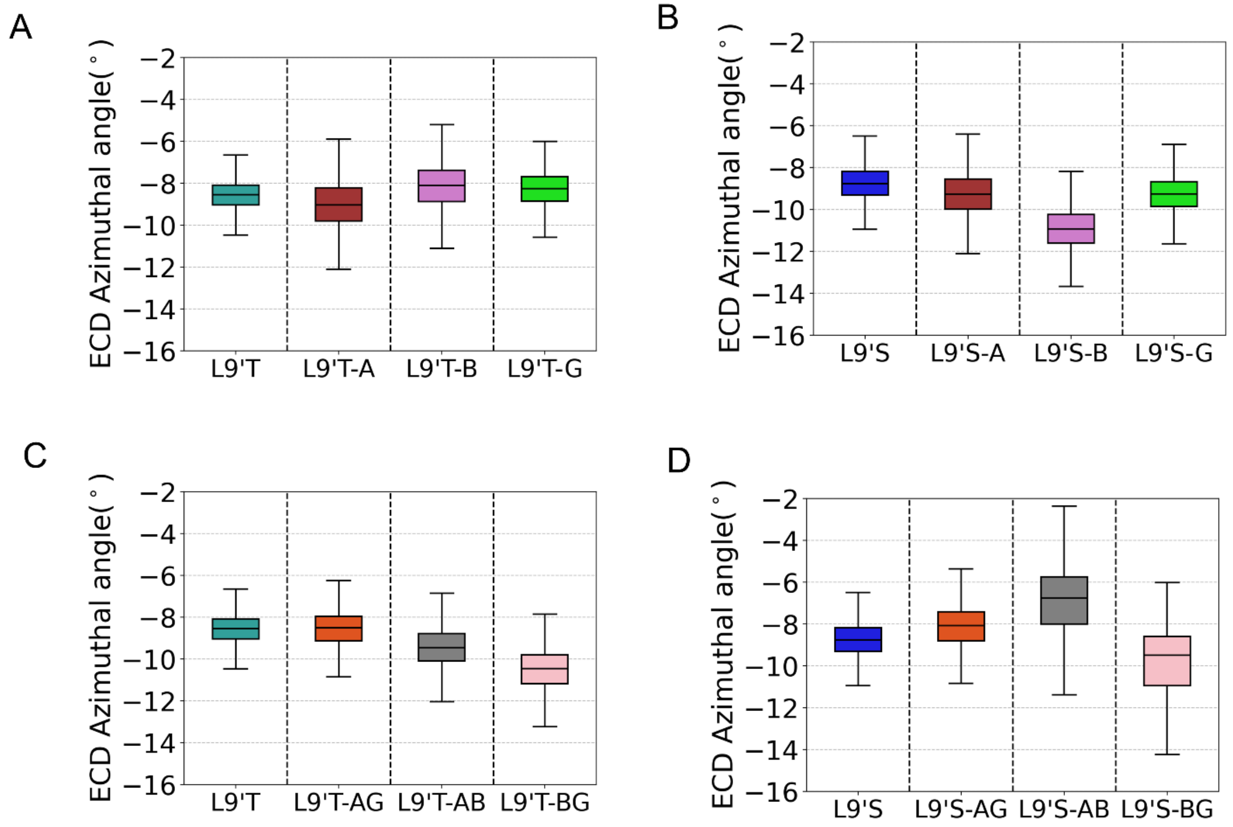


**Figure S25. ECD β-sandwich azimuthal tilt profiles for single- and double-subunit L9'T and L9'S variants.** Azimuthal tilt angle distributions for the ECD β-sandwich in single-subunit L9'T and L9'T (A, B) and double-subunit L9'T and L9'S variants (C, D). All-subunit L9'T (light-seagreen) and L9'S (blue) are shown as reference. Reduced azimuthal tilt indicates ECD compaction. Angle definitions as in Methods. Subunit codes: A = α1, B = β3, G = γ2.

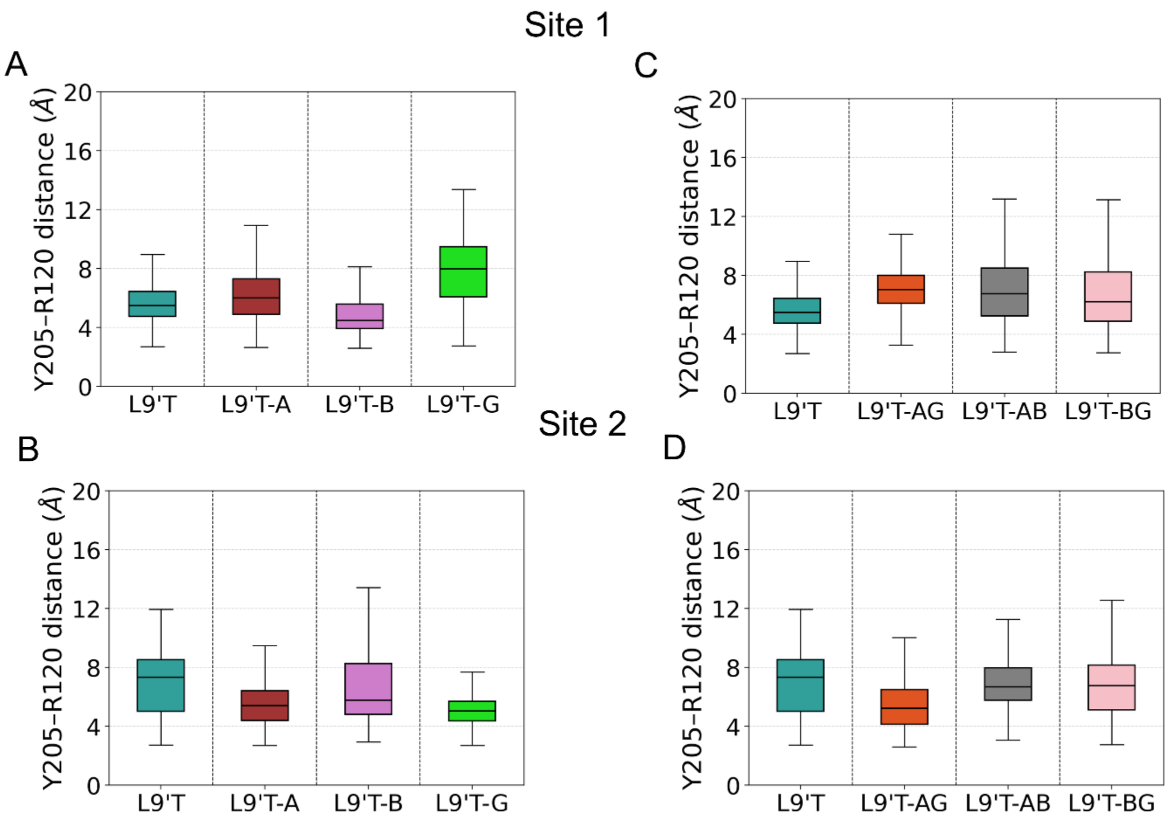


**Figure S26. Orthosteric site C-loop proxy in single- and double-subunit L9'T variants.** β3:Y205-α1:R120 inter-residue distances at orthosteric sites 1 and 2 across L9'T single-subunit variants (A, B) and double-subunit variants (C, D). Atom definitions as in Methods; smaller values indicate C-loop closure. Subunit codes: A = α1, B = β3, G = γ2.

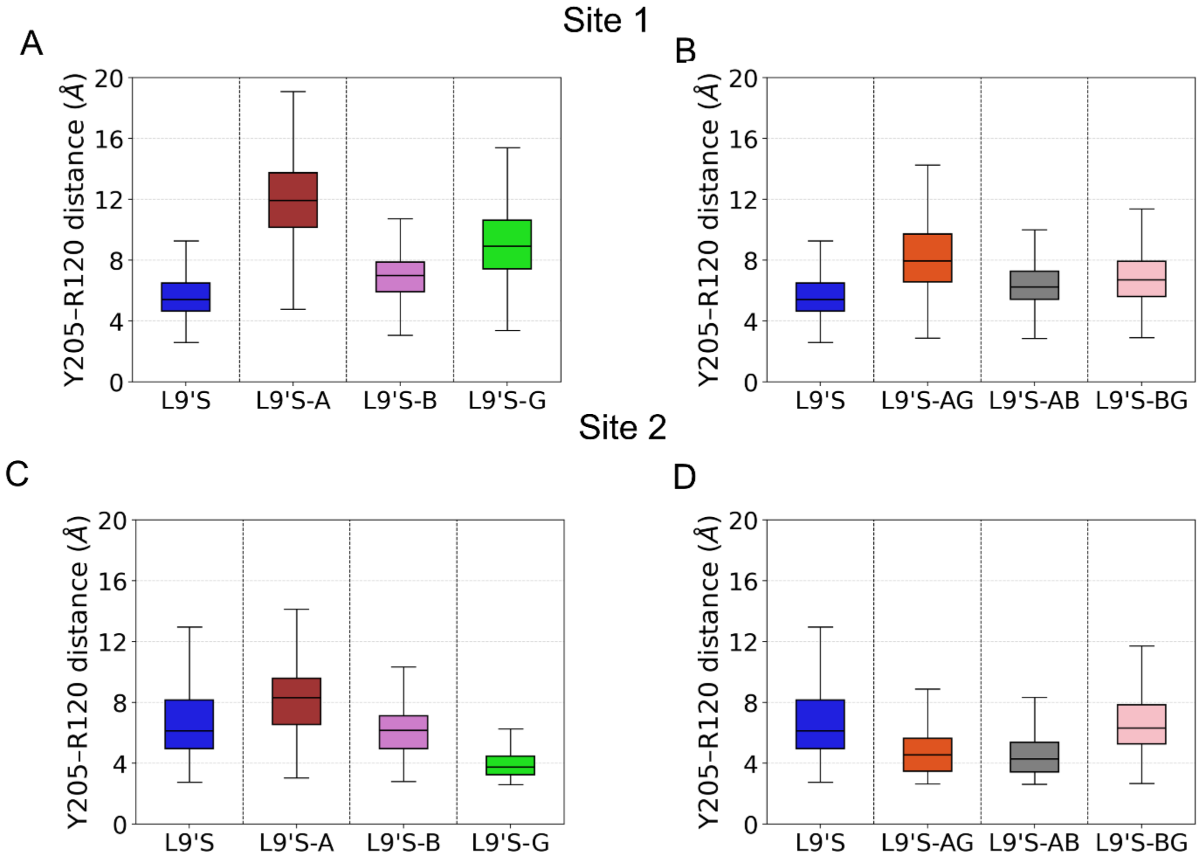


**Figure S27. Orthosteric site C-loop proxy in single- and double subunit L9’S variants.** β3:Y205-α1:R120 inter-residue distances at sites 1 and 2 across L9'S single-subunit variants (A, B) and double-subunit variants (C, D). Atom definitions as in Methods; smaller value indicate C-loop closure. Subunit codes: A = α1, B = β3, G = γ2.
